# Gamblers: an Antibiotic-induced Evolvable Cell Subpopulation Differentiated by Reactive-oxygen-induced General Stress Response

**DOI:** 10.1101/493015

**Authors:** John P Pribis, Libertad García-Villada, Yin Zhai, Ohad Lewin-Epstein, Anthony Wang, Jingjing Liu, Jun Xia, Qian Mei, Devon M. Fitzgerald, Julia Bos, Robert Austin, Christophe Herman, David Bates, Lilach Hadany, P.J. Hastings, Susan M Rosenberg

**Affiliations:** Department of Molecular and Human Genetics; Department of Biochemistry and Molecular Biology; Department of Molecular Virology and Microbiology; The Dan L. Duncan Comprehensive Cancer Center; Graduate Program in Integrative Molecular and Biomedical Sciences Baylor College of Medicine, Houston, TX 77030, USA; Department of Biochemistry and Cell Biology; Systems, Synthetic, and Physical Biology Program Rice University, Houston, TX 77030, USA; Department of Physics; Lewis Sigler Institute Princeton University, Princeton, NJ 08544-0708, USA; Department of Molecular Biology and Ecology of Plants Tel-Aviv University, Tel-Aviv, Israel

**Keywords:** antibiotic resistance, bet hedging, ciprofloxacin, DinB, error-prone DNA polymerases, *Escherichia coli*, evolution, fluoroquinolones, general stress response, mutagenic break repair, reactive oxygen species (ROS), RpoS (σ^S^) stress response, SOS response, starvation stress response, stress-induced mutagenesis, transient differentiation

## Abstract

Antibiotics can induce mutations that cause antibiotic resistance. Yet, despite their importance, mechanisms of antibiotic-promoted mutagenesis remain elusive. We report that the fluoroquinolone antibiotic ciprofloxacin (cipro) induces mutations that cause drug resistance by triggering differentiation of a mutant-generating cell subpopulation, using reactive oxygen species (ROS) to signal the sigma-S (σ^S^) general-stress response. Cipro-generated DNA breaks activate the SOS DNA-damage response and error-prone DNA polymerases in all cells. However, mutagenesis is restricted to a cell subpopulation in which electron transfer and SOS induce ROS, which activate the σ^S^ response, allowing mutagenesis during DNA-break repair. When sorted, this small σ^S^-response-“on” subpopulation produces most antibiotic cross-resistant mutants. An FDA-approved drug prevents σ^S^ induction specifically inhibiting antibiotic-promoted mutagenesis. Furthermore, SOS-inhibited cell division, causing multi-chromosome cells, is required for mutagenesis. The data support a model in which within-cell chromosome cooperation together with development of a “gambler” cell subpopulation promote resistance evolution without risking most cells.

## INTRODUCTION

Antibiotic resistance is a world health threat with 700,000 deaths from resistant infections worldwide annually (O’Neill, 2014). Resistance occurs both by uptake of resistance genes from other bacteria, and de novo mutation of resident genes. Formation of new mutations underpins resistance to diverse antibiotics (Blair et al., 2014; Cannatelli et al., 2014; Palmer et al., 2011), and is a principle resistance route among the World Health Organization’s “priority pathogens” for which new antibiotics are needed (Magrini, 2017). Historically, the challenge of resistance has been met with development of new antibiotics. A complementary approach could be to discover, then inhibit, the molecular mechanisms that drive evolution of resistance (Al Mamun et al., 2012; Cirz et al., 2005; Rosenberg and Queitsch, 2014). Antibiotics not only select resistant mutants but can also induce their formation (Cirz et al., 2005; Gutierrez et al., 2013; Kohanski et al., 2010). Although detailed mechanisms are described by which antibiotics arrest cell growth, how antibiotics induce new mutations is poorly understood.

Fluoroquinolones are widely used antibiotics that inhibit bacterial type-II topoisomerases and kill cells via DNA double-strand breaks (DSBs), by arresting the topoisomerase at DNA double-strand cleavage (Drlica, 1999). Resistance to fluoroquinolones, including ciprofloxacin (cipro), the most used (Hicks et al., 2015), occurs primarily by de novo mutation (Jacoby, 2005). Cipro exposure at so-called “sub-inhibitory” concentrations (below minimal inhibitory concentration, MIC) occurs in ecosystems and during antibiotic therapies, and both induces and selects cipro resistance (Cirz et al., 2005). Another fluoroquinolone, norfloxacin, induced mutations that confer resistance to antibiotics not yet encountered or selected (Kohanski et al., 2010)—antibiotic “cross” resistance. The mutagenesis activity of norfloxacin required reactive oxygen species (ROS) induced by the antibiotic (Kohanski et al., 2010), as does its antibiotic (killing) activity (Kohanski et al., 2007). Yet, how the ROS might lead to mutagenesis—by what molecular mechanism—is unclear. The role of ROS in the antibiotic mechanism is to potentiate lethality by oxidizing DNA bases, the repair of which causes more lethal DNA breaks (Foti et al., 2012; Rasouly and Nudler, 2018; Zhao et al., 2015), but whether this is part of the ROS mutagenic activity is not known.

The general or starvation-stress response of *Escherichia coli*, activated by the sigma-S (σ^S^) transcriptional activator (encoded by the *rpoS* gene), promotes mutagenesis induced by starvation stress (Fitzgerald et al., 2017), and also by beta-lactam (membrane-targeting) antibiotics (Gutierrez et al., 2013). The former occurs by allowing mutagenic repair of spontaneous DSBs, which also requires an SOS DNA-damage response for the upregulation of error-prone DNA polymerase (Pol) IV (Galhardo et al., 2009). The latter occurs by downregulating post-replication error correction (mismatch repair) via a different, SOS-independent mutation mechanism (Gutierrez et al., 2013). The σ^S^ response upregulates Pol IV about two-fold (Layton and Foster, 2003), which might be part of how it promotes mutagenesis during starvation (Fitzgerald et al., 2017).

Bacterial regulatory programs transiently differentiate phenotypically distinct cell subpopulations both stochastically and in response to environmental signals. A potential “bet-hedging” strategy, these subpopulations can allow phenotypes that may be advantageous under stress conditions but deleterious in more permissive environments (Norman et al., 2015; Veening et al., 2008). For example, bacterial “persisters” are a subpopulation of transiently nonproliferating or slowly growing cells, present at about 10^-4^ of the total, that can survive antibiotics without having a resistance mutation, and so lead to persistent infections by resuming growth once antibiotics have gone (Lewis, 2010). Persister formation can occur stochastically, leaving populations ready for a stress that they have not encountered (Balaban et al., 2004), and can also be induced responsively via stress-response regulons including the SOS- (Dorr et al., 2009) and σ^S^-response (Radzikowski et al., 2016) regulons. It is unknown whether antibiotics induce transient differentiation that could promote resistance through mutagenesis, e.g., (Frenoy and Bonhoeffer, 2018).

Here we show that low, sub-inhibitory doses of cipro induce transient differentiation of a small cell subpopulation with high ROS and σ^S^-response activity, that generates mutants, including cross-resistant mutants: a “gambler” subpopulation. We show that the ROS promote mutagenesis in gamblers by activating the σ^S^ response, which allows mutagenic repair of cipro-triggered DSBs—a novel signaling/differentiating role of ROS in mutagenesis. We elaborate the regulatory chain from cipro to ROS to σ^S^ response to mutant production, and also discover a requirement for SOS-induced inhibition of cell division, causing multiple chromosomes per cell. Mathematical analysis supports a model in which multiple chromosomes allow sharing of cellular resources (e.g., recombination, complementation), avoiding deleterious consequences of some mutations during mutagenesis and repair. Thus, multiple chromosomes allow higher mutation rates to be maintained – resulting in faster adaptation. The findings imply a highly regulated, novel transient differentiation process and support a model in which within-cell chromosome cooperation together with development of a gambler subpopulation drive evolution of resistance to new antibiotics without risk to most cells.

## RESULTS

### ROS-dependent Mutagenesis is σ^S^-dependent MBR

We developed two assays to detect mutagenesis induced by cipro independently of cipro selection of the mutants (Figure 1A) (fluctuation tests, **Methods**), and use them to dissect the mechanism of mutagenesis. In both assays, *E. coli* are grown in liquid with low-dose cipro—at the minimum antibiotic concentration (MAC) at which final cfu are 10% of those observed without cipro (Lorian and De Freitas, 1979). The cells are then removed from cipro and plated selectively for colonies with resistance to rifampicin (RifR) or ampicillin (AmpR) antibiotics (Figure 1A), and mutation rates estimated (**Methods**). RifR arises by specific base-substitution mutations in the RNA-polymerase-encoding *rpoB* gene (Reynolds, 2000) (Figure S1A), and AmpR occurs in engineered *E. coli* by *ampD* loss-of-function mutation (Petrosino et al., 2002) (Figures S1B and C, **Methods**). Strikingly, cipro exposure increased apparent RifR and AmpR mutation rates by 26- and 18-fold, above the no-cipro rates, respectively (Figure 1B). The RifR or AmpR mutant cells are not selected in sub-inhibitory cipro, and in fact are at a slight but significant disadvantage (Figure 1C, legend) indicating that mutation rate increases are likely to be underestimates, and that mutation not selection of the mutants is elevated by low-dose cipro. Additional controls show negligible cell death in the low-dose cipro (Figure S1D), obviating potential concerns about death inflating apparent mutation rates (Frenoy and Bonhoeffer, 2018). Other controls for growth rate and colony formation are shown in Figure S2. We conclude that the RifR and AmpR mutations in *rpoB* and *ampD* are induced, and not selected by growth in low-dose cipro.

**Figure 1.**
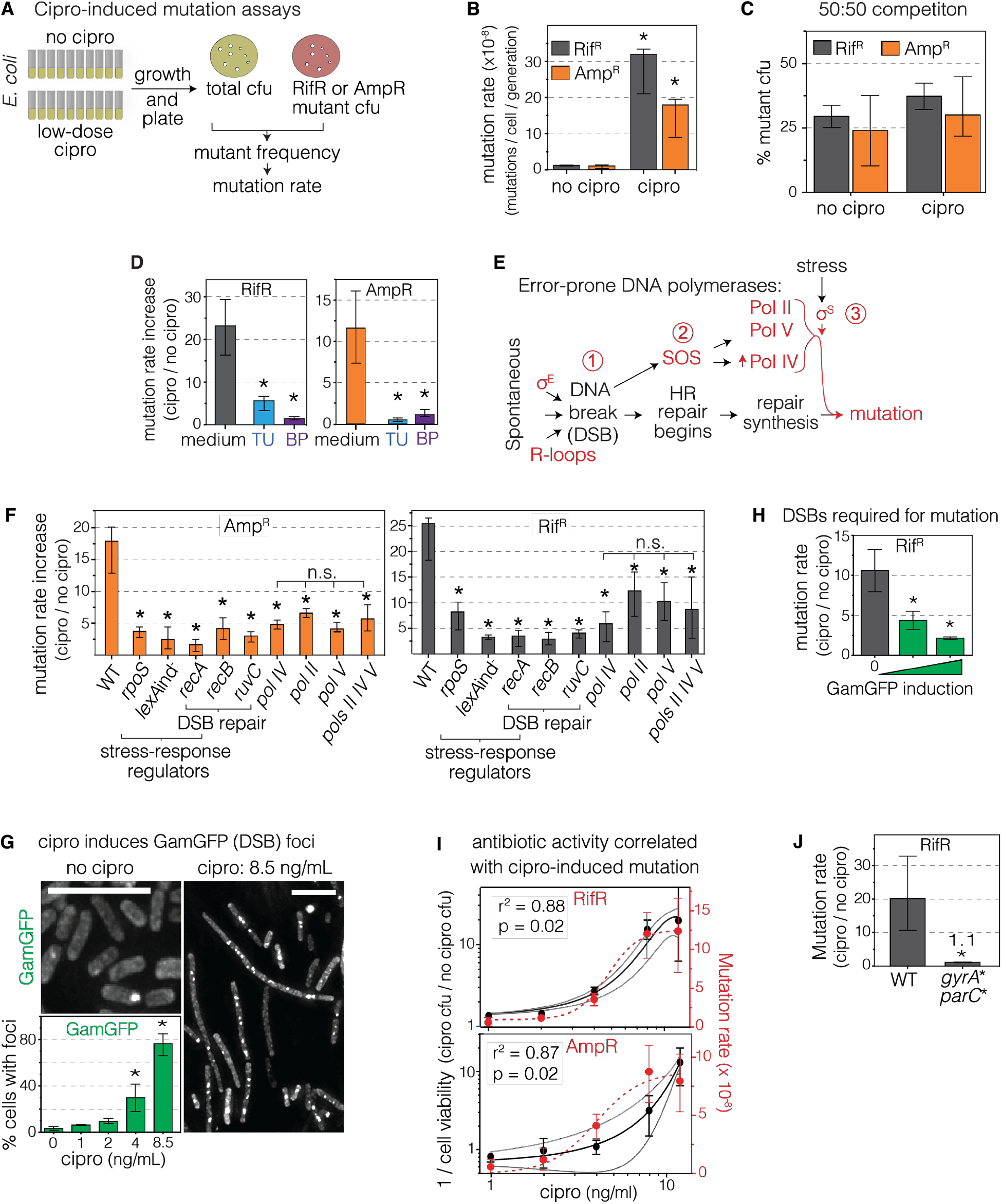
Cipro-Induced Mutagenesis to Antibiotic Cross Resistance via Cipro-Induced ROS and Mutagenic Break Repair. (A) Mutation assays (fluctuation tests). Following growth with/without low-dose cipro, rifampicin- and ampicillin-resistant (RifR, AmpR) mutant cfu are selected (in the absence of cipro) (**Methods**). Mutation rates estimated per **Methods**. (B) Low-dose cipro induces RifR and AmpR mutagenesis. Mutation rates, with and without exposure to sub-inhibitory cipro (8.5 ng/mL). Sequences of RifR base substitutions in *rpoB* gene shown Figure S1A,C; sequences of AmpR null mutations in *ampD*, Figure S1B,C. *Different from no cipro at p<0.001, two-tailed Student’s *t*-test. (C) Competition experiments show that neither RifR *rpoB* mutants nor AmpR *ampD* mutants has a growth advantage in low-dose cipro. Mutant and non-mutant cells mixed at 50% of each, then grown in low-dose cipro per mutation assays. *rpoB* and *ampD* mutants grow less well than their isogenic parents, occupying <50% after growth (AmpR *p* = 0.0098; RifR *p* = 0.0014, 1 sample *t*-test). Thus, the fold-induction of RifR and AmpR mutation rate by cipro (e.g., D) is likely to be an underestimate, and induction of mutagenesis, not selection for RifR or AmpR, underlies increased mutant cfu (B, D, throughout). Means ± SEM of 3 independent experiments. (D) ROS are required for cipro-induced RifR and AmpR mutagenesis. ROS scavenger thiourea (tU) (Wasil et al., 1987), or preventative 2,2’-bipyridine (BP) (Mello Filho et al., 1984) reduces mutation rates (additional controls Figure S2). Data displayed as fold induction of mutation rate. Means ± 95% CI of ≥ 3 independent experiments. *Different from medium at p<0.001, one-way ANOVA with Tukey’s post-hoc test of natural-log transformed data. (E) Starvation-stress-induced mutagenic break repair (MBR), reviewed (Al Mamun et al., 2012; Fitzgerald et al., 2017), requires--(1) a DNA double-strand break (DSB) and its repair by homologous recombination (HR); (2) activation of the SOS response, which transcriptionally upregulates low-fidelity DNA pols IV, V and II; and (3) activation the σ^S^ (RpoS/rpoS) general-stress response, which, by unknown means, licenses use of, or errors made by, low-fidelity DNA polymerases in repair of DSBs. (F) Cipro-induced mutation requires MBR-pathway proteins. Mutants grown at their respective equivalent sub-inhibitory cipro concentrations per **Methods**. Means ± 95% CIs of > 4 independent experiments. *Different from wild-type (WT) at *p*<0.001, one-way ANOVA with Tukey’s post-hoc test of natural-log transformed data; n.s. not significant from each other. Additional double-mutant data (epistasis analyses) Figure S1 E. (G) Cipro induces DSBs dose-dependently. Phage Mu GamGFP labels DSBs as fluorescent foci that are quantified using microscopy, per (Shee et al., 2013). Representative images of GamGFP (DSB) foci in cells grown with or without sub-inhibitory cipro. White scale bar, 10 μm. Mean ± SEM of 3 independent experiments. (H) Reparable DSBs are required for cipro-induced mutagenesis. Production of the DSB-trapping GamGFP protein inhibits DSB repair (Shee et al., 2013) and reduces cipro-induced mutagenesis. Means ± 95% CIs of ≥ 3 independent experiments. *Different from no GamGFP induction at *p*<0.01, one-way ANOVA with Tukey’s post-hoc test of natural-log transformed data. (I) Positive correlation of cipro-induced growth inhibition (antibiotic activity) and mutagenesis implicate cipro-induced DSBs in the mutagenesis. Antibiotic activity, 1/ viable cfu titer, left y axes; mutation rates right y axes. Pearson correlation coefficients in natural-log transformed data, indicate significant correlation of mutation rate with loss of cell viability: RifR r^2^ = 0.87, *p* = 0.02; AmpR r^2^ = 0.88, *p* = 0.02. Means ± 95% CI (right y axes) SD (left y axes) of ≥ 3 independent experiments. (J) Cipro binding to its target type-II topoisomerases is required for induction of mutagenesis, supporting cipro-induced DSBs in the mutagenesis, and obviating potential off-target effects. *gyrA\** S83L/D87Y *parC\** S80I/E84G mutations, which encode functional gyrase and topoisomerase IV proteins that cannot be bound by cipro, prevent mutagenesis. Means ± 95% CIs of 3 independent experiments. *Different from WT at *p*<0.001, two-tailed Student’s *t*-test of natural-log transformed data. See also Figures S1, S2, and S3, and Table S1.

Like mutagenesis induced by norfloxacin (Kohanski et al., 2010), the cipro-induced mutagenesis is ROS dependent, and is inhibited by ROS scavenging/preventing agents thiourea (TU) and 2,2’-bipyridine (BP) (Figure 1D, and below). Whereas the mechanism(s) by which fluoroquinolone-induced ROS promote mutagenesis are unknown, the following data indicate that the ROS instigate a σ^S^-licensed mutagenic DNA break-repair (MBR) mechanism triggered by cipro-induced DSBs.

MBR in starving *E. coli* is regulated mutagenesis during repair of DSBs, requiring the SOS and general (σ^S^) stress responses (Figure 1E), reviewed (Fitzgerald et al., 2017), so that mutation formation, and potentially the ability to evolve, accelerate when cells are maladapted to their environments: when stressed. Spontaneous DSBs induce the SOS DNA-damage response and are repaired by homology-directed DSB repair (HR repair, Figure 1E). The SOS response transcriptionally upregulates error-prone DNA polymerases (Pols) IV, V and II; but repair synthesis does not become mutagenic unless the σ^S^ response is also induced (Ponder et al., 2005; Shee et al., 2011) by starvation (Al Mamun et al., 2012) (Figure 1E). The σ^S^ response, by unknown means, allows use of, or persistence of errors made by, Pols IV, V and II in DSB repair causing mutations (Frisch et al., 2010; Ponder et al., 2005; Shee et al., 2011) near DSBs (Shee et al., 2012).

We find that most cipro-induced *ampD* and *rpoB* mutagenesis requires proteins used in starvation-stress-induced MBR (Figure 1F): DSB-repair proteins RecA, RecB, and RuvC, the SOS and general-stress-response activators, and SOS-upregulated error-prone DNA Pols IV, V, and II, implying a MBR-like mutagenesis mechanism. Isogenic strains grown at their appropriate MACs (Table S1) showed 87% ± 3% and 70% ± 9% decreases in cipro-induction of mutagenesis (AmpR and RifR combined) when carrying an SOS non-inducible *lexAInd* mutation or Δ*rpoS* (σ^S^) deletion, respectively (mean ± 95% CI, Figure 1F). Thus both stress responses are required for the mutagenesis. Moreover, double-defective mutant cells show no further reduction (Figure S1E), indicating their action in the same mutagenesis pathway. Further, ROS and the σ^S^ response also act in the same mutagenesis pathway, in that scavenging ROS with thiourea (TU) caused no further reduction of mutagenesis to Δ*rpoS* cells (Figure S1F, and additional controls Figures S1 D and S2). We conclude that cipro-induced ROS-dependent mutagenesis occurs by the σ^S^-dependent MBR pathway.

Moreover, the mutagenesis also requires DSBs and their repair. We visualized and quantified cipro-induced DSBs as fluorescent foci of the GamGFP DSB-end-specific binding protein (Shee et al., 2013), under our low-dose exposure conditions (8.5 ng/ml) and found that cipro increased GamGFP (DSB) foci by 28 ± 9 times above the spontaneous DSB level (mean ± SEM Figures 1G, Figure S3A,B, additional controls Figure S4A). We also show that GamGFP protein, which binds DSB ends and prevents their repair (Shee et al., 2013), inhibited cipro-induction of mutagenesis (Figure 1H), indicating a requirement for reparable DSBs in mutagenesis. Additionally, RecBCD, interacts specifically with DSB ends in DSB repair (Kuzminov, 1999), and its requirement in cipro-induced RifR and AmpR mutagenesis (Figure 1F, *recB*) also implies the necessity of DSBs in the mutagenesis. The data indicate that DSBs and DSB repair are necessary for mutation formation, and support a MBR mutagenesis mechanism.

The following data implicate specifically cipro-induced DSBs in the MBR mutagenesis. First, cipro antibiotic activity results from DSBs caused by inhibition of *E. coli* type II topoisomerases gyrase and topo IV mid-reaction (Drlica, 1999). In dose-response experiments, we find a tight correlation between cipro antibiotic and mutagenic activities (Figures 1I and S1G, r^2^ = 0.87, 0.88, Pearson correlation), correlating the DSB-induction (antibiotic) activity of cipro with its role in mutagenesis. Second, we used special *gyrA\** and *parC\** mutants that produce functional gyrase- and topo IV-mutant proteins that are not bound by cipro (Khodursky et al., 1995), and find no induction of mutagenesis with cipro (Figure 1J). These data show that cipro action on its target topoisomerases is needed for induction of mutagenesis, eliminate possible “off-target” effects of cipro on mutagenesis, and indicate a role for the cipro-induced DSBs in mutagenesis. Finally, a^E^ and R-loop-promoting proteins, which promote starvation-stress-induced MBR by promoting spontaneous DSBs (Gibson et al., 2010; Wimberly et al., 2013) (Figure 1E) are not required for cipro-induced MBR (Figure S1H), supporting an MBR mechanism like that in starvation, except with the DSBs resulting from cipro inhibition of topoisomerases, rather than spontaneous sources. The data indicate that DSBs generated via cipro trigger the MBR pathway.

Together, the data support a σ^S^-dependent MBR mechanism instigated by cipro-induced ROS and DSBs, allowing “MBR” to fill the previous mechanism void between ROS and mutations. The role of ROS might be contributing to the DNA breakage, as ROS do in antibiotic-killing mechanisms (Foti et al., 2012; Rasouly and Nudler, 2018; Zhao et al., 2015), action after stress-response induction, as in starvation-stress-induced MBR (Moore et al., 2017), or at another stage. The data below show a novel role for ROS in mutagenesis as signaling molecules that activate the general stress response, and, surprisingly, that this is limited to a transiently differentiated cell subpopulation.

### ROS Differentiate a Cell Subpopulation, Activate σ^S^ Response

We surveyed cipro-treated single cells for induction of ROS, and the SOS and σ^S^ stress responses using flow cytometry. We measured SOS induction using an SOS-response-reporter gene, P*_sulA_mCherry*, engineered at a non-genic chromosomal site (Nehring et al., 2015; Pennington and Rosenberg, 2007), and found that cipro promotes SOS dose-dependently and uniformly among cells (Figure 2A), with 208 ± 26 times more SOS-positive cells at the 8.5ng/mL mutagenic dose than without cipro (mean ± SEM Figure 2A). Auto-fluorescence, which is induced by bactericidal antibiotics (Renggli et al., 2013), is negligible compared with the SOS (or the ROS or σ^S^)-activity fluorescence signals (Figures S4B-D).

**Figure 2.**
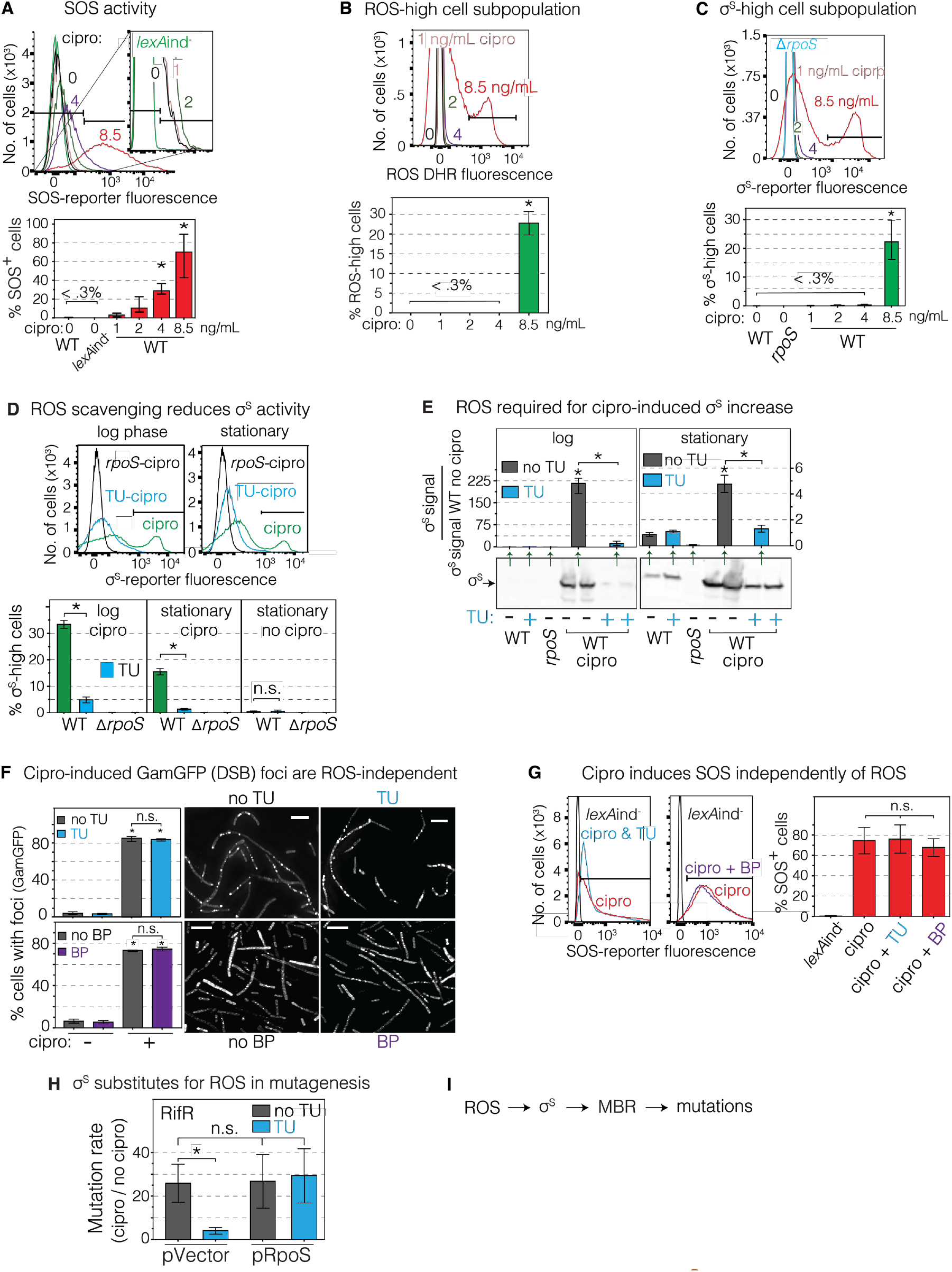
ROS Form in Minority Cell Subpopulation, Activate σ^S^ Response and MBR. (A) Population-wide dose-dependent activation of the SOS response by cipro. Flow-cytometry assay of log-phase cells with chromosomal SOS reporter *PsulAmCherry* quantifies single cells with DNA damage that triggers the SOS response. SOS-positive cells are those right of the gate illustrated (black bars, **Methods**). Fluorescence, arbitrary fluorescence units (afu). Means ± SEM of 3 independent experiments. *Different from no cipro at *p*<0.01, one-way ANOVA with Tukey’s post-hoc test. (B) Low-dose cipro induces high ROS in a minority cell subpopulation: 25% ± 6% of cells in log phase. Flow cytometry assay of log-phase cells showing green fluorescence of ROS-detecting dye dihydrorhodamine 123 (DHR). ROS-high cells scored as those within the gate illustrated (black horizonal bar). Means ± SEM of 3 independent experiments. *Different from no cipro at *p*<0.01, one-way ANOVA with Tukey’s post-hoc test (C) Low-dose cipro induces high σ^S^-general-stress-response activity in a minority cell subpopulation: 22% ± 3% of cells in log phase. Flow cytometry assay of log-phase cells showing σ^S^-response activity as fluorescence from σ^S^-response reporter *yiaG-yfp*. σ^S^-high cells are those right of the gate shown (black bar, Methods). Stationary-phase cells at time of assay for mutants have smaller σ^S^-high subpopulation of ~10% of cells (F below, Figures 3 and 4). Means ± range of 2 independent experiments. *Different from no cipro at *p*<0.01, one-way ANOVA with Tukey’s post-hoc test. (D) ROS are required for cipro induction of the σ^S^ response. ROS scavenger TU reduces σ^S^ activity, removing the σ^S^ high-activity cell subpopulation. Flow cytometric assays per (C). Means ± SEM of 3 independent experiments. \**p*<0.01, one-way ANOVA with Tukey’s post-hoc test. (E) ROS are required for cipro induction of σ^S^-protein levels. Scavenging ROS with TU inhibits σ^S^-protein accumulation in log and stationary phase. Quantification and representative σ^S^ western blot. Means ± range of 2 independent experiments. *Different from no cipro at *p*<0.01, one-way ANOVA with Tukey’s post-hoc test. (F) Cipro induces DSBs independently of ROS. Reduction of ROS with TU or BP does not alter levels of GamGFP (DSB) foci in cells grown with or without cipro. Representative images of GamGFP (DSB) foci, per (Shee et al., 2013). White scale bar, 5 μm. Means ± SEM of ≥ 3 independent experiments. *Different from no cipro at *p*<0.001, one-way ANOVA with Tukey’s post-hoc test; n.s. not significant. (G) SOS induction is independent of ROS. ROS reducers TU and BP do not inhibit SOS-response activation. SOS activity measured by flow cytometry in strains with the chromosomal SOS reporter *PsulAmCherry*. Means ± SEM of ≥ 3 independent experiments. n.s. not significant, oneway ANOVA with Tukey’s post-hoc test. (H) Engineered production σ^S^ substitutes for ROS in cipro-induced mutagenesis, allowing mutagenesis in TU-treated cells. The data imply that the sole or major role of ROS in cipro-induced mutagenesis is activation of the σ^S^ response, making ROS unnecessary when σ^S^ is produced artificially. ROS and σ^S^ also work in the same pathway (are epistatic, Figure S1F). Means ± range of 2 independent experiments. \**p* <0.01, one-way ANOVA with Tukey’s post-hoc test; n.s., not significant. (I) Summary: cipro-induced ROS induce the σ^S^ response, which allows mutagenic break repair (MBR) and generation of mutations. Not shown: the ROS and σ^S^ response occur in minority cells subpopulation(s). See also Figures S1, S2, S3, and S4, and Tables S1 and S2.

Surprisingly, low-dose cipro induced both ROS and the σ^S^ general stress response strongly in only a discreet cell subpopulation(s). In flow-cytometric assays of ROS in log-phase cipro-treated cells using the peroxide (H_2_O_2_)-sensitive dye dihydrorhodamine 123 (DHR, Figure 2B), high ROS levels appeared in only in a distinct 25% ± 6% cell subpopulation (8.5ng/mL, Figure 2B, mean ± SEM). Similarly, measuring σ^S^ activity with the *yiaG-yfp* fluorescence reporter (Al Mamun et al., 2012) in log-phase cipro-treated cells also revealed high σ^S^ activity in a discreet subpopulation of 22% ± 3% of the cells (Figures 2C and S3B, additional controls Figure S3C). For both ROS and the σ^S^ response, induction occurred above a threshold with low ROS or σ^S^ activity at doses below the 8.5ng/mL dose at which mutagenesis is assayed (Figure 2B, C). Growth inhibition, a known ROS-dependent process (Kohanski et al., 2007), also occurred above an 8.5ng/ml threshold (Figure S3D). A threshold is also seen in the kinetics of induction of translation of the *rpoS* mRNA (to σ^S^ protein) by three small RNAs (sRNAs) (Soper et al., 2010), examined below. Thus, despite uniform/unimodal, dose-dependent induction of DBSs (Figure 1G) and SOS (Figure 2A), fluoroquinolone induction of high-ROS and the general stress response occurs in only a well separated subpopulation of ~20% of exposed log-phase cells, and these have very high ROS and σ^S^-activity levels, respectively (Figure 2B, C). The subpopulation is smaller, near 10%, when the cells reach stationary phase (Figure 2D), when mutagenesis is assayed.

We found that ROS are required for, and promote mutagenesis by, activation of the σ^S^ response. First, ROS scavenging/preventing agents TU or BP blocked induction of σ^S^-response activity removing the σ^S^-high cell subpopulation (Figures 2D and S3E), reduced the accumulation of σ^S^-protein (western blotting, Figure 2E), or of a σ^S^-β-galactosidase fusion protein (Figure S3F) from an *rpoS-lacZ* reporter, indicating that ROS are required for induction of the σ^S^ response. Additional controls (Figure S5A and B) show that ROS are not generally needed for fluorescence, transcription, or protein accumulation, just for activation of the σ^S^ response by cipro. We conclude that σ^S^-response induction by cipro requires ROS.

Furthermore, ROS promote cipro-induced MBR not by creation of DSBs, but instead, by induction of the σ^S^ response in a transiently differentiated cell subpopulation. The role of ROS in antibiotic (growth-inhibitory) activity (Table S1) is creation of DNA breaks via oxidized guanine (8-oxo-dG) in DNA (Dwyer et al., 2007; Foti et al., 2012; Kohanski et al., 2007). By contrast, we observed that reduction of cellular ROS levels with TU or BP, though profoundly inhibitory to the MBR mutagenesis (Figure 1D), did not reduce induction of DSBs by low-dose cipro, quantified as GamGFP foci (same TU and BP concentrations as mutation assays, Figure 2F and S3G). Neither did TU or BP diminish the SOS DNA-damage response (Figure 2G), implying that ROS promote mutagenesis independently of damage to DNA. Moreover, 8-oxo-dG incorporation appears not to be the principal role of ROS in the mutagenesis in that ROS-mediated 8-oxo-dG-signature mutations [G·C➝T·A and A·T➝C·G, (Schaaper and Dunn, 1987)] are a *less* important part of cipro-induced than spontaneous forward-mutations (*ampD*, Figure S1C). These data indicate that ROS promote mutagenesis other than by promoting DSBs or SOS-responsive DNA damage, or base misincorporation opposite oxidized guanine during DNA replication.

Further, the main or only role of ROS in cipro-induced MBR mutagenesis is σ^S^ induction, in that artificial/engineered upregulation of σ^S^ fully substituted for ROS in mutagenesis (Figures 2H and S5C). RifR mutagenesis was restored in the presence of TU by IPTG induction of σ^S^ (Figures 2H). ROS and the σ^S^ response also act in the same mutagenesis pathway (Figure S1E). The data indicate that ROS are needed for MBR only or mostly for induction of σ^S^, such that if σ^S^ is otherwise supplied, ROS are no longer required for the mutagenesis (Figure 2H,I). Importantly, cipro induction of SOS, ROS, and σ^S^ activities all require cipro interaction with its topoisomerase targets, in that cells with active but cipro-non-binding mutant *gyrA\** and *parC\** alleles (Khodursky et al., 1995) showed no induction of SOS, ROS, or σ^S^ responses by cipro (Figure S3H-J, respectively). These data demonstrate that the events that lead to SOS, ROS, and σ^S^ induction begin with cipro interaction with its topoisomerase targets.

Together, these data show that cipro action on topoisomerases leads to induction of high ROS levels in a discreet cell subpopulation (Figure 2B), that the ROS activate σ^S^ in a subpopulation (Figure 2C-E), and that activation of the σ^S^ response is how ROS promote cipro-induced MBR (Figure 2H, I). This constitutes a novel role for ROS in mutagenesis—signaling induction of the σ^S^ general stress response—unlike those in antibiotic activity (Dwyer et al., 2007; Foti et al., 2012; Kohanski et al., 2007) or starvation-stress-induced MBR (Moore et al., 2017).

### σ^S^-active Gambler Cell Subpopulation Generates Mutants

We used fluorescence-activated cells sorting (FACS) to demonstrate that the small σ^S^-response-high cell subpopulation, which encompasses 13% (± 1%) of the stationary-phase cells used in mutagenesis assays, produces most cipro-induced mutants (Figure 3). We sorted σ^S^ high- and low-activity cells to at least 97% enrichment (Figures S6A-C). Remarkably, whereas unsorted and mock-sorted cells show (mean) 25 ± 3-fold induction of RifR mutant frequencies by cipro (Figure 3A), the sorted σ^S^-high cells displayed a 400 ± 7-fold induction of RifR mutagenesis—16 ± 3-times higher than unsorted and mock-sorted cells (Figure 3A, controls for the sorted populations Figures 3B, S6D, E, and S7A, B, **Supplemental Discussion S1**). The large σ^S^-low-activity cell subpopulation, 87% ± 1% of cells, showed only 3 ± 1-fold induction of RifR mutagenesis by cipro, or 8 ± 2-times lower than unsorted and mock-sorted cells (Figure 3A), indicating that most mutants did not arise in the majority cell subpopulation. We can estimate the contribution of each subpopulation to total yield of mutants as follows. Because the σ^S^ low-activity cells display only a 3 ±1-fold increase in RifR mutants (Figure 3A), we can conclude that the σ^S^-low cells produced about 12% of the mutants (3-fold increase / 25-fold increase in un/mock-sorted = 12%, Figure 3A). Because the σ^S^-low cells will be contaminated with some σ^S^-high cells, this means that *at least* 88% of RifR mutant yield originates in the σ^S^ high-activity cell subpopulation. The data demonstrate that most of the cipro-induced RifR mutants originate in the small σ^S^-high cell subpopulation.

**Figure 3.**
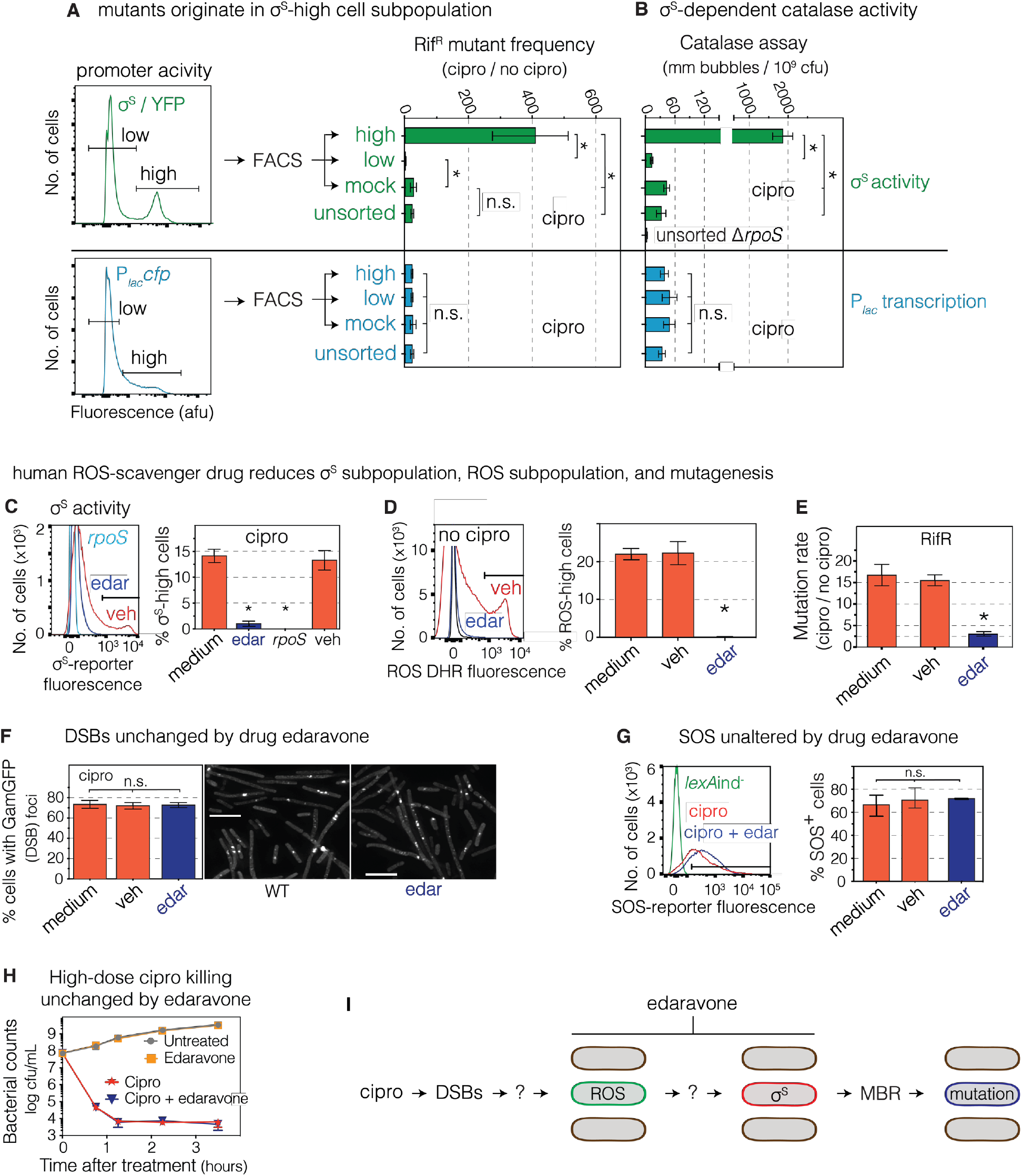
σ^S^-response-high Gambler Cell Subpopulation Generates Mutants, Is Inhibited by FDA-approved Drug. (A) Most cross-resistant mutants are produced by the minority σ^S^ high-activity cell subpopulation. Cells with high and low fluorescence from σ^S^-response or *lac* reporters were sorted by FACS and assayed for mutants. Induction of mutants by cipro: 400 ± 7-fold in minority σ^S^-high cells (13±1% of cells); 25 ± 3-fold in unsorted and mock-sorted cells; 3 ± 1-fold in the majority σ^S^-low cells (87±1% of cells). These equate to ≥88% of cipro-induced mutants arising in 13% of cells (text). Mutant frequencies, means ± 95% CI of 3 independent experiments. \**p*<0.01, one-way ANOVA with Tukey’s post-hoc test; n.s., not significant. (B) High σ^S^-dependent catalase activity in σ^S^ high-activity cells, confirms their σ^S^-high status. HPII is the σ^S^-regulated catalase. Means ± SEM of 3 independent experiments. \**p*<0.01, one-way ANOVA with Tukey’s post-hoc test; n.s., not significant. (C) FDA-approved antioxidant drug edaravone inhibits appearance of the σ^S^-response-activated cell subpopulation. Flow-cytometry of stationary-phase cells with the *yiaG-yfp* σ^S^-response reporter. Representative flow cytometry histograms of cells grown with cipro, with or without edaravone. Means ± range of 2 independent experiments. *Different from medium at *p*<0.001, one-way ANOVA with Tukey’s post-hoc test. (D) FDA-approved antioxidant drug edaravone inhibits appearance of the ROS-high cell subpopulation required for σ^S^-response induction (Figure 2D,E,H). Flow-cytometry of stationary-phase cells with DHR123 ROS dye. Different from medium at *p*<0.001, one-way ANOVA with Tukey’s post-hoc test. (E) FDA-approved antioxidant drug edaravone inhibits cipro-induced mutagenesis. Means ± range of 2 independent experiments. *Different from no-drug samples at *p*<0.001, one-way ANOVA with Tukey’s post-hoc test (F, G) Edaravone does not affect (F) cipro induction of DSBs, quantified as GamGFP foci per (Shee et al., 2013), or (G) the SOS response, measured by flow cytometry in cells carrying the chromosomal SOS reporter *PsulAmCherry*. Means ± range of 2 independent experiments. n.s., not significant from WT, one-way ANOVA with Tukey’s post-hoc test. (H) FDA-approved antioxidant drug edaravone does not reduce high-dose cipro antibiotic killing activity. Log-phase cells grown with or without high-dose cipro (1.5μg/mL) with or without edaravone, and cfu/mL determined. Means ± range of 2 independent experiments. (I) Summary: cipro induces high ROS levels in a minority cell subpopulation. The σ^S^ high-activity cell subpopulation generates most resistant mutants: a gambler cell subpopulation. FDA-approved anti-oxidant drug edaravone inhibits mutagenesis by reducing ROS and appearance of the σ^S^-high gambler cell subpopulation. Whether the ROS-high cells are the same cells as the σ^S^-high cell subpopulation is addressed Figure 4. The source of ROS induced by cipro, shown with ? to the left of the ROS-high cell, and how ROS promote σ^S^ activation, shown with ? to the right of the ROS-high cell, are unknown and are addressed below. Ovals, *E. coli* cells. See also Figures S2, S5, and S6, and Tables S1 and S2.

We excluded the possibility that the greater production of mutants by σ^S^-high cells might result indirectly from their high fluorescence (possible high metabolic activity), using a fluorescence-reporter gene not controlled by σ^S^: P*_lac_cfp* (cyan fluorescent protein, Figure 3A, above autofluorescence, Figure S4E). Additional controls for sorted cells’ growth rates / colony formation are shown in Figures S2A,B.

Further, we find that the hypermutability of the σ^S^-high cell subpopulation that generates RifR mutants appears to be transient, and not a heritable mutator state, as, for example, a “mutator” mutation would confer. RifR mutants recovered are not heritably mutator (Figure S5D).

Collectively, the data demonstrate that a small, transiently differentiated subpopulation of σ^S^ high-activity cells is transiently hypermutable and generates most cipro-induced mutants with de novo rifampicin-(cross)-resistance mutations. These data suggest a potential “bet-hedging” developmental strategy (Norman et al., 2015; Veening et al., 2008) that may allow evolution while risking mutagenesis in only some cells; we call these gambler cells. How the gambler cell subpopulation is differentiated, and a drug that prevents it, follow.

### FDA-approved Drug Inhibits Evolvability

The σ^S^-response high-activity cell subpopulation may be considered a novel therapeutic target for potential inhibition of cipro-induced mutagenesis to antibiotic cross resistance. We show that the drug edaravone, an ROS scavenger indicated for human use in ALS in the U.S. and cerebral infarction in Japan (Miyaji et al., 2015; Watanabe et al., 2018), inhibits cipro induction of mutagenesis but not its antibiotic (killing) activity. Edaravone, at concentrations seen in new formulations (100μM) (Corporation, 2014; Li et al., 2012; Parikh et al., 2016), inhibited the appearance of σ^S^-high cells (Figure 3C), accumulation of σ^S^-fusion protein (Figure S3F), appearance of ROS-high cells (Figure 3D), and most (82% ± 1% of) RifR mutagenesis (Figure 3E). Edaravone did not affect cipro induction of DSBs (Figure 3F), SOS activation (Figure 3G), cell growth (Figure S2A), colony formation (Figures S2B), or negative-control β-gal activity (Figure S5B), implying that its inhibition of mutagenesis reflects specific inhibition of σ^S^-response activation (Figure 3I). Importantly, edaravone did not alter the ability of high-dose cipro to kill *E. coli* (Figure 3H), showing that edaravone can reduce mutagenesis induced by cipro without altering its utility as an antibiotic. These data serve as a proof-of-concept for small-molecule inhibitors that could be administered with antibiotics to reduce resistance evolution, by impeding differentiation of σ^S^-response-active gambler cells, without harming antibiotic activity. We explored the basis of differentiation of the σ^S^ response-high cell subpopulation—how ROS activate the σ^S^ response in the subpopulation cells (Figure 3I)—as follows.

### ROS-high Cells Become Gamblers via sRNAs

The ROS-high subpopulation cells could, in principle, induce σ^S^ activity in other cells, or themselves, or both. We distinguished these possibilities by following single cells over time through induction of ROS then the σ^S^ response, using fluorescence reporters, flow cytometry and time-lapse microscopy after cipro (Figure 4).

**Figure 4.**
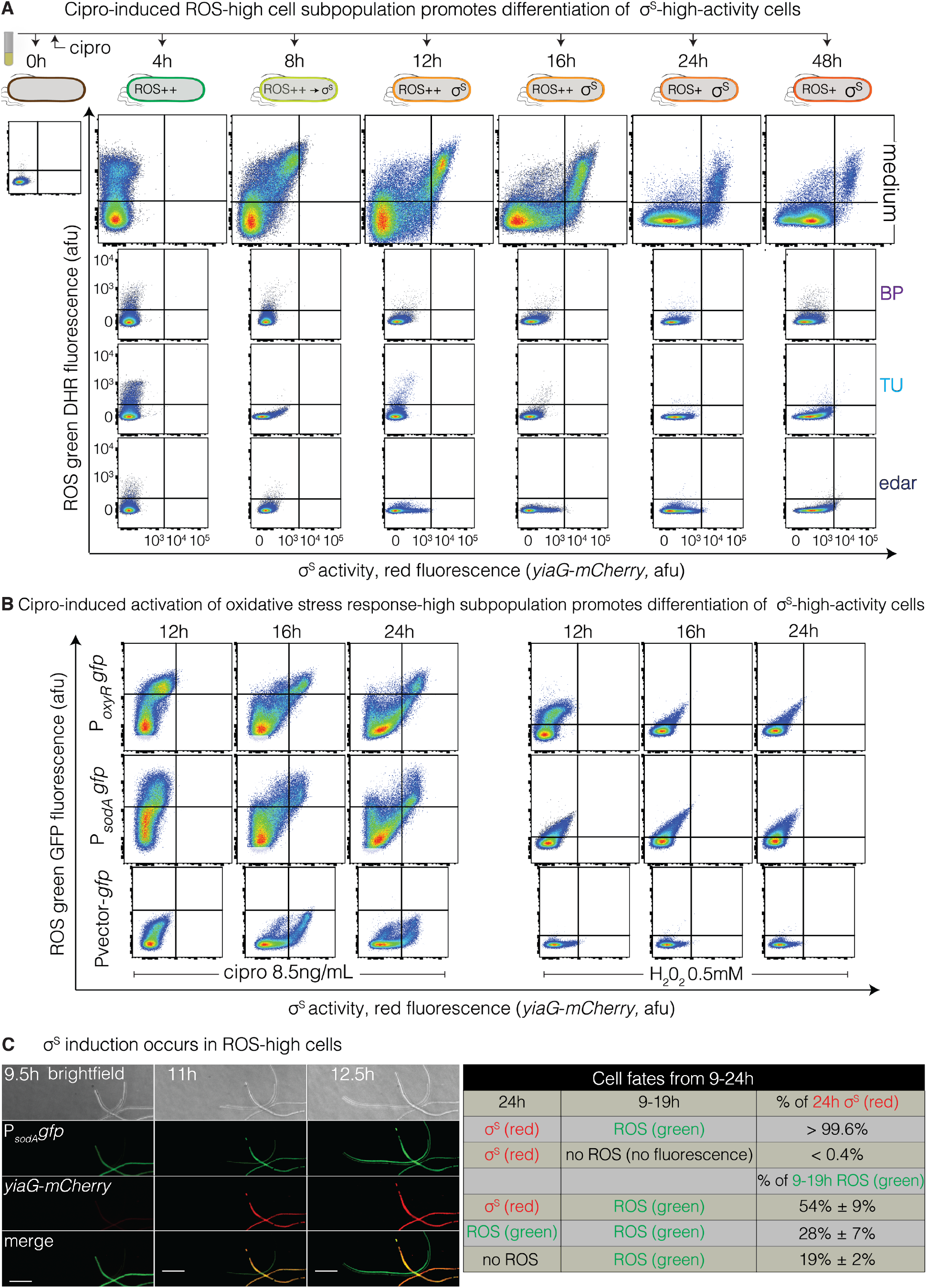
ROS-high Subpopulation Cells Become σ^S^ High-Activity Cells. (A) Cipro-induced ROS-positive cells give rise to many of the σ^S^ high-activity cells. Cells grown with subinhibitory cipro, with and without the ROS reducers 2, 2’ bipyridine (BP), thiourea (TU), or edaravone (edar) were collected serially and analyzed by flow cytometry for ROS-positive cells (DHR dye) and σ^S^ activity using the *yiaG-mCherry* σ^S^-response reporter. ROS-high cells precede σ^S^-high cells, and the presence of double-positive cells indicates that many of the σ^S^-active red cells arise from ROS-high green cells. Double positives are seen in the upper right quadrant, times 8-48h. Data, representative of 3 experiments. (B) Cipro-induced ROS-high cells, detected by OxyR and SodA oxidative-stress-response reporters, promote σ^S^ activation. Live cells carrying the *PoxyRgfp* or *PsodAgfp* oxidative-stress-response reporters and the *yiaG-mCherry* σ^S^-response reporter were grown with subinhibitory cipro, or 0.5mM H_2_O_2_ control (**Supplemental Discussion S3**), collected and analyzed by flow cytometry. Double-positive ROS-high, σ^S^-high cells indicate that many σ^S^-high cells had high ROS. The OxyR response is activated by endogenous H_2_O_2_ and SodA by superoxide. Data are representative of 2 experiments. (C) Most or all σ^S^ high-activity (red) cells arise from oxidative stress-response-activated green cells. Live-cell time-lapse imaging of cells carrying the *PsodAgfp* oxidative-stress-response reporter and *yiaG-mCherry* σ^S^-response reporter were grown with sub-inhibitory cipro for 8 hours, then imaged during growth for 12 additional hours using time-lapse deconvolution microscopy. Representative image and quantification show that essentially all σ^S^-active red cells at 24h arose from cells that were ROS-high and green at 9-19h (>99%). Also, most (54%) but not all (28%) ROS-high cells at 19h have become σ^S^-active at 24h, and some ROS-high cells at 19h lose their ROS by 24h (19%). Scale bar, 10μM. Mean ± range of 2 experiments tracking ≥ 250 cells. See also Figure S5, and Movie S1.

We found that ROS-high cells appear before σ^S^-active cells. The high-ROS cell subpopulation is detectable with ROS dye DHR (green) and flow cytometry at 4 hours after cipro is added, when no σ^S^ activity from a σ^S^-response fluorescence reporter (red) is detectable (4h, Figure 4A, **Supplemental Discussion S2**). Then, double-positive cells, dyed for ROS (green) and σ^S^ activity (red) develop between 8 and 16 hours (Figures 4A and S5E), implying that at least some σ^S^-high cells begin as ROS-high cells. At 24h, the time at which cells were harvested for sorting (Figure 3A)/mutagenesis assays (Figures 1A,B), many double-positive ROS-/σ^S^-high cells (upper right quadrant, Figure 4A 24h), and also some σ^S^-high single-positive cells, were present (lower right quadrant, Figure 4A 24h). Whether the σ^S^-single-positive cells at 24 hours originated from (had been) ROS-high cells before 24 hours was unclear. We used live-cell imaging with fluorescence-reporter genes (green) for two different oxidative stress responses, in cells that also carry the red σ^S^-response reporter, to follow individual cells over time from their burst of ROS to σ^S^-response induction. The reporters are transcriptional GFP fusions (Zaslaver et al., 2006) for *oxyR* (peroxide) and *sodA* (superoxide) responses, and both show double-positive and some σ^S^-single-positive cells at 24 hours (Figure 4B, green, the peroxide control discussed **Supplemental Discussion S3**).

Using the *sodA* reporter with live-cell time-lapse deconvolution microscopy, we show that essentially all red σ^S^-active cells arose from oxidative-stress-response-activated green cells (> 99%, Figure 4C and Movie S1). Some of the σ^S^-response-induced (red) cells showed decreased ROS (green) after σ^S^-response induction (Figure 4C and Movie S1), suggesting amelioration of high ROS levels by the σ^S^ response. The data demonstrate that cipro induces MBR by activating differentiation of a subpopulation of ROS-high cells that become mutable, σ^S^-response-high gamblers that generate most of the antibiotic cross-resistant mutants (Figure 3A).

We investigated how ROS activate the σ^S^ response in subpopulation cells (Figure 5). σ^S^ is regulated at multiple levels (Battesti et al., 2011) including translational upregulation by small RNAs (sRNAs). The ArcZ, RprA, and DsrA sRNAs activate σ^S^ translation assisted by the Hfq RNA chaperone (Battesti et al., 2011). We found that DsrA and ArcZ, but not RprA, are required for both cipro induction of σ^S^ protein (Figure 5A), and differentiation of the σ^S^-high gambler subpopulation (Figure 5B). The Hfq RNA chaperone is also required for cipro induction of σ^S^ protein (Figure 5A), σ^S^-response activity (Figure 5B), and RifR mutagenesis (Figure 5C). Moreover, the requirement for Hfq in RifR mutagenesis can be substituted by artificial upregulation of σ^S^ from a plasmid, which restored 86% ± 10% of RifR mutagenesis to Δ*hfq* cells (Figure 5C, controls Figure S2A,B). The data indicate that the Hfq RNA chaperone promotes cipro induction of mutagenesis mostly or wholly by promoting σ^S^-response induction, presumably via the ArcZ and DsrA sRNAs. Knock out of *hfq* in Δ*arcZ* or Δ*dsrA* cells causes no further reduction in σ^S^-β-galactosidase (Figure 5A), supporting this role of Hfq. Furthermore, we found that cipro induced *dsrA* and *arcZ* transcription by 2.3 ± 0.3- and 53 ± 3-fold, respectively in log phase (Figure 5D), shown with transcriptional *lacZ* fusions to the promoters of the *dsrA* and *arcZ* genes (Mandin and Gottesman, 2010; Sledjeski et al., 1996); and this transcriptional upregulation was ROS dependent, and was reduced by ROS reducers TU, BP, and edaravone (Figure 5D). The data demonstrate that cipro-induced ROS promote transcription of sRNAs DsrA and ArcZ, which, with RNA chaperone Hfq, upregulate σ^S^, activating the general stress response. Thus, these sRNAs underlie the differentiation of ROS-high cells into the σ^S^-active gambler subpopulation (Figure 5E) that generates antibiotic cross-resistant mutants.

**Figure 5.**
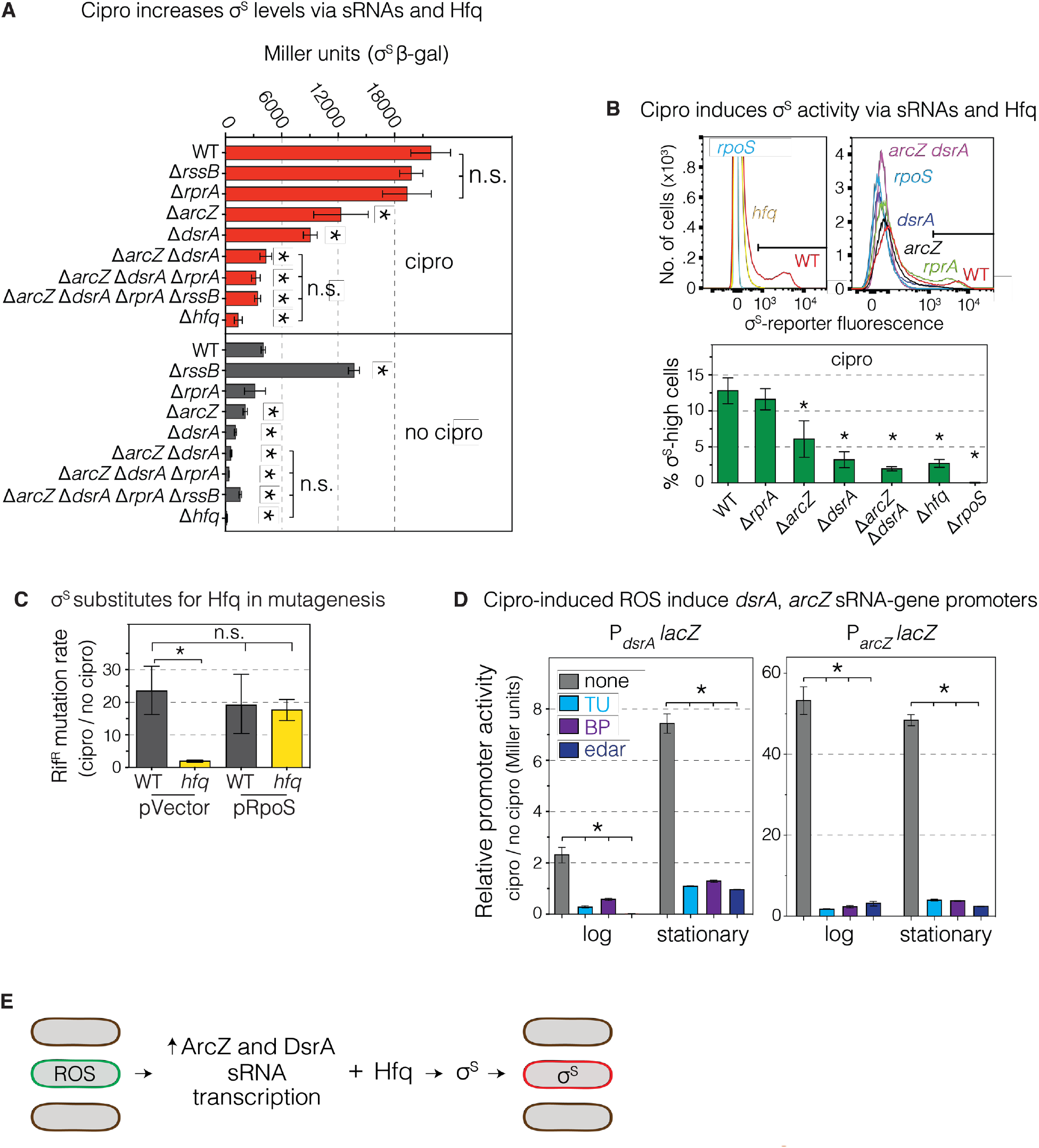
ROS induce transcription of sRNAs that upregulate σ^s^ and the general stress response. (A) Small RNAs (sRNAs) DsrA and ArcZ and the RNA chaperone Hfq are required for cipro-induction of σ^S^ protein. DsrA and ArcZ, assisted by Hfq, promote translation of *rpoS* mRNA to σ^S^ protein (Battesti et al., 2011), implying translational upregulation of σ^S^ response by cipro-generated ROS. σ^S^-β-galactosidase activity reports σ^S^ production and degradation (Wassarman et al., 2001). The data imply that cipro induction of σ^S^ protein occurs via sRNA-facilitated translation. Because RssB facilitates degradation of σ^S^ protein, the increase of σ^S^ protein levels in Δ*rssB* cells without cipro, but not with, implies that reduction of σ^S^ degradation is also part of how cipro-induced ROS upregulate σ^S^. Means ± SEM of 3 independent experiments. *Different from WT with cipro (top half) or WT without cipro (bottom half) at *p*<0.01, one-way ANOVA with Tukey’s post-hoc test. (B) DsrA, ArcZ, and Hfq mediate cipro induction of σ^S^-response activity in stationary-phase cells. Representative flow cytometry histograms showing the loss of σ^S^ high-activity cells in *dsrA, arcZ* and *hfq* null mutants. Means ± SEM of 3 independent experiments. * *p*<0.01, one-way ANOVA with Tukey’s post-hoc test; n.s. not significant. (C) Artificial upregulation of σ^S^ substitutes for Hfq in cipro-induced mutagenesis. Thus, the major role of Hfq in mutagenesis is upregulation of σ^S^; if σ^S^ is otherwise supplied, Hfq is no longer needed. RifR mutation rates quantified with or without sub-inhibitory cipro. Means ± range of 2 independent experiments. \**p*<0.01, one-way ANOVA with Tukey’s post-hoc test; n.s. not significant. (D) Cipro-induced ROS upregulate transcription from the promoters of the *dsrA* and *arcZ* sRNA genes, quantified as beta-galactosidase activity from *P_dsrA_lacZ* and *P_arcZ_laZ* transcriptional-fusion reporters in cells grown with or without sub-inhibitory cipro, with or without ROS reducers TU, BP, or edaravone. Means ± range of 2 independent experiments. \**p*<0.01, one-way ANOVA with Tukey’s post-hoc test. (E) Summary: Cipro-induced ROS in subpopulation cells induce transcription of the DsrA and ArcZ sRNAs which, with the Hfq RNA chaperone, upregulate σ^S^ in the same ROS-high cells (Figure 4).

There may be an additional component of σ^S^ upregulation by inhibition of σ^S^-protein degradation. One of the multiple ways that the σ^S^ response is kept “off” in unstressed cells is via RssB, which delivers σ^S^ protein to the ClpXP protease for degradation. Using a *rpoS-lacZ* reporter that makes a σ^S^-β-galactosidase fusion protein with an intact RssB-binding region (Zhou and Gottesman, 2006), we see that deletion of *rssB* increased σ^S^-β-galactosidase activity in cells untreated with cipro, but not in cipro-treated cells (Figure 5A), implying that detectable RssB-mediated σ^S^ degradation ceases after cipro treatment. The data could mean either that σ^S^-protein degradation may be inhibited by cipro/ROS, or that the upregulation of translation of σ^S^ by the DsrA and ArcZ sRNAs might cause saturation of RssB-mediated σ^S^-protein degradation.

### ROS Induced via SOS Response and Ubiquinone

Although antibiotics including fluoroquinolones induce ROS in cells (Dwyer et al., 2015), the ROS-induction pathway is only partly characterized (indicated by a “?” left of ROS in Figure 3I). Our data above show that cipro interaction with its target topoisomerases (DSB formation) is required for ROS formation (Figure S3I), that ROS arise in a cell subpopulation (Figure 2B), and previous work implicated Fenton chemistry and components of electron transfer (Dwyer et al., 2015). We discovered that the SOS response and ubiquinone oxidoreductase are required for induction of the ROS-high cell subpopulation by cipro.

First, we examined mutagenesis in cells defective for components of three electron-transfer chain (ETC) protein machines shown to promote the σ^S^ response during starvation-stress-induced MBR: NuoC (ubiquinone oxidoreductase I, an ETC “complex I” subuint), UbiD (biosynthesis of ubiquinone), and CyoD (a subunit of cytochrome bo’ oxidase, an ETC “complex II” subunit) (Al Mamun et al., 2012). Whereas CyoD and NuoC were not required for RifR or AmpR mutagenesis, UbiD (ubiquinone) was required for most of both (Figure 6A, controls Figure S2A,B). Moreover, UbiD (and ubiquinone) appear to act upstream of σ^S^-response induction in mutagenesis, in that artificially induced production of σ^S^ substituted for UbiD, restoring most or all of RifR mutagenesis to Δ*ubiD* mutant cells (87% ± 16% restored, Figure 6B). We observed reduction of σ^S^ accumulation and σ^S^ activity in *ubiD* null-mutant (ubinquinone-deficient) cells (Figure 6C and 6D), demonstrating that ubiquinone, and by implication, electron transfer, are required for cipro induction of the σ^S^ response. By contrast, UbiD/ubiquinone was not required for SOS-response activity (Figure 6E). Together, the data show that ubiquinone is required for cipro-induced mutagenesis, which it allows by promoting induction of the σ^S^ response.

**Figure 6.**
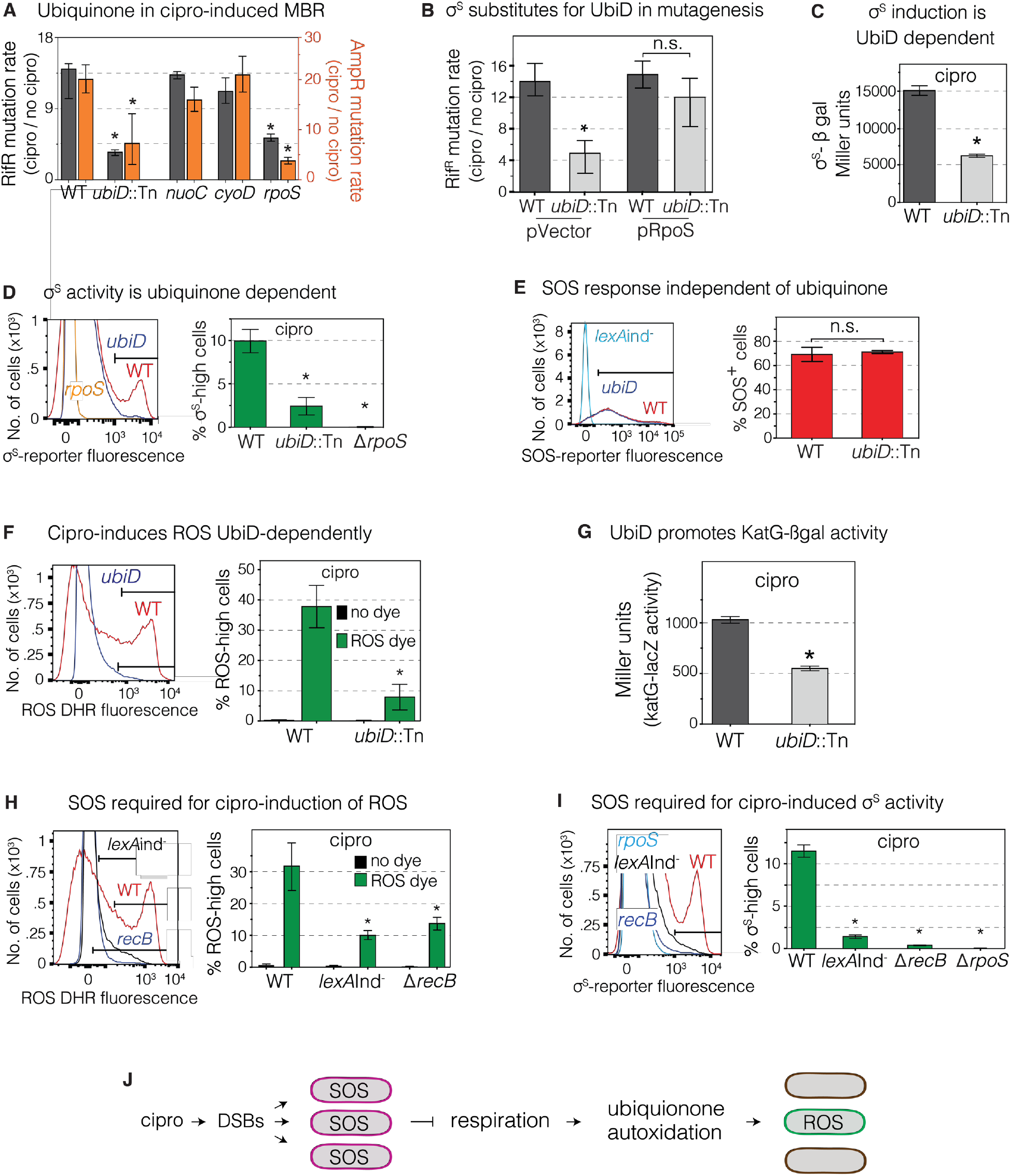
ROS Induction is SOS and Ubiquinone (Electron Transfer) Dependent. (A) Ubiquinone promotes cipro-induced RifR and AmpR mutagenesis. Left *y*-axis, gray bars (RifR); right *y*-axis, orange bars (AmpR). Mutants were grown at their respective subinhibitory cipro concentrations. Means ± 95% confidence intervals (CIs) of ≥ 3 independent experiments. *Different from wild-type (WT) at *p*<0.01, one-way ANOVA with Tukey’s post-hoc test of natural-log transformed data. (B) Artificial upregulation of σ^S^ substitutes for UbiD in mutagenesis. The data imply that UbiD promotes mutagenesis by upregulation of σ^S^, and so is not needed when σ^S^ is otherwise upregulated. Means ± 95% CIs of ≥ 3 independent experiments. *Different from WT at *p*<0.01, one-way ANOVA with Tukey’s post-hoc test of natural-log transformed data; n.s. not significant. (C) Cipro-induced σ^S^ protein accumulation is promoted by UbiD. σ^S^-β-galactosidase activity reports σ^S^ protein levels. Means ± range of 2 independent experiments. *Different from wild-type at *p*<0.01, two-tailed Student’s *t*-test. (D) High cipro-induced σ^S^ activity requires UbiD. Flow-cytometry assay of σ^S^ activity as fluorescence from the *yiaG-yfp* σ^S^-response reporter shows loss of the σ^S^ high-activity cell subpopulation in a *ubiD* null mutant. Means ± SEM of 2 independent experiments. *Different from wild-type at *p*<0.01, one-way ANOVA with Tukey’s post-hoc test. (E) Cipro induction of the SOS response does not require UbiD. SOS activity in stationary-phase cells measured by flow cytometry as fluorescence from SOS-reporter gene *P_sulA_mCherry*. Representative flow cytometry histograms show no difference in SOS activity in *ubiD-null* cells. Means ± range of 2 independent experiments. n.s. not significantly different from WT, two-tailed Student’s *t*-test. (F) Cipro induction of ROS requires UbiD. ROS-positive cells in log phase dyed with dihydrorhodamine 123 (DHR). Means ± SEM of 3 independent experiments. *Different from WT at *p*<0.01, one-way ANOVA with Tukey’s post-hoc test. (G) UbiD promotes cipro induction of the H_2_O_2_ responsive *katG-lacZ* fusion. WT and *ubiD* strains grown with sub-inhibitory cipro were assayed for *katG-lacZ* activity in log-phase. Means ± range of 2 independent experiments. *p<0.01, two-tailed Student’s *t*-test (H) The SOS response is required for cipro induction of the ROS-high cell subpopulation. SOS non-inducible *lexA*Ind^-^ and *recB* cells, which are defective in SOS induction by DSBs, do not generate the ROS-high cell subpopulation. Means ± range of 2 independent experiments. *Different from WT at *p*<0.01, one-way ANOVA with Tukey’s post-hoc test. (I) The SOS response is required for cipro induction of the σ^S^ high-activity cell subpopulation, which is prevented in SOS non-inducible *lexA*ind^-^ cells, and *recB* cells deficient in DSB-induction of the sOs response. Means ± range of 2 independent experiments. *Different from wild-type at *p*<0.01, one-way ANOVA with Tukey’s post-hoc test. (J) Model for cipro-induction of ROS via the SOS response and ubiquinone. Cipro causes DSBs, which activate the SOS response in all cells. The SOS response can slow aerobic respiration (Swenson and Schenley, 1974), which promotes autoxidation of ubiquinone (Gonzalez-Flecha and Demple, 1995; Skulachev, 1998), we suggest in a cell subpopulation, which generates ROS in that population. Because UbiD is not needed for cipro induction of the SOS response, but UbiD and SOS are required for cipro-induction of ROS and σ^S^-high cells, we infer that SOS acts upstream of electron transfer/ubiquinone in inducing the ROS-high cell subpopulation that becomes the σ^S^-high cell subpopulation (Figure 5). See Figure S4 and Tables S1 and S2.

We found that the ubiquinone role is specifically in the induction of ROS. Ubiquinone, or coenzyme Q, functions in the aerobic ETC by mediating oxido-reduction cycles required for ATP energy production (Meganathan and Kwon, 2009). We found that ubiD-defective cells showed severely reduced ROS generation in cipro compared with wild-type cells (32% ± 9% ROS-high cells in wild-type, and 8% ± 4% in *ubiD-null* cells, Figure 6F). ubiD-null-mutant cells also displayed reduced *katG-lacZ* activity, a reporter activated by H_2_O_2_ (Liu and Imlay, 2013) (Figure 6G). The data show ubiquinone-promoted induction of ROS, which are required for the cipro induction of the σ^S^ response.

Perhaps surprisingly, the SOS response is also required for cipro induction of ROS and the σ^S^ response. SOS-non-inducible (lexAind^-^) cells, and cells lacking RecB, which is required for SOS induction by DSBs (McPartland et al., 1980), showed reduced induction of ROS (Figure 6H) and the σ^S^ response (Figure 6I) by cipro, contributing to at least 70% ± 4% of ROS-high subpopulation cells (Figure 6H). Because ubiquinone was not needed for SOS-response induction by cipro (Figure 6E), we can infer that SOS acts upstream of, or in parallel with, ubiquinone in ROS induction (Figure 6H); not downstream of ubiquinone, which is not needed for SOS induction (Figure 6E). The SOS response, induced by UV light, was reported to inhibit aerobic respiration (Swenson and Schenley, 1974), and, also in assays without cipro, slowing of respiration increased autoxidation of quinols leading to superoxide production (Gonzalez-Flecha and Demple, 1995; Skulachev, 1998). Together with our data using cipro, these data support a model in which SOS activation may inhibit the ETC leading to ROS generation (Figure 6J). SOS action upstream of the ubiquinone contribution to ROS generation (Figure 6J) is harmonious with our data (Figure 6E). The data support a model in which the cipro-induced SOS response may inhibit/slow aerobic respiration, per (Swenson and Schenley, 1974), in only a subpopulation of cells, allowing autoxidation of ubiquinone to generate high ROS levels in those cells (Figure 6J).

### Multi-Chromosome Cells Allow Evolvability

We prevented cipro from inducing multi-chromosome cells by deletion of the *sulA* gene (Figure 7), the product of which is induced by SOS and inhibits cell division, causing multichromosome cells during a DNA-damage response (Huisman and D’Ari, 1981). SulA inhibits polymerization of the microtubule-like “Z-ring”, which pinches off daughter cells (Bi and Lutkenhaus, 1993). Because SulA does not block DNA replication or elongation of the rodshaped *E. coli* cells, cells grow long and “filamentous” or snake-like with multiple chromosomes in them (Huisman and D’Ari, 1981) when SOS and SulA are induced (Figure 7A and B). We used microscopy and a protein that marks the chromosome as a fluorescent focus (Figure 7A-C) (Joshi et al., 2013), to show that Δ*sulA* cells do indeed make much shorter cells in low-dose cipro (Figure 7D-F), that these have fewer chromosomes per cell (Figure 7E,F), and are deficient in mutagenesis (Figure 7G). Without cipro only 1% ± 0.7% of exponential wild-type cells have four or more chromosome copies (Figure 7C), so we defined a multi-chromosome cell as those with ≤4 chromosome copies. With cipro, 33% ± 2% of wild-type cells have ≥4 chromosome copies (Figure 7B). By contrast, Δ*sulA* cells show much reduced cell length and chromosome content (Figure 7D-F). We find that these Δ*sulA* shorter cells show reduced cipro-induced RifR and AmpR mutant production (Figure 7G), showing 67% ± 5% and 70% ± 5% fewer mutants, respectively. Further, deletion of the *ruvC* HR-repair gene from Δ*sulA* mutant cells caused no further decrease in mutagenesis, indicating that SulA acts in the same HR-dependent MBR mutation pathway as RuvC. Thus, SulA, and the filamented, multi-chromosome state, are required for cipro-induced MBR.

When exposed to low-dose cipro, *E. coli* forms long, multi-chromosome cell “filaments” that “bud off” small, normal-length daughter cells that produce high frequencies of cipro-resistant mutants (Bos et al., 2015). These data suggested that multi-chromosome cells might promote adaptation by coupling mutagenesis, which can generate resistance but also deleterious mutations, with recombination or allele sharing, which might mitigate deleterious effects of many recessive mutations, allowing the multi-chromosome cell to produce resistant, surviving/adapted progeny (Bos et al., 2015). We tested directly whether the multi-chromosome state is required for cipro-induced mutant production, and explored whether it could promote adaptation via mutagenesis.

**Figure 7.**
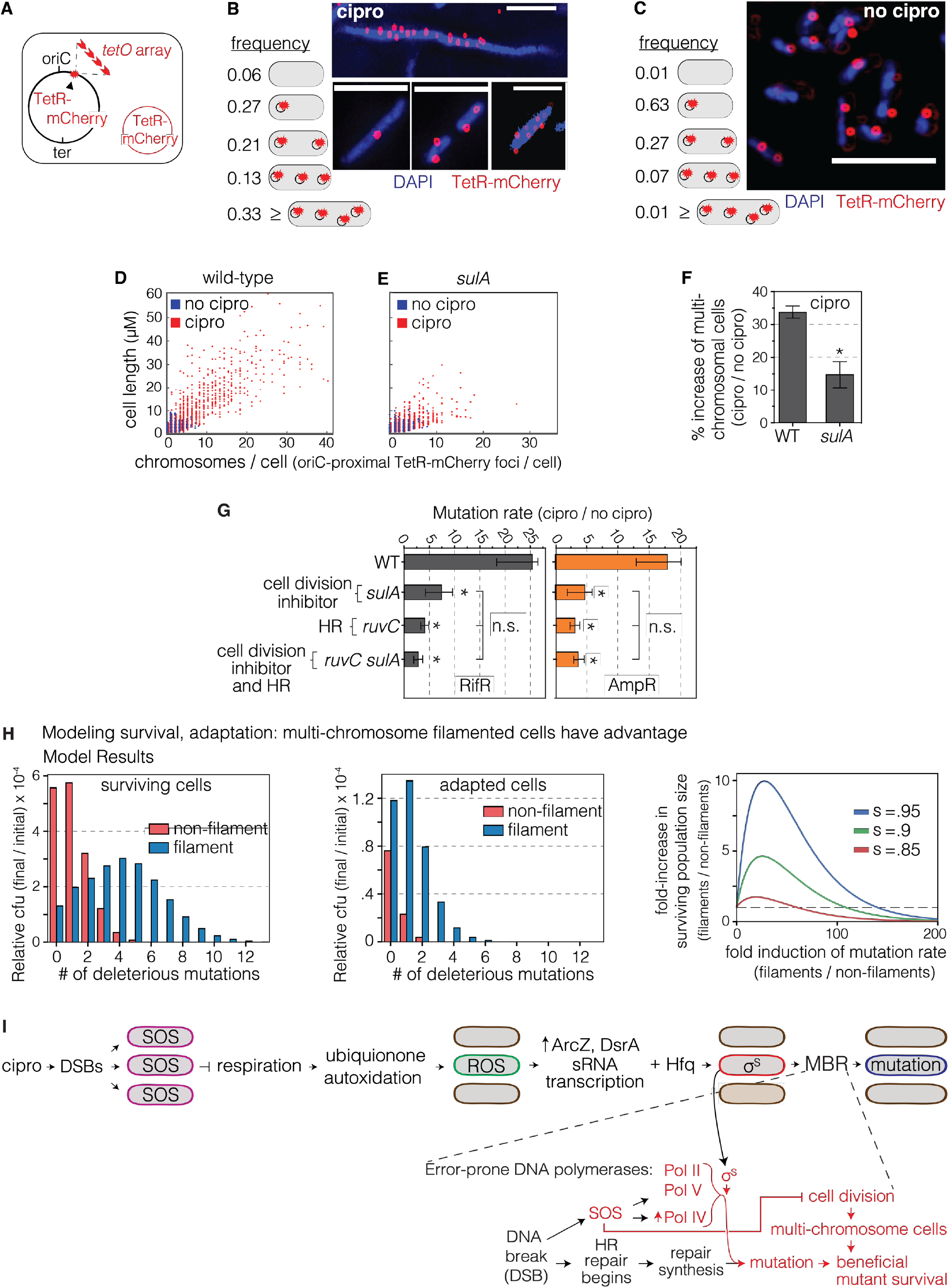
Multi-chromosome Bacterial Cells Promote Cipro-induced Mutagenesis. (A) Scheme for labeling chromosomes as red fluorescent foci using a chromosomal *tetO* array bound by Tet-repressor-mCherry (TetR-mCherry) in a replication-origin (*oriC*)-proximal site. Red circle, plasmid that produces TetR-mCherry. Multiple TetR-mCherry foci represent the approximate number of *ori*-proximal chromosomal equivalents in a bacterial cell (Joshi et al., 2013). (B) More than 33% of cells grown in low-dose, sub-inhibitory cipro carry multiple chromosomes, defined as ≥4 chromosomes per cell, quantified as TetR-mCherry foci, per (A), 1919 cells counted. Representative fluorescence images of DAPI-stained log-phase WT cells. White scale bar, 5 μm. (C) Fewer than 1% of cells grown without cipro have multiple chromosomes, quantified as TetR-mCherry foci, per (A), 3915 cells counted. Representative fluorescence images of DAPI-stained log-phase cells. White scale bar, 5 μm. (D-F) Cipro induction of the multi-chromosome, filamented cell state requires SOS-induced cell-division inhibitor, SulA. Scatterplots show the microscopically determined distributions of cell lengths (μM) and chromosome (TetR-mCherry) foci per cell in cells grown with and without low-dose cipro. Data from 3 independent experiments. (D) Cipro induction of filamented, multi-chromosome cells. (E, F) SulA is required for the cipro-induction of long, multi-chromosome cells. 98% of untreated cells show ≤4 chromosomes per cell. Means ± SEM of 3 independent experiments. *Different from WT at *p*<0.001, one-way ANOVA with Tukey’s post-hoc test; n.s. not significant. (G) The SulA-dependent multi-chromosome state promotes cipro-induced mutagenesis. SulA and RuvC act in the same MBR pathway (are epistatic). Means ± 95% CIs of ≥ 4 independent experiments. *Different from wild-type at *p*<0.001, one-way ANOVA with Tukey’s post-hoc test of natural-log transformed data; n.s. not significant. (H) Mathematical model shows that multi-chromosome “filamented” cells have a large advantage for adaptation and survival at high mutation rates. Left panel, expected relative cfu of all surviving cells (adapted and not-adapted) is plotted as a function of the number of deleterious mutations accumulated, for both filamented and non-filamented cells. Middle panel, the expected relative cfu of adapted cells is plotted as function of the number of deleterious mutations accumulated. Right panel, formation of multi-chromosome filaments can increase the surviving population size when selection is harsh. The fold-increase of surviving population size due to filamentation is plotted as function of the fold increase in mutation rate due to filamentation, for several selection parameters. s – selection coefficient of the major stress (e.g., antibiotics). Model description and parameters in **Methods**. (I) Model: mechanism of cipro-induced transient differentiation of an evolvable gambler cell subpopulation that allows stress-responsive MBR without risk to most cells, facilitated by the multi-chromosome state. Left to right: cipro-binding to type-II topoisomerases causes DSBs that activate the SOS response throughout the cell population. SOS upregulates error-prone DNA polymerases and SulA, which inhibits cell division causing multi-chromosome cells. SOS also slows aerobic respiration, we suggest, in a cell subpopulation, which generates ROS promoted by autoxidation of ETC component ubiquinone in that subpopulation. The ROS activate transcription of σ^S^-upregulating sRNAs DsrA and ArcZ, which, with Hfq RNA chaperone, promote translation of *rpoS* mRNA to σ^S^ protein, thus activating the general stress response in the cell subpopulation, and allowing mutagenic DNA break repair (MBR) in those cells—a transient hypermutable state in gambler cells (red cells). The limitation of mutable cells to a subpopulation allows exploration of new phenotypes generated by genome instability in gamblers without risk to the whole population—a potential “bet-hedging” strategy. The multi-chromosome state promotes survival and adaptation of highly mutated cells by amelioration (complementation and reassortment) of deleterious recessive mutant phenotypes generated. See also Figures S2 and S3 and S4 and Tables S1 and S2.

We used mathematical modeling to address the hypothesis of Bos et al. (2015) that multichromosome cell filaments have an advantage over normal cells in adaptation to changing (stress) environments via mutagenesis. We formulated a mathematical model to test possible benefits of multi-chromosome cell filaments for rapid adaptation (**Methods**). Results of the model, presented in Figure 7H, show that increasing filament mutation rate can greatly increase the probability of both adaptation and survival of a chromosome in multi-chromosome filaments relative to non-filamented cells. This supports recent work showing that cooperation can accelerate complex adaptations (Obolski et al., 2017). Further, the advantage of the multi-chromosome state increases with increasing selection coefficient (Figure 7H). Selection coefficient represents, for example, the lethality of cipro in cipro-treated cells, which are under selection for cipro-resistance mutations. This model demonstrates that the multi-chromosome state is capable of facilitating adaptation by mutagenesis, illustrated in a model in Figure 7I.

## DISCUSSION

The mechanism of mutagenesis induced by the fluoroquinolone antibiotic cipro, revealed here (model, Figure 7I), demonstrates three new biological principles in mutagenesis (three headings below), and also unites quinolone-induced mutagenesis with a large canon of stress-induced mutagenesis mechanisms. Stress-induced mutagenesis mechanisms, from bacteria to human, are defined as mutation-producing mechanisms upregulated by stress responses (Fitzgerald et al., 2017). Activation of mutagenic DNA break repair (MBR, Figures 1E-J) by the general stress response (Figures 1F, 2C-E, H,I, 4, 5, 7I, couples mutagenesis to the σ^S^ response (Fitzgerald et al., 2017; Harris et al., 1994; Ponder et al., 2005; Rosenberg et al., 1994) causing mutations during stress. Fluoroquinolones were not known previously to provoke stress-induced/a^S^-dependent mutagenesis. Temporal regulation of mutagenesis by stress responses causes mutant generation preferentially when cells or organisms are maladapted to their environments—when stressed—potentially accelerating adaptation (Fitzgerald et al., 2017; Ram and Hadany, 2012, 2016).

### A Regulatory Role for ROS in Mutagenesis

We discovered a novel role of ROS in mutagenesis—their activation of the general stress response (Figures 2D, E, H, 3C, D, 4, 5), which allows stress-inducible MBR (Figure 1E,F and model, Figure 7I). ROS have long been known to promote mutagenesis by other more direct mechanisms, including by oxidation of guanines in nucleotide pools and in DNA to 8-oxo-dG. 8-oxo-dG pairs with A, causing G-to-T and T-to-G (A-to-C) mutations (Schaaper and Dunn, 1987), a signature that is *less* important for cipro-induced than spontaneous forward mutations *(ampD*, Figure S1 B,C), indicating a minor role. DNA base mispairs of 8-oxo-dG with A are also attacked by base-excision repair, which can lead to DNA breaks that are part of various antibiotics’ killing mechanisms (Foti et al., 2012; Kohanski et al., 2007; Rasouly and Nudler, 2018; Zhao et al., 2015), but we show are not an important source of the DSBs that drive cipro-induced MBR (Figure 2F,G). By contrast, we found that, surprisingly, the mutagenic role of ROS can be fully substituted by engineered production of σ^S^, the transcriptional activator (bacterial RNA-polymerase sigma factor) of the general/starvation stress response (Figure 2H), indicating that induction of the σ^S^ response is the predominant or only role of ROS in cipro-induced mutagenesis; if σ^S^ is supplied artificially, ROS are no longer needed. We found that ROS induce transcription of ArcZ and DsrA small RNAs (sRNAs, Figure 5), which, assisted by the Hfq RNA chaperone (Figure 5A-C), promote translation of the *rpoS* mRNA into σ^S^ protein (Battesti et al., 2011) (Figure 7I). This differs from a role of Hfq and another sRNA in mutagenesis via direct downregulation of translation of an mRNA encoding a mismatch-repair protein (Chen and Gottesman, 2017). We saw that induction of ROS by cipro precedes and is required for σ^S^-response activation (Figure 4), and that cells with high levels of ROS become σ^S^-response highly activated cells (Figure 4C, movie S1) that generate the mutants (Figure 3). These data highlight the centrality of stress-response-control of mutagenesis, even ROS-induced mutagenesis, and identify ROS as signaling molecules in this regulation. The DNA crosslinking agent mitomycin C also induces ROS that induce the σ^S^ response (Dapa et al., 2017), which might occur by similar mechanism with similar mutagenic consequences as shown here.

It is conceivable that several other ROS-promoted mutagenesis mechanisms may involve ROS upregulation of the general stress response, which can be mutagenic by allowing MBR (Lombardo et al., 2004; Ponder et al., 2005; Shee et al., 2011), downregulation of mismatch repair (Gutierrez et al., 2013), transposon movement (Ilves et al., 2001), and possibly other mechanisms. Because stress-response regulators, such as σ^S^, are non-redundant hubs in the MBR network (Al Mamun et al., 2012), these are attractive candidates for targets of proposed drugs to slow evolution of pathogens, averting evolution of resistance and evasion of immune responses (Al Mamun et al., 2012; Fitzgerald et al., 2017; Rosenberg and Queitsch, 2014). Moreover, the FDA-approved antioxidant drug edaravone behaved as such an “anti-evolvability” drug in our experiments (Figure 3C-E) and did so without reducing the killing power of cipro as an antibiotic (Figure 3H), providing a promising proof-of-concept. Reactive oxygen affects many aspects of the biology of cells from lipid and protein oxidation (Pradenas et al., 2012; Tamarit et al., 1998) to DNA damage and mutagenesis (reviewed above). The addition of stress-response activation causing MBR to this list identifies ROS as evolution-promoting signaling molecules.

### Mutagenesis in Transiently Differentiated Gamblers

Our data reveal that ROS, and then the general stress response, are induced strongly in a 1020% subpopulation of cipro-treated cells (Figures 2B-D and 4) that is transiently differentiated into a mutable state (Figure S5D), and produces most of the mutants (Figure 3A,C). Transient differentiation in bacterial subpopulations is a recognized potential evolutionary “bet-hedging” strategy, in which only some cells take the risk of “trying” a phenotype that may be advantageous in some environments and maladaptive in others (Norman et al., 2015; Veening et al., 2008). Transient phenotype examples include the persister state, which allows tolerance of lethal drugs but slows or halts proliferation (Balaban et al., 2004), competence for “natural transformation”— DNA uptake and incorporation into the genome (Chen and Dubnau, 2004), sporulation—a dormant but environmentally resistant state (Norman et al., 2015), and even programmed cell death (Amitai et al., 2009; Gonzalez-Pastor et al., 2003), hypothesized (Amitai et al., 2009; Rosenberg, 2009) or demonstrated (Gonzalez-Pastor et al., 2003) to benefit siblings of the sacrificed cells. The regulated/”programmed” limitation of mutagenesis itself—a major evolutionary driver—to a cell subpopulation appears to embed the apparent evolutionary strategy of environmentally tuned mutagenesis within another evolutionary strategy: “bet hedging” (Norman et al., 2015; Torkelson et al., 1997; Veening et al., 2008). Though transiently hypermutable cell subpopulations have been hypothesized (Hall, 1990; Ninio, 1991), supported by genetic evidence (Torkelson et al., 1997), and cells with stress responses linked to mutagenesis of unknown mechanism (Woo et al., 2018), our data provide the first isolation (Figure 3) of a hypermutable cell subpopulation in the act of mutagenesis, and show the defining, differentiating inputs: ROS and general stress-response activation (Figures 3 and 4) as well as the mutagenesis mechanism: MBR, illustrated in Figure 7I. We found that the subpopulation is relatively large at 10-20%, and that its transient differentiation (Figure S5D) is achieved by stress-response activation in the subpopulation cells, cell autonomously (Figure 4)—all novel mechanisms of potential means of promotion of the ability to evolve. Unlike “persisters,” these cells take the risk of inducing mutations, which can lead to heritable resistance to never-before-encountered antibiotics, and so could be called “gamblers.”

### Multi-chromosome Cells Promote Evolvability

We found that the multi-chromosome state, caused by the SOS-response-induced SulA inhibitor of cell division (Huisman and D’Ari, 1981), is required for cipro-induced mutagenesis (Figure 7). We observed SOS-dependent multi-chromosome cell “filaments” previously during low-dose cipro exposure and noted their “budding” off of small cells that produce high frequencies of cipro-resistant mutants (Bos et al., 2015). Here, we showed that “filamentation” is required for mutant production (Figure 7A-G). For cipro-induced mutagenesis by mutagenic repair of DNA doublestrand breaks (Figure 1E,F), induced here by cipro (Figure 1G-J, 2F and S1G), more than one chromosome is needed for DSB repair. Our modeling indicates that the multiple chromosomes may additionally facilitate survival and adaptation to stresses of highly mutating cells by cooperation (Obolski et al., 2017): sharing of gene products and/or alleles (recombination) between the chromosomes, masking deleterious recessive phenotypes (Figure 7H). Our data imply that “filamenting” cells may be biomarkers of rapid evolution. Bacteria like *Bacillus subtilis* undergo natural transformation—acquiring sibling cells’ DNA—simultaneously with, and activated by same Com stress response that upregulates a stress-induced mutagenesis mechanism (Sung and Yasbin, 2002). Thus, *B. subtilis* engages the adaptation-boosting combination of recombination and mutagenesis (Lenhart et al., 2012). Our data indicate that *E. coli*, which is famously incapable of natural transformation, may employ the same adaptation-accelerating strategy via multiple sibling chromosomes within one cell, rather than sibling DNA taken up exogenously. The data suggest that in addition to targeting stress-response regulators as an anti-evolvability drug strategy (Al Mamun et al., 2012; Fitzgerald et al., 2017; Rosenberg and Queitsch, 2014) and Figure 3C-H, dividing (and conquering) the multiple chromosomes might also prove to be an effective anti-evolvability, anti-pathogen therapeutic strategy.

## Supporting information

## AUTHOR CONTRIBUTIONS

JPP, LGV, OL-E, JB, RHA, CH, LH, and SMR conceived the project, advanced hypotheses and/or designed experimental approaches; JPP, LGV, YZ, AW, JL, JX, QM performed or guided the work; DB provided advice/assistance, JPP, PH and SMR wrote the manuscript.

## ACKNOWLEDGEMENTS

We thank S Gottesman, J Imlay, I Matic, and L Zechiedrich for kind gifts of *E. coli* strains, N Majdalani for guidance on sRNAs and σ^S^ activation, S Kozmin for advice on dose-dependent induction of mutagenesis, S Henikoff for helpful conversation, and KM Miller, and Meng Wang for improving the manuscript. This work was supported by the NIH Grants R35-GM122598 (SMR), R01-GM088653 (CH), R01-GM102679 (DB), R01-GM106373 (PJH), the Israeli Science Fund ISF 1568/13 (LH), and by the Integrated Microscopy Core at Baylor College of Medicine with funding from the NIH (dK56338, and CA125123), the Dan L. Duncan Comprehensive Cancer Center, RP160283 Baylor College of Medicine Comprehensive Cancer Training Program Postdoctoral Fellowship (DMF) and American Cancer Society Postdoctoral Fellowship 132206-PF-18-035-01-DMC (DMF); and the John S. Dunn Gulf Coast Consortium for Chemical Genomics. This project was supported by the Cytometry and Cell Sorting Core at Baylor College of Medicine with funding from the NIH (P30 AI036211, P30 CA125123, and S10 RR024574) and the expert assistance of Joel M. Sederstrom.

## STAR★Methods

### Key Resources Table

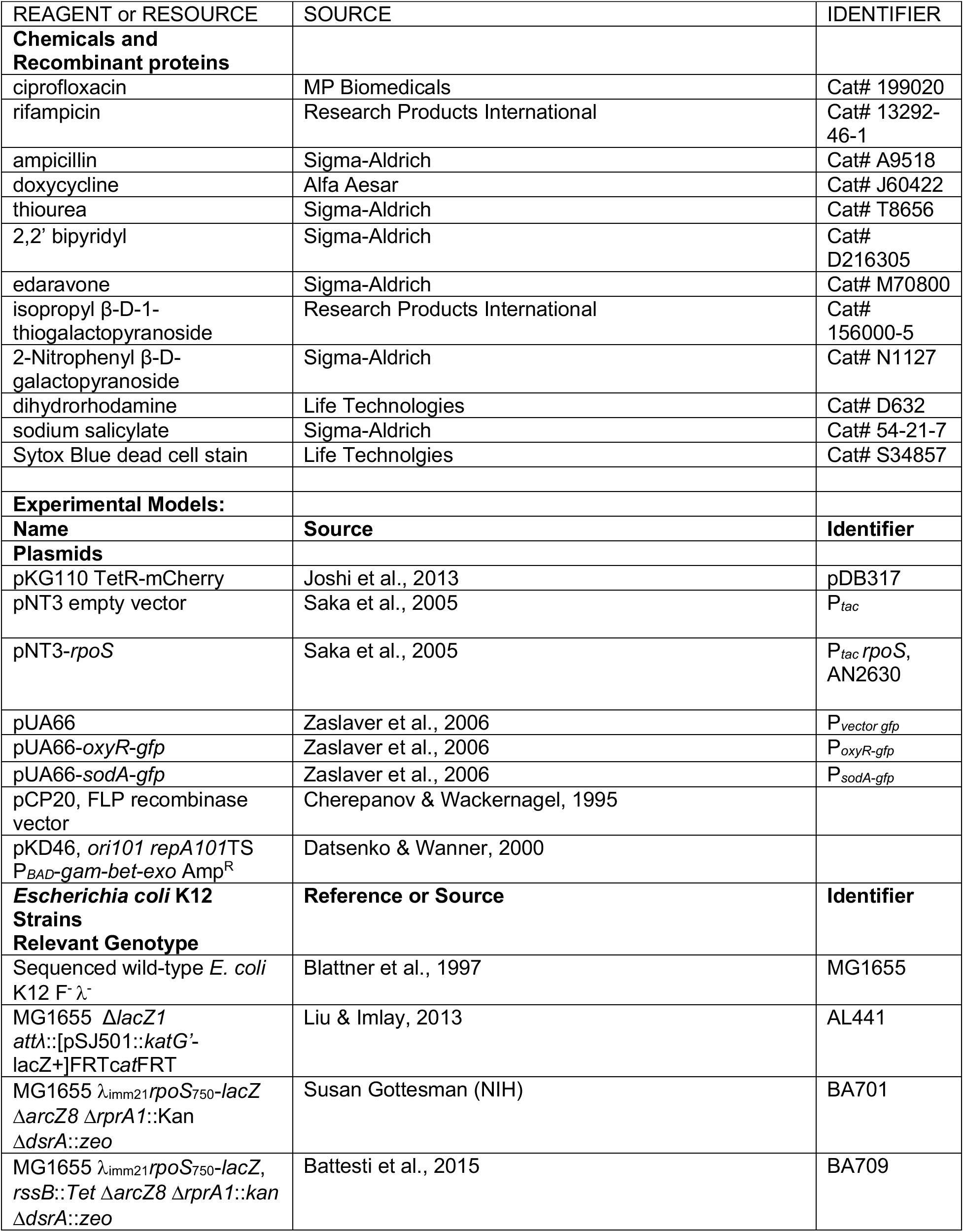
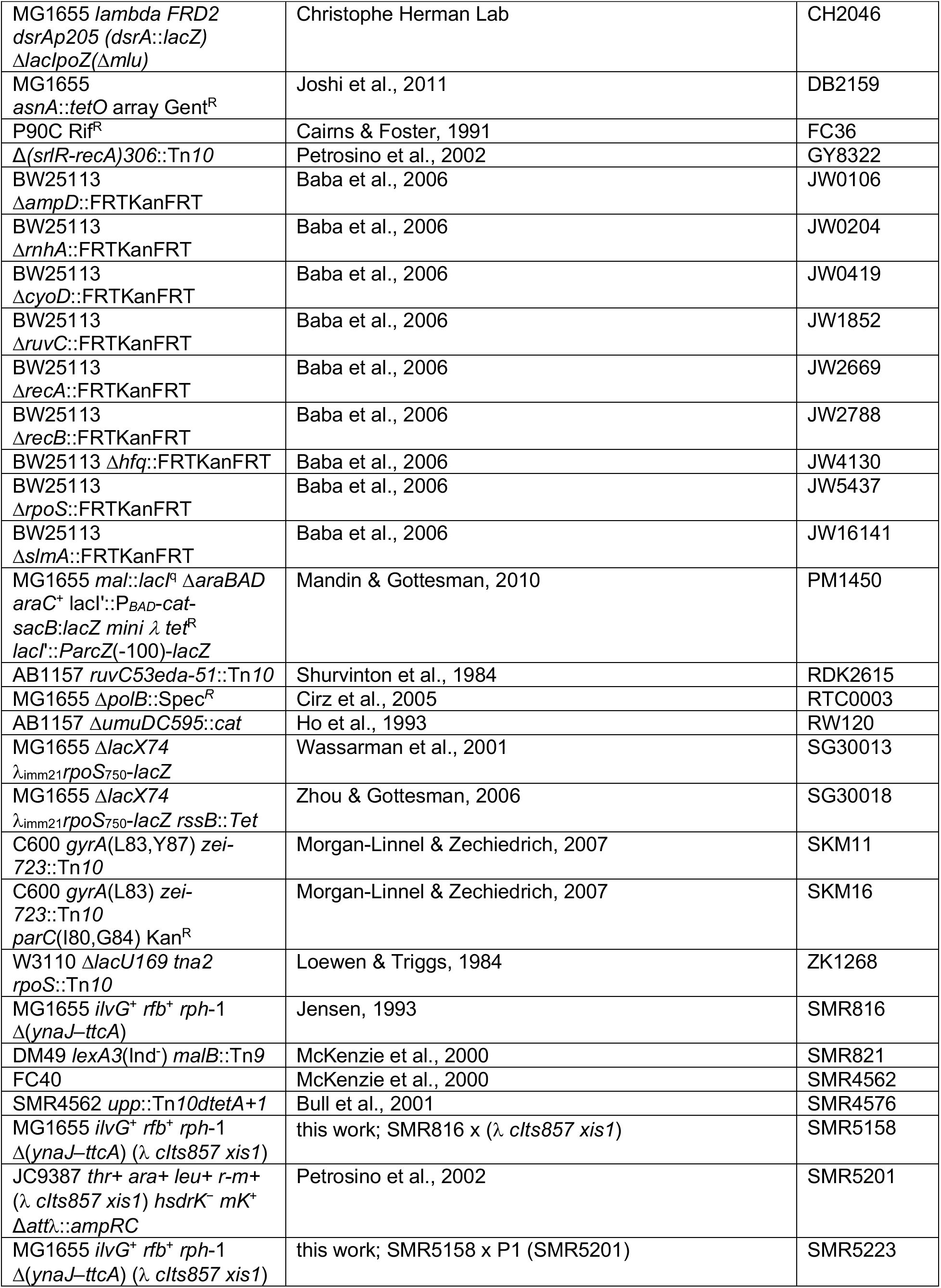
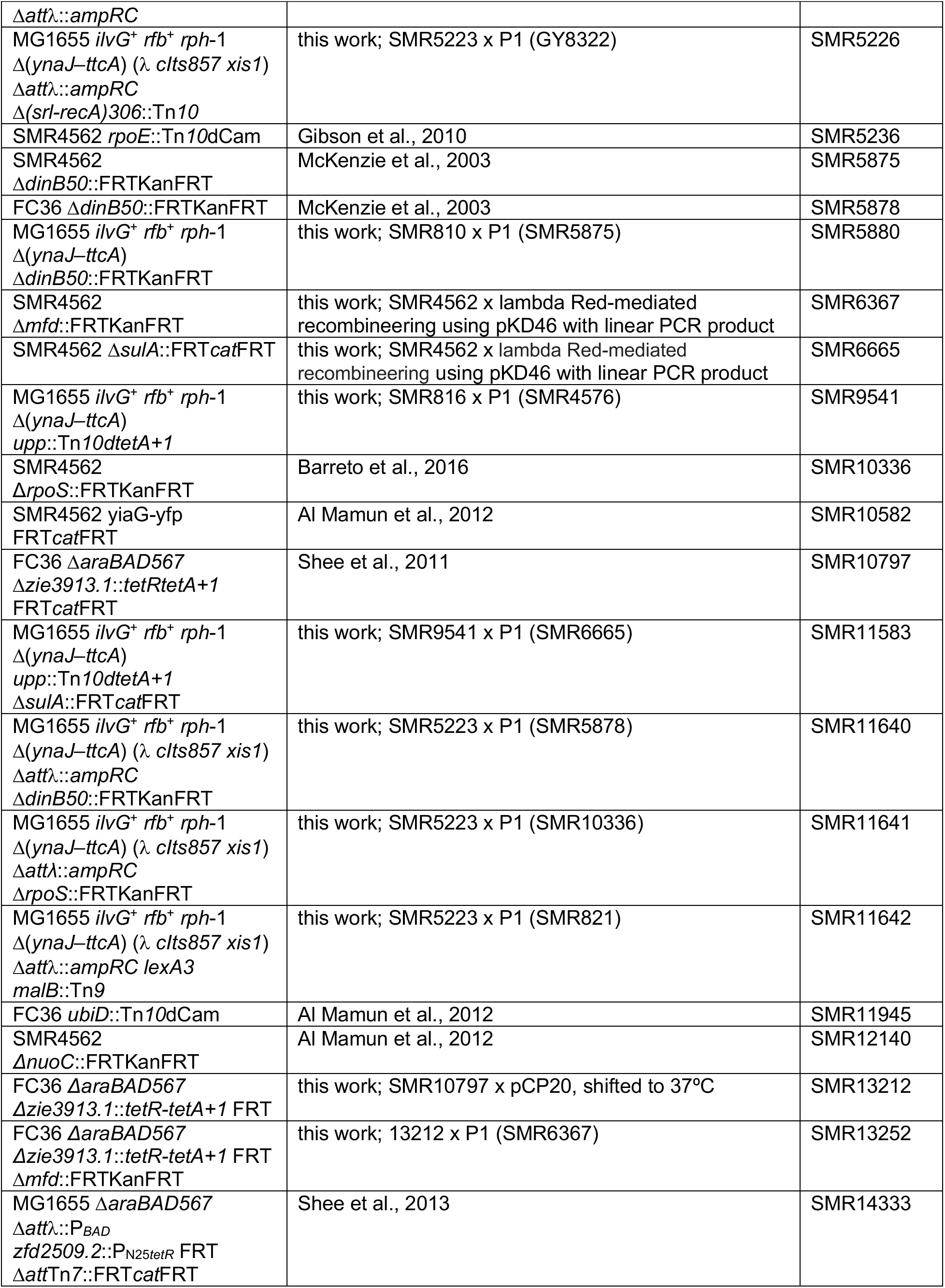
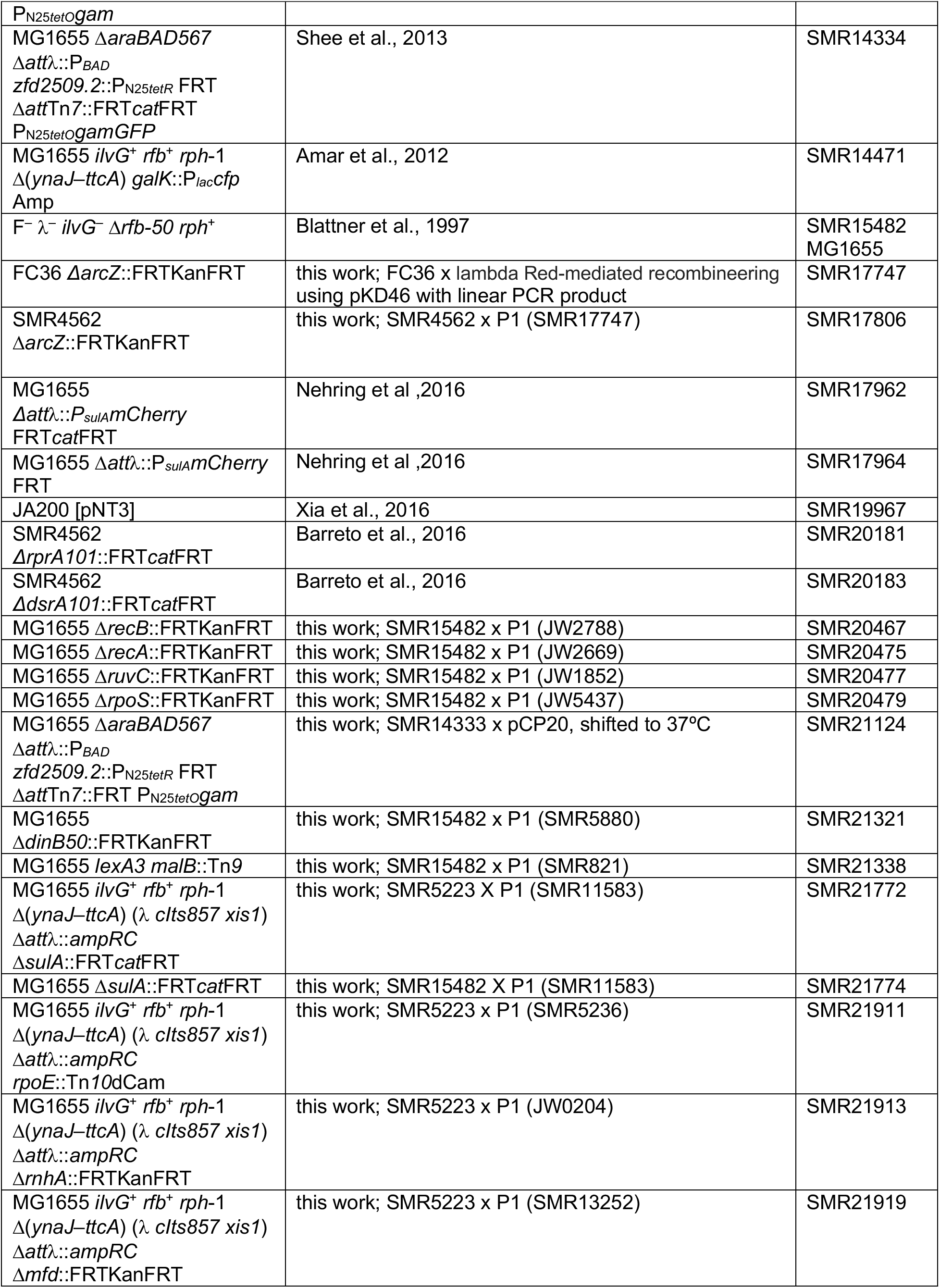
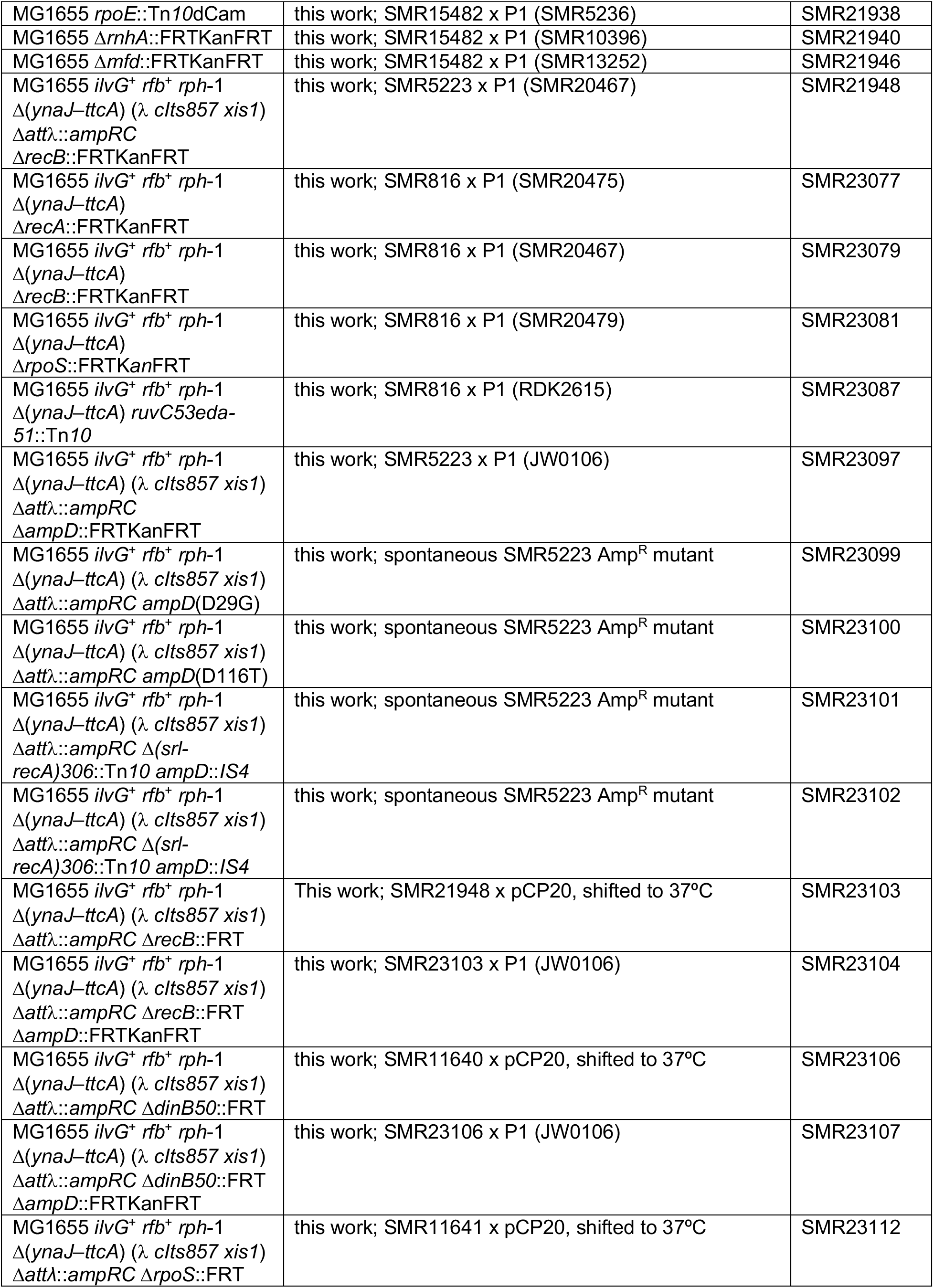
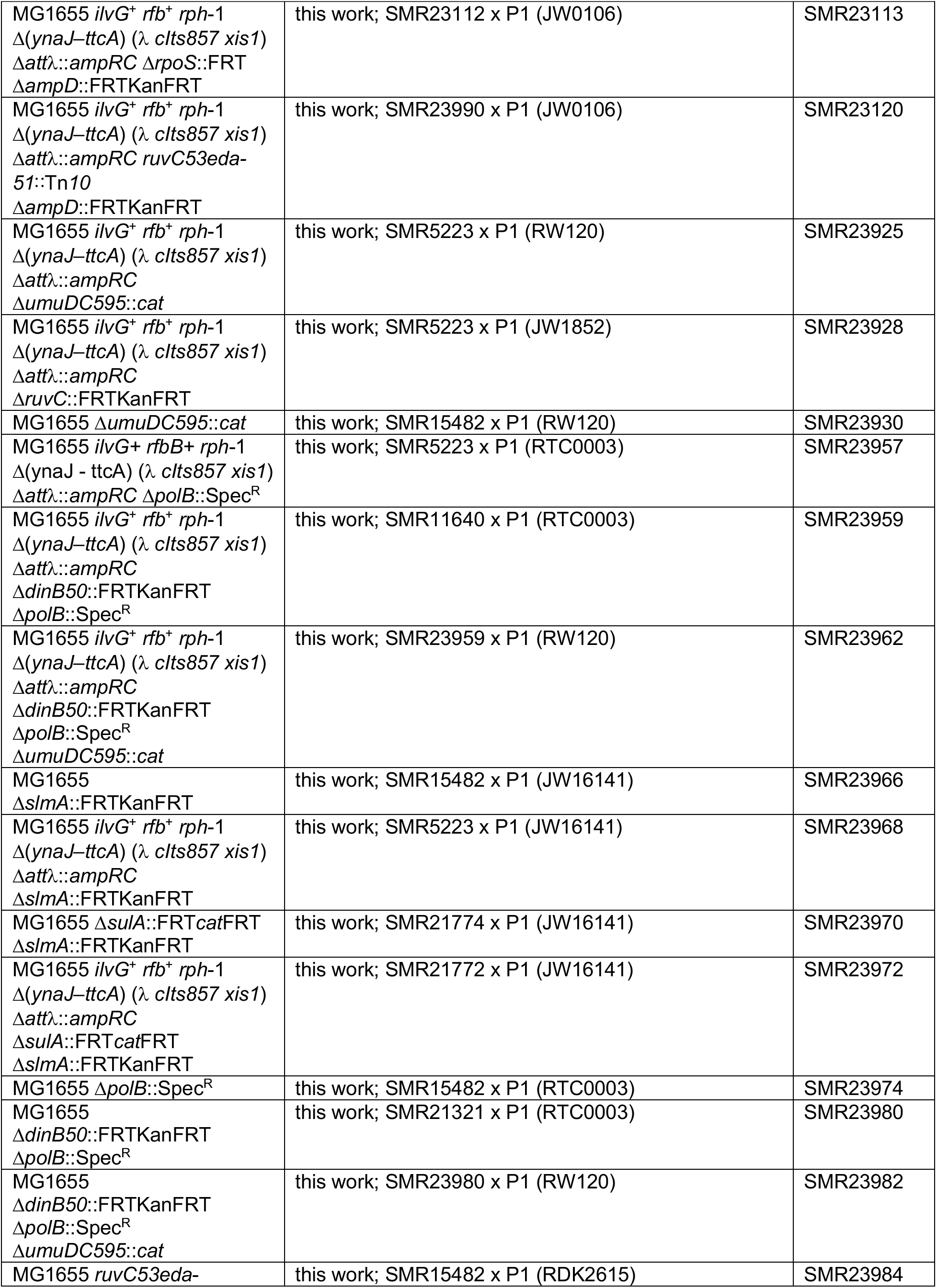
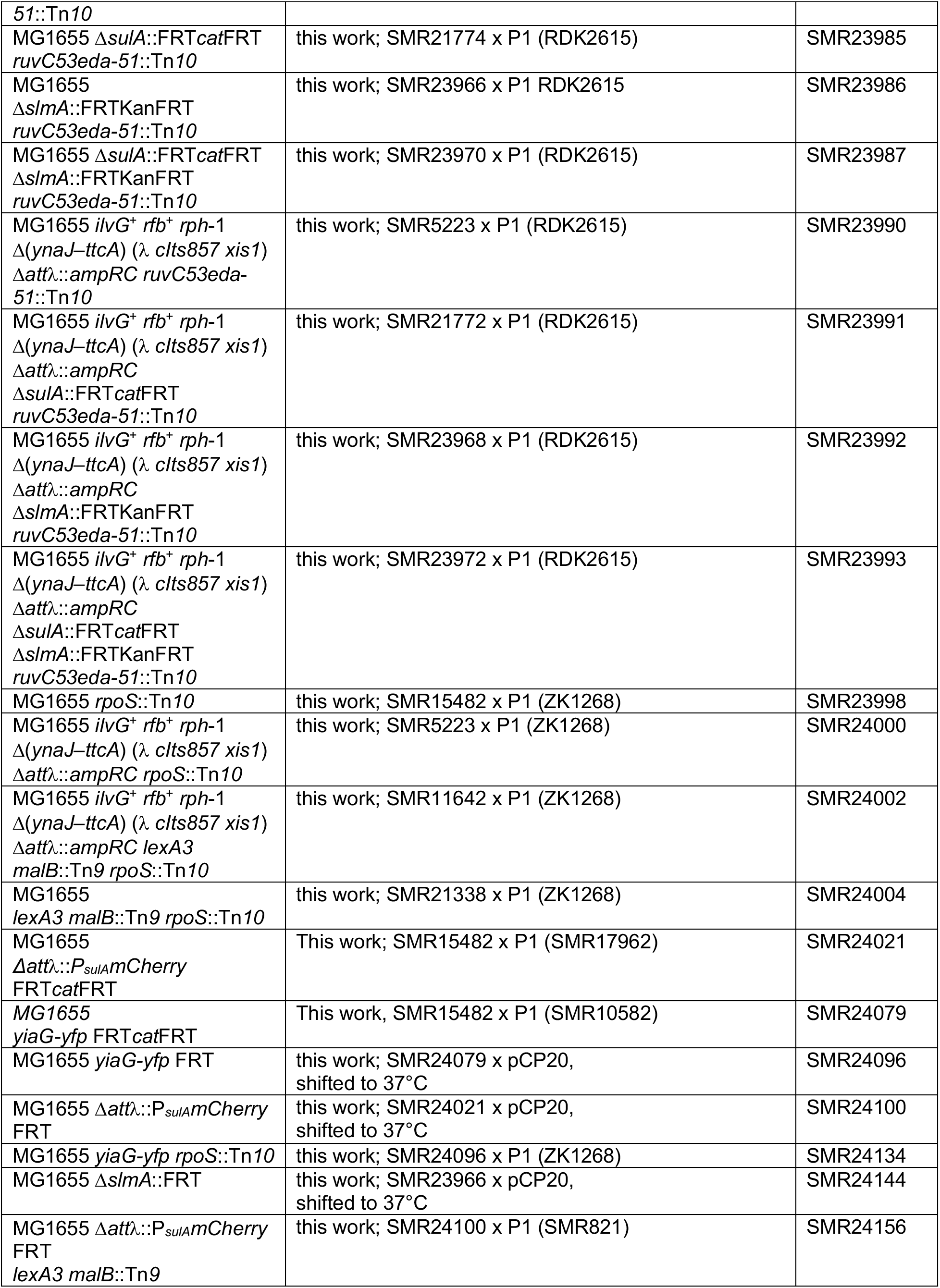
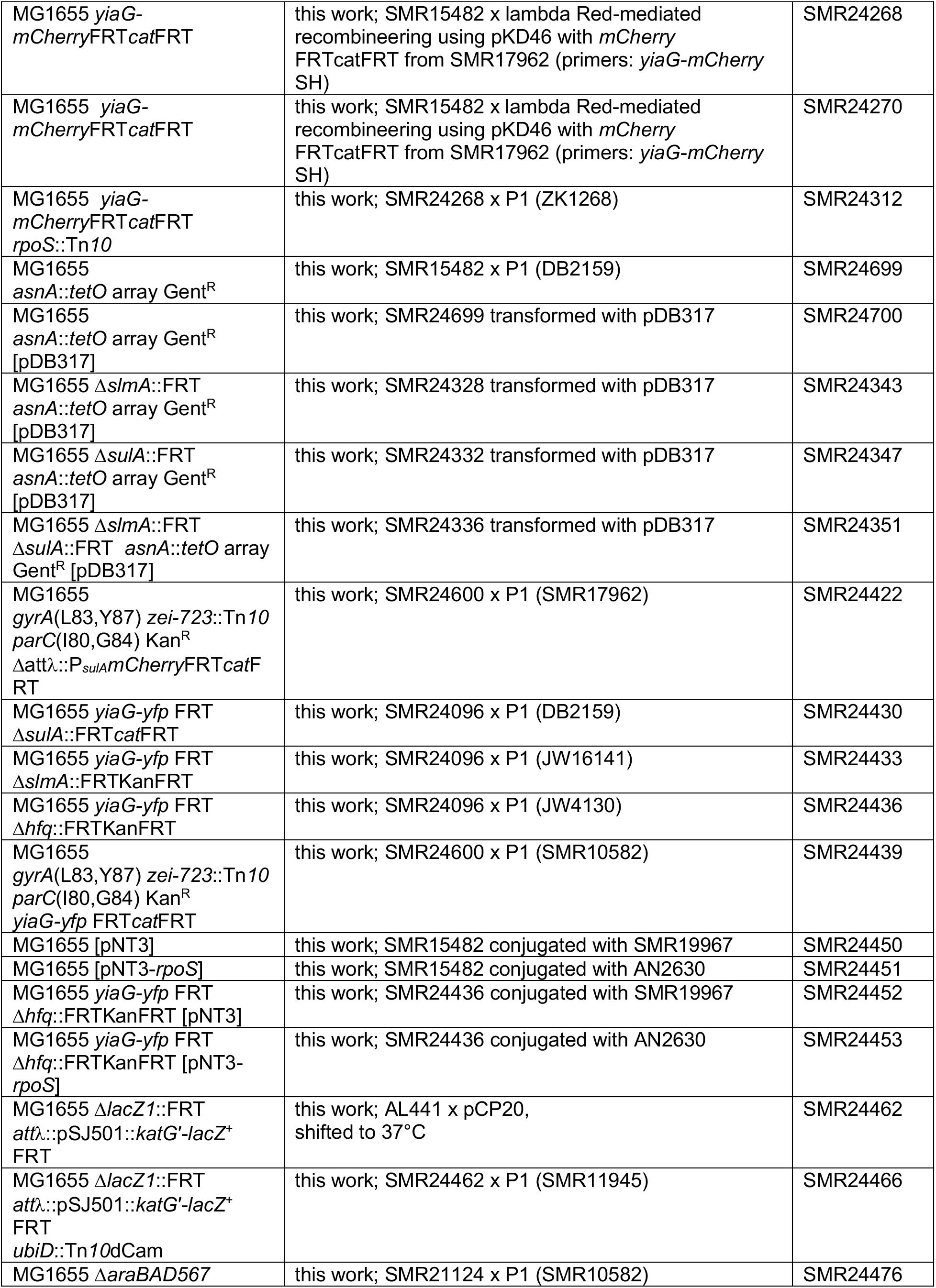
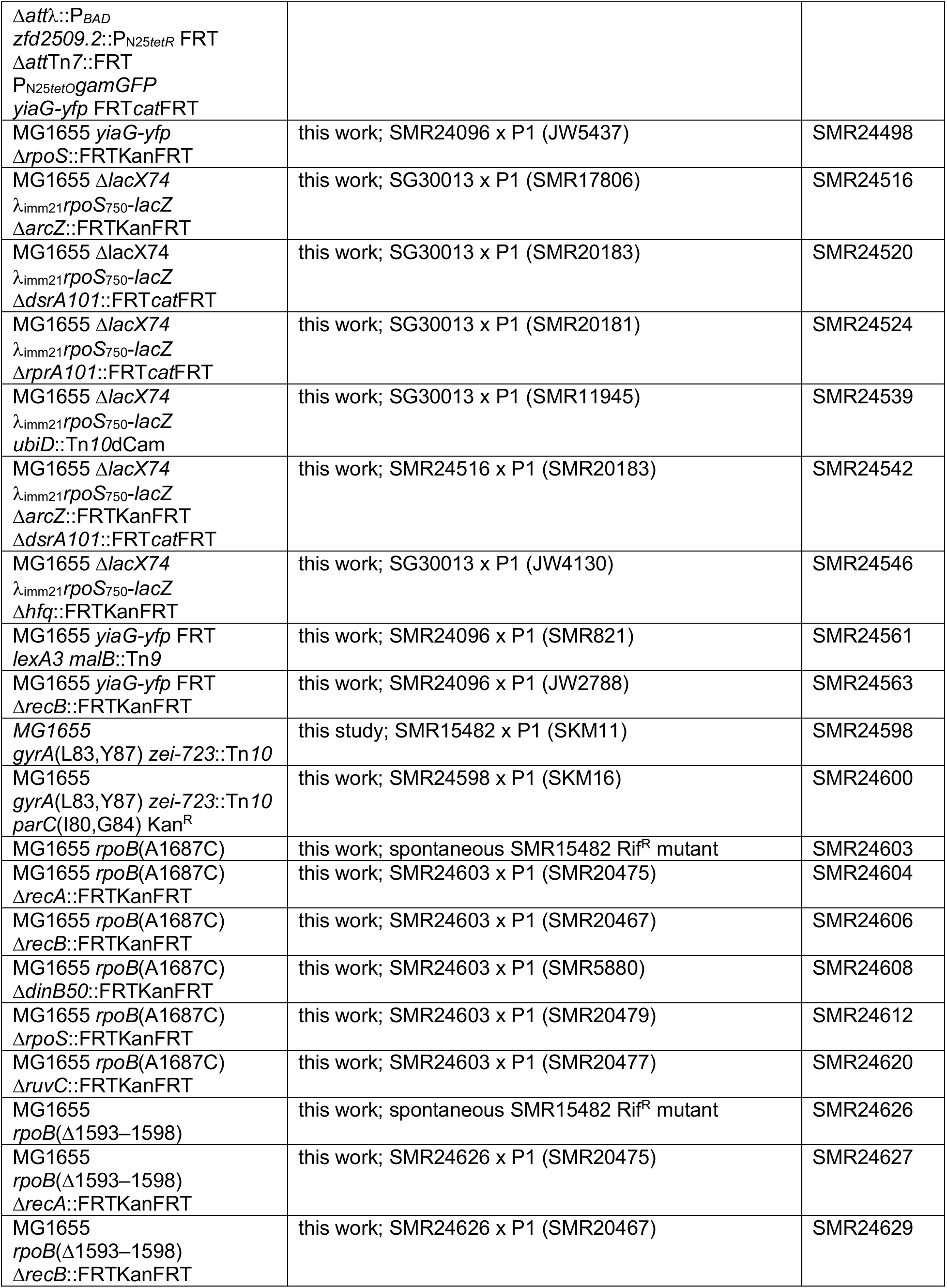
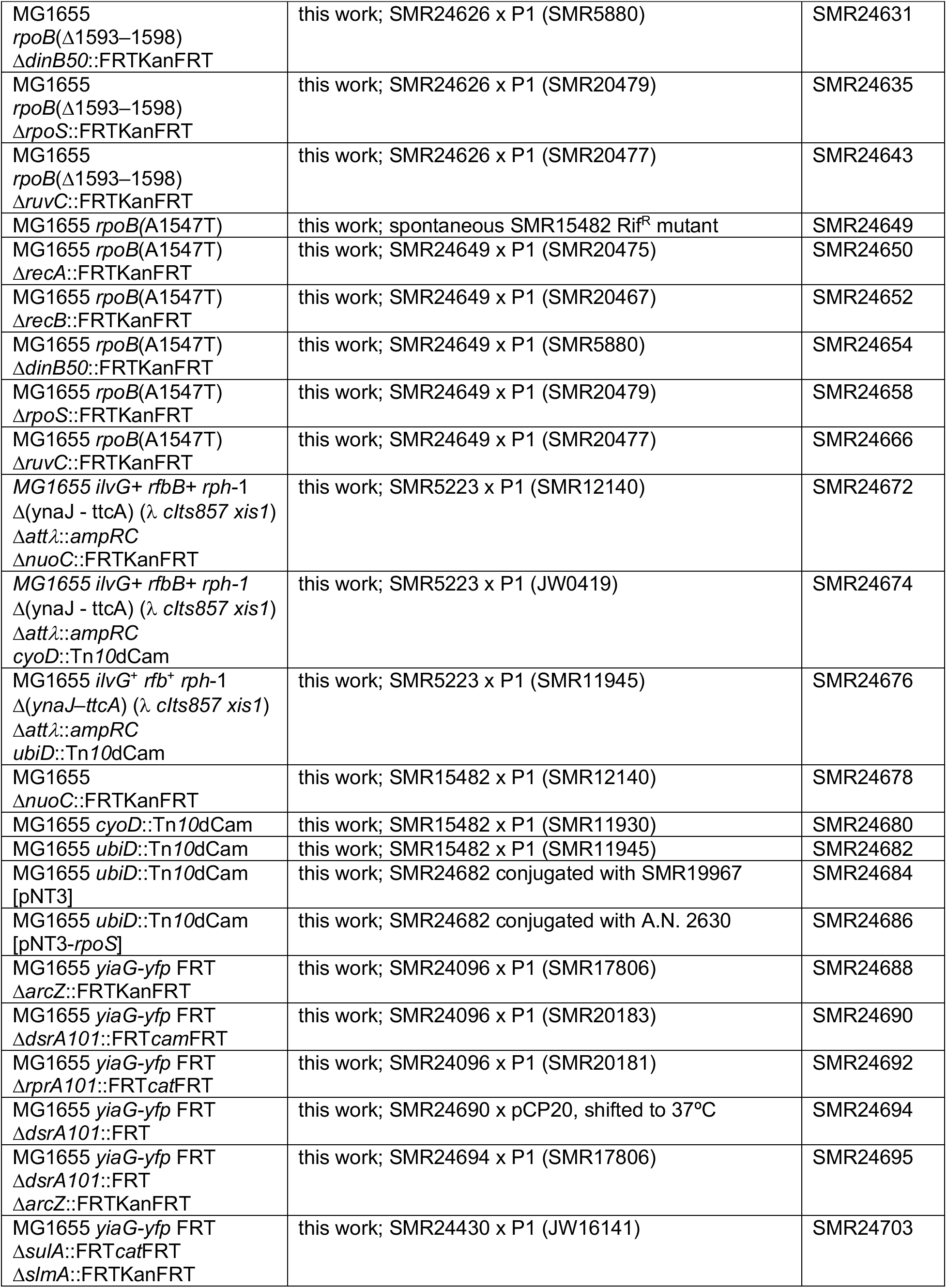
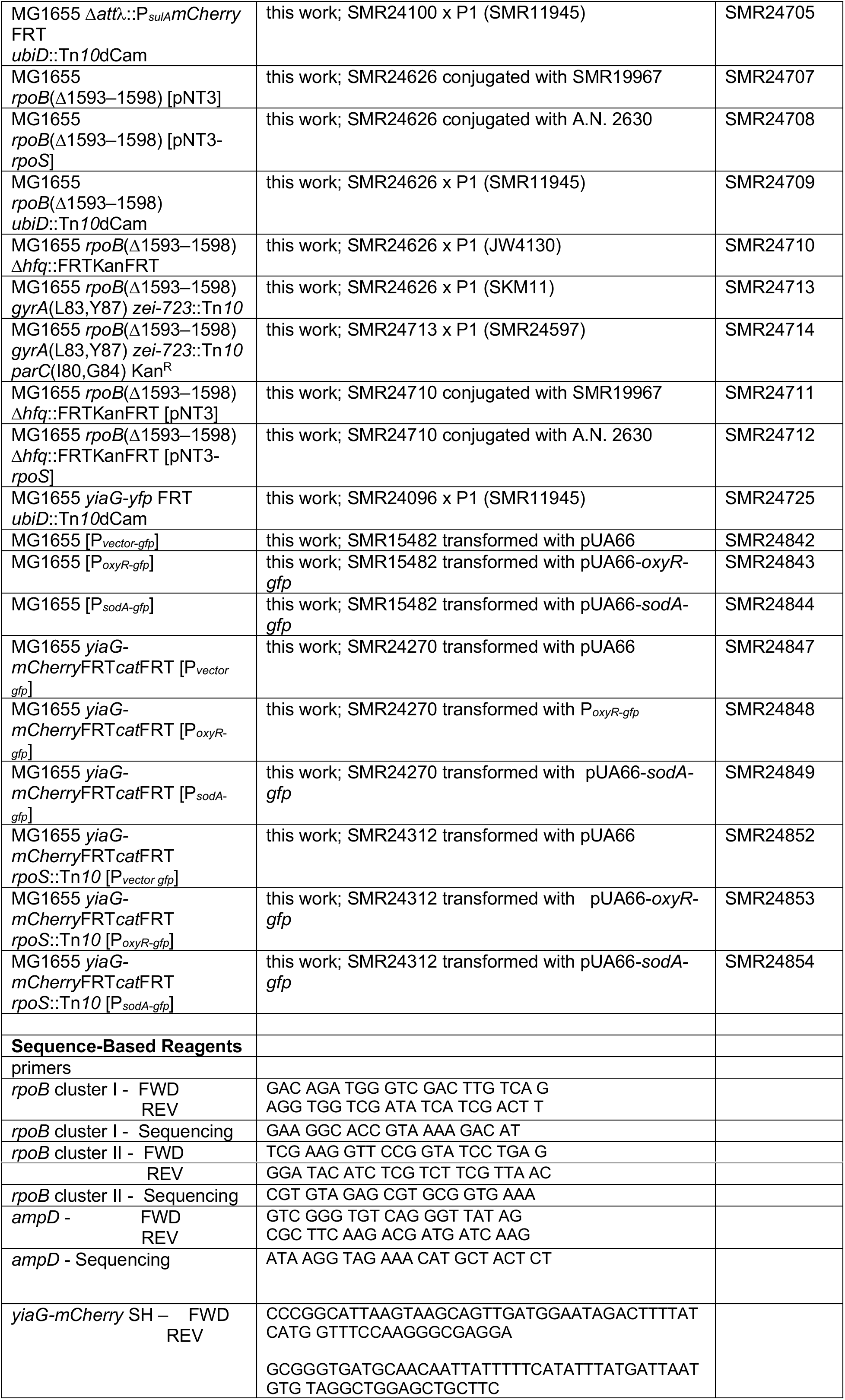

## CONTACT FOR REAGENTS AND RESOURCE SHARING

The corresponding author, Susan M. Rosenberg, is the contact for reagents and resource sharing.

## EXPERIMENTAL MODEL AND SUBJECT DETAILS

*Escherichia coli* (strain MG1655) and isogenic derivatives were used for all experiments.

## METHODS DETAILS

### Strains, Media, and Growth

The Key Resources Table lists strains used in this study. Bacteria were grown in LBH rich medium (Torkelson et al., 1997) at 37°C with aeration, and additives where indicated at the following concentrations: ciprofloxacin (cipro, 1–24 ng/mL, Table S1), ampicillin (100 μg/ml), chloramphenicol (25 μg/ml), kanamycin (50 μg/ml), tetracycline (10 μg/ml), rifampicin (110 μg/ml), and sodium citrate (20 mM).

### Assays for Ciprofloxacin-induced Mutagenesis

Saturated overnight LBH cultures, started each from a single colony, were diluted 1:4×10^6^ into 25 ml in a 250ml flask in fresh LBH broth and incubated at 37°C with shaking for 3–3.5 h, then diluted 1:3 into fresh LBH broth (“no-cipro” controls) or into LBH with cipro at a final “sub-inhibitory” concentration minimal antibiotic concentration (MAC) that caused a final cfu titer of 10% of the titer observed in the no-cipro control, by the final 24h or 48h time point (fluctuation tests, below). This concentration was determined individually for each experimental strain. For dose-response fluctuation tests, the final cipro concentrations were 1, 2, 4, 8.5, and 12 ng/ml.

For all fluctuation tests, between 10 and 60 1-ml aliquots of cultures diluted 1:3 were dispensed into 96-deep-well plates or 14-ml tubes and incubated at 37°C with shaking. After 24h (RifR) or 48h incubation (AmpR), samples were plated onto LBH agar for determination of total viable cell titers or selective LBH-agar plates containing rifampicin (110 μg/ml) or ampicillin (100 μg/ml) to select mutants resistant to each drug. Total and resistant cfu were counted, and mutation rates (mutations per cell per generation) estimated with the MSS-MLE algorithm using the FALCOR calculator (Hall et al., 2009). The fold change in cipro-induced mutagenesis for each strain was determined as the ratio of the mutation rates of the treated divided by the untreated control samples.

For fluctuation tests performed with addition of reagents that reduce reactive oxygen, the final concentrations were 100 mM for thiourea, 0.25 mM for 2,2’-bipyridine, and 100 μM for edaravone. For assays in which GamGFP was produced to trap double-strand breaks (DSBs) (Shee et al., 2013), GamGFP was induced from the chromosome using 10 and 20 ng/mL doxycycline in LBH liquid or in plates, as used for determining cfu/ml. Plasmids for σ^S^ artificial upregulation and the empty-vector control were obtained from the mobile plasmid collection (Saka et al., 2005), and were induced with 30 μM isopropyl β-D-1-thiogalactopyranoside (IPTG) present throughout growth, except in the plates used to determine RifR or total cfu/ml. σ^S^ production was confirmed by western blotting.

### Reconstruction Experiments

Reconstruction experiments were performed to verify that differences in cipro-induced mutant cfu titers observed between wild-type and various mutant strains were not caused by differences in colony-formation efficiency under exact reconstructions of selection conditions: selective plates with varying amounts of isogenic sensitive neighbor cells (10^8^ or 10^9^). About 100 cfu of ampicillin-resistant *ampRC* Δ*ampD* cells or rifampicin-resistant *rpoB* Δ1687C, *rpoB* Δ1593-1598, *rpoB* A1547T mutant cells of each experimental strain genotype were mixed with ~10^9^, ~10^8^ or ~10^7^ isogenic sensitive neighbor cells and plated onto ampicillin or rifampicin selective plates, respectively, and their numbers and speed of forming colonies scored. These platings reconstruct the experimental conditions in which mutant cells form colonies scored in our **Assays for Ciprofloxacin-induced Mutagenesis**. Resistant mutants were also plated alone for reference. For each strain, we quantified cfu observed after 24 h (ampicillin) or 48 h (rifampicin) at 37°C. Two replicates for each culture condition were performed per strain. See figure legends for numbers of independent experiments.

### Competition Experiments

Cultures of sensitive and resistant mutants of each experimental strain genotype were mixed at a 50:50 ratio and grown per fluctuation tests, then plated at the end of the growth period on selective rifampicin or ampicillin medium and non-selectively, to obtain the final ratios of sensitive and resistant cfu after growth in competition. Pure cultures were also established as controls. These experiments showed that neither RifR not AmpR mutants is selected (wins the competition ending over 50% of cfu), and both are actually significantly counter-selected relative to their sensitive parent strains (e.g., Figure 1C and legend). These data indicate that all of our estimates of the induction of mutagenesis to RifR and AmpR are underestimates. See figure legends for number of independent experiments.

### Flow Cytometric Assays for σ^S^- and SOS-Response-Regulated Promoter Activity

Quantifications of cells that have induced their σ^S^ or SOS responses, and how much they have, were achieved using engineered chromosomal fluorescence reporter genes and flow cytometry, per (Nehring et al., 2015; Pennington and Rosenberg, 2007) for SOS, and per (Al Mamun et al., 2012) for σ^S^-response activation. We used the *yiaG-yfp* σ^S^-response reporter (Al Mamun et al., 2012) and the Δ*attλ*::*P_sulA_mCherry* SOS reporter (Pennington and Rosenberg, 2007) modified by Nehring et al. (Nehring et al., 2015) in separate strains grown under fluctuation-test conditions as described for **Assays for Ciprofloxacin-induced Mutagenesis**, with or without cipro, at indicated concentration/s, and harvested the cells in late log phase or stationary phase. For quantification, flow cytometry “gates” were calibrated, for SOS, using the negative-control SOS-off lexA(Ind^-^), and SOS-response proficient cells, per (Pennington and Rosenberg, 2007) as the dividing place between peaks of the bimodal distribution of SOS-proficient cells at which most cells, diverge from the spontaneously SOS-induced fluorescent cell subpopulation, usually at between 0.5% and 1% of cells for cells cultured in LBH broth. Essentially all SOS-non-inducible *recA* or lexAInd^-^ cells fall below this gate (~10^-4^ of them cross the gate). Cells that fell below this gate (less fluorescence) were scored as SOS negative, and above the gate as SOS positive. For the σ^S^ response, gates for σ^S^-high activity cells were set to the point at which fewer than 0.5% of cells with cipro but without the reporter gene were positive. At this gate fewer than 10^-3^ of Δ*rpoS* cells, which are deficient in σ^S^-response induction, cross the gate and would be scored as positive. For all, the percent of the population that scored as positive is reported.

### Fluorescence-Activated Cell Sorting

Cell sorting was performed using a FACS Aria II cell sorter (BD Biosciences, San Jose, CA) with a 70-μm nozzle. *E. coli* cells were identified using forward and side scatter parameters, and these were sorted using sterile 1X phosphate buffered saline (PBS) as sheath fluid. After treatment with cipro, yellow fluorescent protein-positive (σ^S^ activity, *yiaG-yfp*) and non-fluorescent cells were sorted into 14 mL conical tubes (20-30×10^6^ negative cells and 3-8×10^6^ positive cells) and plated on LB agar with and without rifampicin to determine cfu/mL (per **Assays for Ciprofloxacin-induced Mutagenesis**, above). These data were used to calculate RifR mutant frequencies in the sorted σ^S^ high-activity, σ^S^ low-activity, unsorted, and mock-sorted populations, the last being cells run through the machine and all cells collected. Control sorts for cyan fluorescent protein, encoded by the *Placcfp* gene, a negative control for metabolically active cells, and mutagenesis assays, were performed similarly in parallel with the experimental sorts.

### HPII Catalase Activity

HPII (σ^S^-dependent catalase) activity was measured as described (Iwase et al., 2013). The viable cell titers (cfu/mL) of cells growing in LBH broth were determined at appropriate time points in log or stationary phase. HPI catalase was inactivated by heating 100 μL culture aliquots at 55°C for 15 min. After inactivation, 100 μL 30% H_2_O_2_ and 1% Triton-X 100 (sigma) were added. After an additional 15 min incubation, the height of bubble formation was measured in millimeters. The millimeters of bubbles were then normalized to cfu/mL of cells. Controls in Δ*rpoS* cells demonstrated that these assays report on σ^S^-response-dependent catalase activity.

### Microscopy and Quantification of GamGFP (DSB) and TetR-mCherry (Chromosome) Foci

Cells containing the chromosomal inducible GamGFP cassette were diluted 1:4×10^6^ into 250mL flasks and grown for 3 h. These were then diluted 1:3 in media with or without cipro (1-8.5 ng/ml). GamGFP, a DNA DSB-specific binding protein that traps DSBs and inhibits their repair (Shee et al., 2013), was induced in late log phase using 40 ng/mL of doxycycline. After 2 h of induction, cells were fixed with 1% paraformaldehyde and placed at 4°C until microscopy images were taken. Cells containing the inducible TetR-mCherry plasmids and the *tetO* chromosomal array were diluted 1:4×10^6^ 2.5μl into 10ml into 250mL flasks and grown for 3 h. These were then diluted 1:3 in media with or without cipro (MACs). The TetR-mCherry protein binds to the chromosomal *tetO* array labeling oriC-proximal chromosomal units as red foci, and was induced in late log-phase using 2 μM of sodium salicylate. After 4h of induction, cells were fixed with 1% paraformaldehyde and placed at 4°C until microscopy images were taken. Images were visualized with an inverted DeltaVision Core Image Restoration Microscope (GE Healthcare) with a 100X UPlan S Apochromat (numerical aperture, 1.4) objective lens (Olympus) and a CoolSNAP HQ2 camera (Photometries). Captured images for analysis were chosen randomly. The images were taken with Z-stacks (0.15-μm intervals) and then deconvoluted (DeltaVision SoftWoRx software) to visualize the whole cell for precise and accurate quantification of foci per (Xia et al., 2016; Xia et al., 2018). For each experiment, >400 cells were counted using ImageJ software (NIH) with visual inspection from each independent experiment. Only foci that overlapped with DAPI DNA stain were quantified (≥99% of all foci).

### Live Cell Deconvolution Microscopy

Cells were grown as for **Assays for Ciprofloxacin-induced Mutagenesis**. At 8 hours after the addition of ciprofloxacin (8.5 ng/mL), 4 μL of culture were plated onto 35mm glass bottom cell culture plates. An agar pad containing spent medium from replicate cultures (8.5 ng/mL cipro in cells grown for 8h) was placed on top of the cells, and a glass cover slip placed over the agar pad and sealed with silicon grease to limit evaporation. Images were taken every 1-2 hours for 12 hours with an inverted DeltaVision Core Image Restoration Microscope (GE Healthcare) with a 100X UPlan S Apochromat (numerical aperture, 1.4) objective lens (Olympus) and a CoolSNAP HQ2 camera (Photometrics). Captured images for analysis were randomly chosen. The images were taken with Z-stacks (0.15-μm intervals) and then deconvoluted (DeltaVision SoftWoRx software) to visualize the whole cell. For each experiment, >250 cells were followed to track the activation of the GFP (P*sodA-gfp* oxidative stress response) and mCherry (σ^S^ activity) using ImageJ software (NIH) with visual inspection from each independent experiment.

### *rpoB* and *ampD* Sequencing

A sole RifR or AmpR colony was isolated from each of 24 cipro-treated or 24 control independent cultures and the *rpoB* or *ampD* gene sequenced, respectively. RifR *rpoB* mutations occur mostly within two mutation clusters (Reynolds, 2000), and all isolated mutants contained mutations within these two clusters. *ampD* loss of function mutations confer ampicillin resistance in our *E. coli* assay strain due to the insertion of *Enterobacter cloacae ampRC* genes in the chromosome, as previously described (Petrosino et al., 2002). The *rpoB* cluster I and II were amplified, as described (Reynolds, 2000), see also **STAR METHODS RESOURCE TABLE** for primers (Reynolds, 2000). The *ampD* gene was amplified using primers described in **STAR METHODS RESOURCE TABLE**. All PCR fragments were subjected to Sanger sequencing (GeneWIZ, Massachusetts) to identify insertions, deletions, and/or base substitutions in the *ampD* or *rpoB* genes.

### Western Blot Analyses of σ^S^ Protein Levels

Western blots for quantification of σ^S^ protein levels in cultures were performed as described (Barreto et al., 2016). Proteins were separated by SDS-PAGE and transferred to 200 polyvinylidine (PVDF) membranes (Amersham Biosciences), blocked with 2% blocking buffer, and probed with polyclonal mouse anti-aS antibody (1:700 dilution) (Neoclone) (85). Goat antimouse antibody conjugated to Cy5 fluorescent dye (1:5000 dilution) (Amersham Biosciences) was used to detect the antibody-bound σ^S^ protein. Fluorescence was quantified using a Typhoon scanner, with a PMT of 500 and 670BP 30Cy5emission filter, and the bands were quantified using ImageJ software (NIH). Quantifications from three separate western blots for σ^S^ are reported, each with band intensities normalized to the values from isogenic wild-type cells with no cipro treatment run in parallel, and the means ±SEM shown.

### Beta-galactosidase Assays

Cells were grown as for **Assays for Ciprofloxacin-induced Mutagenesis** to equivalent ODs and frozen at −20°C until assays were carried out. Determination of the β-galactosidase activity of the *P_arcZ_-lacZ, P_dsrA_-lacZ, rpoS-lacZ*, and *katG-lacZ* fusions was accomplished using the standard assay described by Miller (Miller, 1992), except that the assays were carried out in 96-well plates to ease sample processing.

### Flow Cytometric Detection of Intracellular ROS or GFP and σ^S^ Activity in Single Cells

Cells were grown in the absence or presence of cipro MAC (8.5 ng/mL) to early-, late-log, and stationary phase as for **Assays for Ciprofloxacin-induced Mutagenesis** (above). The ROS measurement protocol was modified from Gutierrez et al. and Xia et al. (Gutierrez et al., 2013; Xia et al., 2018). Cells were incubated with ROS-staining dye DHR123 (Invitrogen) for 30 min at 37°C in PBS. After washing twice with PBS buffer, flow cytometry analyses were performed immediately. Positive gates for ROS-positive cells were set so that <0.5% of cells treated with cipro without DHR dye were positive. For experiments in which ROS or GFP and σ^S^ activity were measured, cells were grown in the absence of presence of cipro MAC (8.5ng/mL) or with 0.5mM H_2_O_2_ as for **Assays for Ciprofloxacin-induced Mutagenesis** (above), then harvested serially from cultures at 4, 8, 12, 16, 24, and 48 hours for ROS detection using dihyrdorhodamine 123 (DHR), or at 12, 16, and 24 for hours for ROS detection using transcriptional fusions of the *oxyR* and *sodA* promoters to GFP (Zaslaver et al., 2006). For ROS detection using DHR, cells containing σ^S^-activity reporter *yiaG-mCherry* were collected and ROS were detected as green fluorescence, and σ^S^ activity as red fluorescence. For ROS detection using *PoxyR-gfp* and *PsodA-gfp*, cells containing both σ^S^-activity reporter *yiaG-mCherry* and plasmids carrying the *PoxyR-gfp* or *PsodA-gfp* transcriptional fusions, or a promoterless *gfp* parental plasmid Pvector-*gfp*, were maintained with 35μg/mL kanamycin, and used to detect both GFP and red fluorescence. Single color controls were also collected at time points for spectral compensation. For the *PoxyR-gfp* or *PsodA-gfp* transcriptional fusions, gates were drawn so that the promoterless-gfp vector Pvector-gfp had < 0.5% GFP-positive cells. σ^S^ high-activity-cell gates were drawn so that spontaneous σ^S^ activation in non-cipro-treated cells after growth (<0.5% of cells without cipro) were positive, and wild-type cells without the chromosomal σ^S^-response reporter (autofluorescence) had fewer than 0.5% of their cipro treated cells scored as positive.

### Mathematical Modeling of Cipro-Induced Multi-chromosome Cell Filaments

In our model, a population of microbes is exposed to severe external stress (e.g., antibiotics), and two strategies are available: either growing into “filament” cells, that can contain multiple DNA copies, or reproducing individually. We consider a case in which resistance to the external stress can be acquired by a single mutation, with baseline rate *μ*, and deleterious mutations occur at many other loci, with the number of deleterious mutations per replication following a Poisson distribution with average *λ*. We assume that during the external stress the basic mutation rates of all cells (both *μ* and *λ*) increase *A*-Fold, and mutation rates in filament cells are further increased *B*-fold relative to non-filament cells.

We denote by *s* and *δ* the selection coefficients against the external stress and each deleterious mutation, respectively. We denote by *I_a_* the level of adaptation to the external stress, where 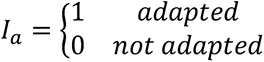. The fitness (modeled here as the probability to replicate) of an individual that possess *n* deleterious mutations is thus *ω*(*I_a_, n*) = (1 − *s*)^1−*I_a_*^ · (1 − *δ*)*^n^*. In the filament population, we assume that DNA copies in the same cell filament share gene products, and that deleterious mutations are recessive. Once a genome copy within the filament acquires the beneficial mutation that confers resistance to the major stress, it buds out of the filament, and begins to duplicate regularly (in proportion to the number of deleterious mutations it possesses).

We follow the two strategies for *k* replication cycles, starting from a population that doesn’t carry any deleterious mutations nor is adapted to the external stress. In the filament population the cells duplicate their genome without dividing and have up to 2^C^ DNA copies. Because the populations begin without any deleterious mutations, we neglect filaments in which all DNA copies share the same deleterious mutation. Therefore, the fitness of DNA copies in the filament population is affected only by the external stress, while in the non-filament population the fitness of each DNA copy (or cell) is affected both by the external stress and by the number of deleterious mutations it carries. After *k* replication cycles the filaments divide to cells, each containing a single DNA copy. We then compare the population size and fitness, the proportion of adapted individuals, and the distribution of deleterious mutations, between the filament population and the nonfilament population.

Parameter values in figure 7H: *λ* = 0.003, *μ* = 6 · 10^−7^, *δ* = 0.03, *A* = 100, = 4. In the left and middle panels we use *B* = 4 and *s* = 0.9, whereas in the right panel *B* is the value on the x-axis. The value *B = 4* is derived from empirical results presented in Figure 7G, in which we see that during antibiotic stress the mutation rate of cells that do filament (WT) have a fold-increase of ~4 relative to non-filamented cells.

The model tests the effect of filaments on evolvability, where mutation serves as the variation mechanism. However, if chromosomes in filaments also experience recombination, then the system corresponds to the case of Fitness-Associate Recombination (FAR) (Hadany and Beker, 2003b) - the less fit chromosomes experience higher recombination rate then the fitter ones. Previous work has shown that this mode of recombination results in increased mean fitness and improved adaptability (Hadany and Beker, 2003a).

Parameters:

Beneficial mutation rate *~Ber*(*μ*)
Deleterious mutation rate *~Poisson*(*X*)
*A* – stress-induced increase in mutation rate
*B* – filament cells fold increase in mutation rate relative to non-filament cells
*s* – selection coefficient of the antibiotic
*δ* - selection coefficient of each deleterious mutation (multiplicative model)
*k* – number of replication cycles

### Measurement of High-Dose Cipro Antibiotic Activity

Cells were grown to log phase OD600 ~0.5, then cipro (1.5 μg/mL) with or without edaravone (100μM) was added, and cells were harvested 0.75, 1.25, 2.25, and 3 hours later to determine cfu/mL. Cells were washed twice with PBS and then assayed for viable cfu.

### Nalidixic-Acid Test for Heritable Hypermutability

Tests for heritable mutator phenotype were as described (Torkelson et al., 1997). Ten independent (different mutations in *rpoB*) cipro-induced RifR mutant isolates were grown in parallel with control wild-type (non-mutator) and *mutS* mismatch repair-defective (mutator) strains each in duplicate independent cultures. 100μL of each saturated overnight culture was spread onto an LBH agar plate. After 10 minutes, dry nalidixic acid powder was spotted onto each plate using a capillary tube. The plates were incubated for 24 hours at 37°C, after which the number of microcolonies in the zones of inhibition were counted, and compared with the positive (*mutS*) and negative (isogenic wild-type) controls.

### Flow-Cytometric Detection of Dead Cells

Cells were grown in the presence of cipro MAC per **Assays for Ciprofloxacin-induced Mutagenesis** (above), and harvested serially from cultures at log phase (4 and 12 hours) and stationary phase (24 hours) for dead cell detection using SYTOX blue dead cell stain. Cells were stained according to manufactures recommendation. Cells were incubated with SYTOX blue dye (1μM) for 30 minutes at room temperature and flow cytometry analyses were performed immediately. As a positive control, cells were incubated in 95% ethanol for 10 minutes before staining. Positive gates for dead cells were set so that <0.2% of undyed cipro-treated cells were positive, at which 90% ± 5% of the SYTOX-blue dyed positive-control ethanol-treated cells were positive.

### Statistics

All statistics were performed in Microsoft Excel or GraphPad PRISM. For comparisons of two groups, a two-tailed Students *t*-test was used if data were normally distributed and homoscedastic. For comparisons of 3 or more groups, ANOVA with Tukey post-hoc test was used if data were normally distributed and homoscedastic, otherwise a Kruskal-Wallis non-parametric test was used. For mutation rates and ratios, which are not normally distributed, natural-logarithm transformed data were used to calculate 95% confidence intervals as well as performing ANOVAs.

## GRAPHICAL ABSTRACT

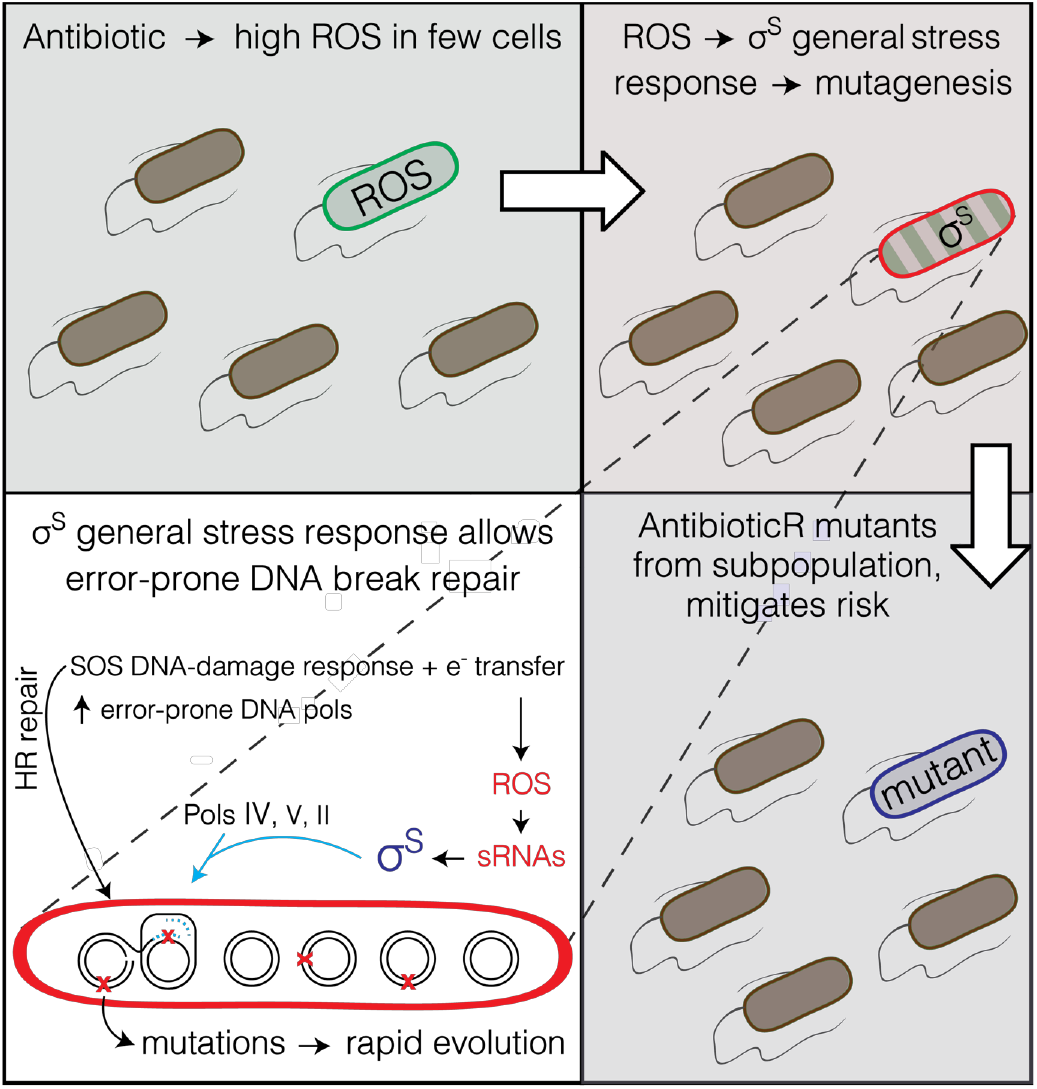

### In Brief

Bacteria exposed to antibiotic transiently differentiate a small subpopulation of gambler cells that increase mutation rate and evolve resistance, while most cells avoid the risk. The gamblers are differentiated beginning with the antibiotic inducing reactive oxygen only in subpopulation cells. The reactive oxygen activates the general stress response, which allows mutagenic DNA break repair in the gambler cells. Multi-chromosome cells are required and, modeling shows, can allow high mutation rates and rapid evolution by chromosome cooperation buffering deleterious mutations.

### Highlights

- Antibiotic-induced mutable cell subpopulation generates resistant mutants
- Mitigates risk to most cells; reactive oxygen ➔ σ^S^ stress response ➔ gamblers
- FDA-approved drug blocks σ^S^ response and mutagenesis: anti-evolvability drug
- Multiple chromosomes needed: chromosome cooperation can allow rapid adaptation

## SUPPLEMENTAL INFORMATION

Supplemental information includes a discussion file, seven figures and two tables that can be found with the article online at ***

### Supplemental Discussion S1 Controls for the FACS-sorted σ^S^-high Cell Subpopulation

We verified that FACS sorted fluorescent σ^S^-reporter carrying cells had high σ^S^ activity by showing that they displayed, first, significantly higher levels of σ^S^-dependent catalase activity (Figure 3B), and, second, more σ^S^ protein accumulation (Figure S7A) than the cells in the low-fluorescence or mock-sorted populations. We also showed by microscopic analyses that the σ^S^-response-high and -low cells did not differ detectably in cell lengths or sizes (Figure S6D,E) and that σ^S^-response-high and -low cells have no difference in the numbers of dead or dying cells (Figure S7B), indicating that their mutant frequencies can be taken at face value.

### Supplemental Discussion S2 Controls for Appearance of ROS-high Subpopulation Before σ^S^-high Subpopulation

In Figure 4A, the ROS-high cell subpopulation is apparent hours before the σ^S^-high cell subpopulation, with ROS detected by DHR dye and σ^S^ activity by mCherry fluorescence from a gene the transcription of which requires σ^S^. We can be sure that the appearance of ROS before σ^S^ activity is not the result of the lag between induction of transcription and appearance of a translated fluorescent protein because the same result is obtained when ROS and σ^S^ activity are both detected by fluorescent reports each of which requires transcription and translation Figure 4B. Additionally the lag between induction and appearance of flow-cytometry-detectable fluorescent protein is under 15 minutes (Pennington and Rosenberg, 2007), much less than the lag between ROS-high and σ^S^-high cells (Figure 4).

### Supplemental Discussion S3 Peroxide Control for OxyR and SodA Stress-response Activation by Cipro

Paradoxically, oxidative stress (H_2_0_2_) is known to inhibit σ^S^ activation through activation of *oxyS* sRNA (Zhang et al., 1998). We find peroxide activates both SodA and OxyR reporters, but peroxide alone is not sufficient for activation of the σ^S^ high-activity population (Figure 4B, right panel), indicating that something in addition to H_2_0_2_ is necessary for σ^S^-response activation. There might be an additional signal induced by cipro that allows σ^S^-response activation, or exogenous ROS in the form of H202 might not substitute adequately for ROS induced endogenously by cipro.

**Figure S1.**
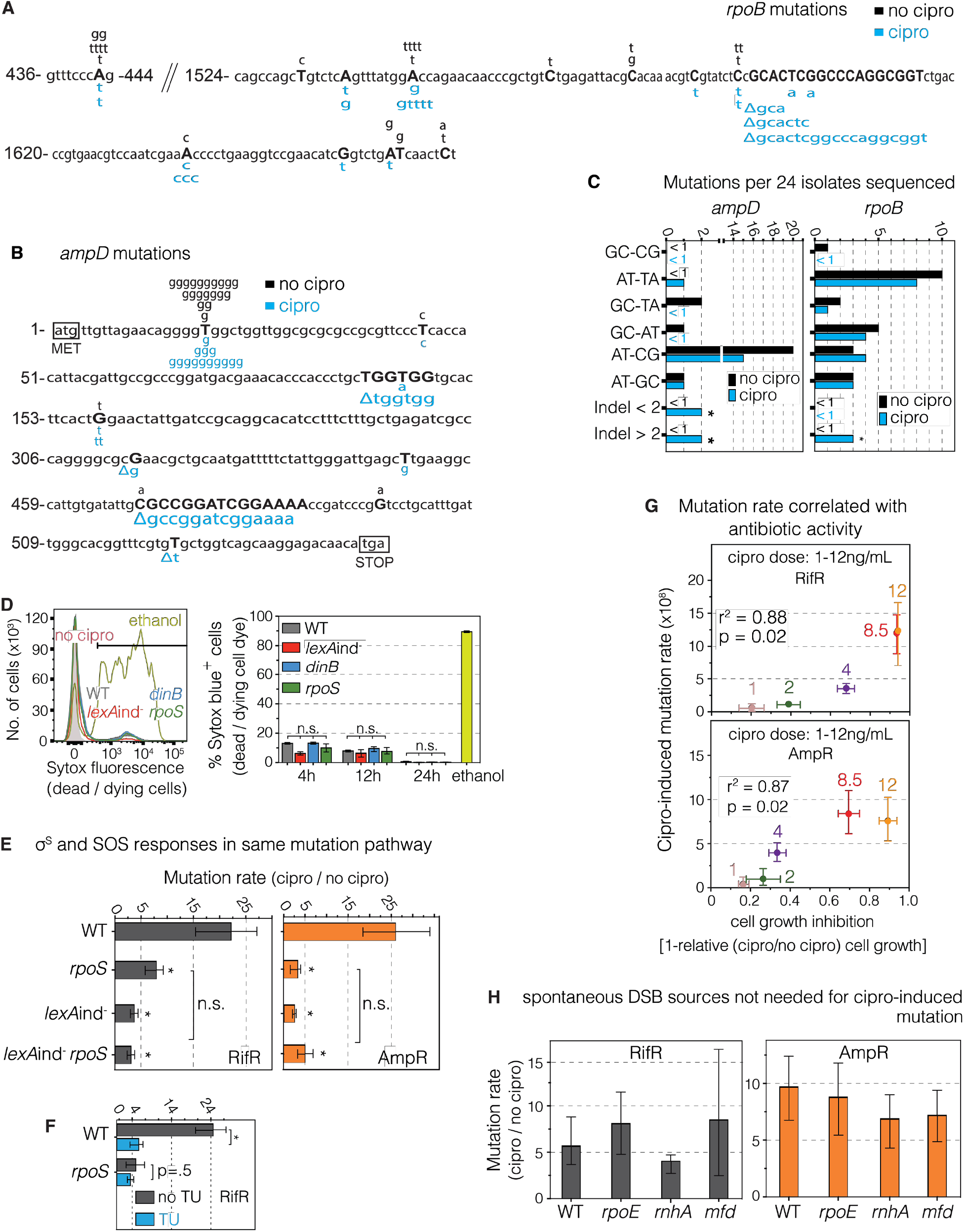
Mutation Sequences, Same Pathway (Epistasis), Independence from Spontaneous DSB Generators, and Correlation with Antibiotic Activity of Mutagenesis (Figures 1 and 2) (A) RifR cfu carry *rpoB* base-substitution mutations found in two clusters. These cause aminoacid changes that reduce rifampicin binding to the RpoB RNA polymerase subunit (Reynolds, 2000), seen also here. Black, spontaneous; blue cipro-induced mutations. (B) AmpR cfu carry *ampD* mutations. Summary of *ampD* mutation sequences from independent AmpR clones, isolated from cipro-induced and spontaneous mutants. Loss-of-function mutations in *ampD* confer AmpR to *E. coli* strain carrying a chromosomal cassette of Enterobacter *ampRC* genes (Petrosino et al., 2002) by allowing constitutive expression of the AmpC beta-lactamase, which confers resistance. Black, spontaneous, blue; cipro-induced mutations. (C) Indels are more abundant in cipro-promoted than spontaneous mutations, \**p*<0.001, Chi-squared test. Sequences from 24 independent isolates grown in the absence or presence of cipro MAC. There were significantly fewer 8-oxo-dG-signature mutations (G·C➝T·A and A·T➝C·G) in cells grown with cipro compared with no cipro, *p=0.01, two-tailed Student’s *t*-test. Counts of 24 independent RifR and AmpR isolates; *. σ1 indicates zero mutations of the type indicated among the 24 isolates, i.e., σ 1 per 24 is σ 4%. (D) SYTOX blue (Life Technologies) detection of dead/dying cells show that cell death rates do not artificially inflate cipro-induced mutation rates in wild-type above MBR-mutant strains. SYTOX-blue-detectable cell death does not differ between the wild-type strain and its MBR-defective derivatives that are defective for the SOS response (*lexA*Ind^-^), the g–^s^ response (*rpoS*), or the major error-prone DNA polymerase (*dinB*). Thus the concern of Frenoy and Bonhoeffer (Frenoy and Bonhoeffer, 2018) that bacterial cell death might cause overestimation of apparent antibiotic-induced mutation rates, predicted by their mathematical modeling, cannot account for the higher mutation rate in wild-type than MBR-mutant strains (Figure 1F). Moreover, it cannot account for the difference between the large σ^S^ low-activity (non-mutagenic) cell subpopulation and the small σ^S^ high-activity (mutagenic gambler) cell subpopulation, which show similar levels of cell death (Figure S7B). Furthermore, the mathematical modeling of (Frenoy and Bonhoeffer, 2018) showed no such potential inflation of mutation rate in the case of either—(i) a cell subpopulation producing most mutants; or (ii) multi-chromosome cells (Frenoy and Bonhoeffer, 2018), both of which we show are true for cipro-induced cross-resistance mutagenesis (Figure 4A-C and Figure 7A, respectively). (E) The SOS and general σ^S^ stress responses are epistatic for mutagenesis, i.e. act in the same mutation pathway. Cipro-induced mutation rates per Figure 1 and Methods. Means ± 95% confidence intervals of n≥ 3 independent experiments. *Differs from WT value, *p*<0.01, one-way ANOVA with Tukey’s post-hoc test of natural-log transformed data. (F) Thiourea does not reduce mutagenesis further in *ΔrpoS* cells, which lack σ^s^. The data imply that ROS promote mutagenesis in the σ^S^-response-dependent mutation pathway. (G) Correlation of antibiotic and mutation-promotion activities of cipro. Because the antibiotic activity of cipro results from DSB-generation, these data imply that cipro-provoked DSBs also drive the mutagenesis. RifR and AmpR mutation rates were assayed and estimated for different doses of cipro (1, 2, 4, 8, and 12 ng/mL). Pearson correlation coefficient. (H) Known spontaneous DSB-promoting proteins required for MBR in starvation-stressed cells are not required for MBR induced by cipro. The σ^E^ (RpoE) membrane stress response (Gibson et al., 2010) and RNA-DNA hybrids (Wimberly et al., 2013) promote DSBs at some loci in starvation-stress-induced MBR. RNA-DNA-hybrid removal by RNase HI (*rnhA*), and prevention by loss of Mfd (which dislodges stalled RNA polymerases) promote DSBs and underlie about half of mutagenesis in starvation-induced MBR (Wimberly et al., 2013), but neither is required for MBR instigated by cipro, supporting the hypothesis that cipro-provoked DSBs drove mutagenesis. Mutation rates estimated using the MSS-maximum likelihood method. Data and statistics per (E).

**Figure S2.**
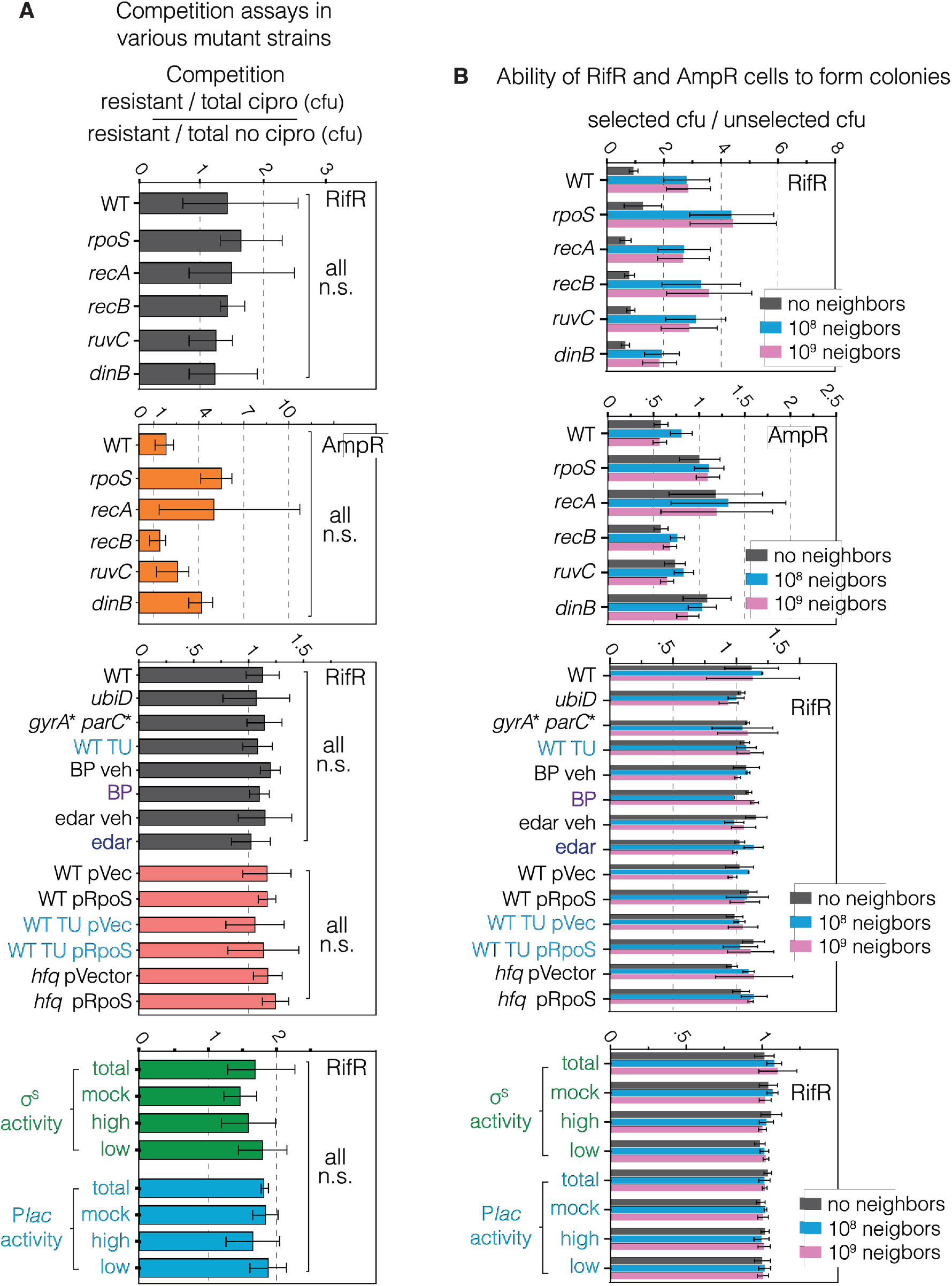
No Growth or Colony-Formation Defects in Rifampicin- and Ampicillin-Resistant Mutants (Figures 1, 2, 3, 4, 5, 6, and 7) We excluded the possibility that possible reduced growth rates of MBR-defective-mutant strains, or ROS-scavenged cells, in cipro, or as colonies on Rif and Amp plates after cipro might cause artificial apparent reductions in mutant frequencies (used to estimate mutation rates) for various mutants tested. We show no growth disadvantage in cipro of RifR or AmpR MBR-mutant or ROS-scavenged cells relative to wild-type unscavenged cells, and no defect in forming colonies afterward. (A) None of the RifR or AmpR derivatives of mutant strains assayed in this study is more disadvantaged by cipro than wild-type cells. This means that reductions in RifR or AmpR mutant cfu in these strains reflects reduced mutagenesis, not inability of the RifR or AmpR mutants to survive the assay relative to wild-type cells. Competition assays measuring percent of RifR or AmpR cells in culture after growth to saturation under conditions identical to mutation assays in the absence or presence of cipro MAC. Initial conditions were 50% sensitive and 50% resistant cells. Results shown as ratio of *%* RifR or AmpR cells grown in the presence of cipro relative to *%* of RifR cells grown in the absence of cipro. A value of 1 indicates no difference in growth. Mean and 95% CI of at least 3 independent experiments. For σ^S^ activity and *lac* activity bar graphs, data represent mean and range of 2 independent experiments. n.s., not significantly different from the wild-type value at *p*<0.01 one-way ANOVA with Tukey’s post-hoc test. (B) Reconstruction experiments show that RifR and AmpR derivatives of the various MBR- and other-mutant strains assayed form colonies under reconstructions of selective conditions as well as those in the wild-type background. These data indicate that reductions in RifR or AmpR mutant cfu in strains assayed reflect reduced mutagenesis relative to the wild-type strain background, not inability of the RifR or AmpR derivatives to form colonies under when selected. Results expressed RifR cfu titers from colonies formed on cipro divided by the total cfu assayed on LBH (non-selective) plates without cipro, and plated with or without sensitive neighbor cells. Sensitive neighbor cells are expected to be present on initial contact with the drug selection plates, but to die over time from exposure to rifampicin or ampicillin. A value of 1 indicates no deviation in the number of cfu scored in the presence of cipro from those in the absence of cipro. If greater than 1, more resistant cfu appeared under selective conditions than on no-drug plates. If less than 1, fewer mutant cells were able to form cfu on drug plates than no-drug plates. The conclusion is that reduction of RifR or AmpR cfu of the various mutants tested under selective conditions is not greater than in the wild-type, such that reductions relative to WT reflect reduced mutagenesis, not reduced mutant-cell outgrowth into a visible colony. Mean ± 95% CI of at least 3 independent experiments. For σ^S^ activity and *lac* activity bar graphs, data represent mean ± range of 2 independent experiments. None was significantly different from the wild-type, one-way ANOVA with Tukey’s post-hoc test.

**Figure S3.**
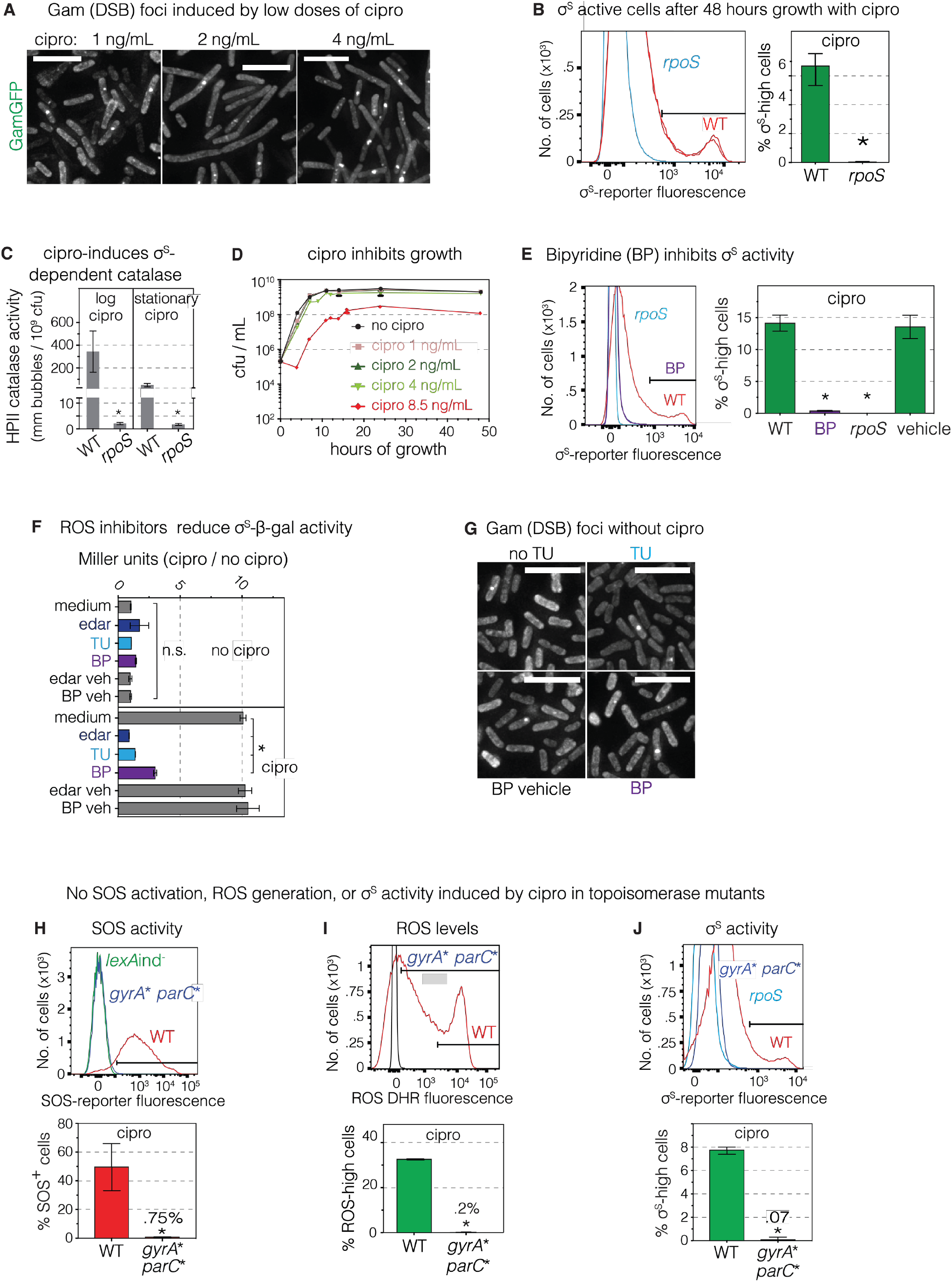
Cipro Induces GamGFP DSB Foci Dose-Dependently, SOS, ROS and σ^s^ via Binding Target Topoisomerases, and Inhibition of σ^S^- or σ^S^-*lacZ* Activity by ROS Reducers BP, TU, and Edaravone (Figures 1, 2, and 3) (A) Cipro induces GamGFP DSB foci dose dependently. Representative GamGFP foci in cells grown with 1, 2, and 4 ng/mL of ciprofloxacin. Quantification, Figure 1G. White scale bar indicates 5μm. (B) σ^S^ high-activity cells remain significantly activated after growth in cipro for 48 hours (AmpR assays conditions) detected using the σ^S^ response reporter *yiaG-yfp* (yellow fluorescence). Means ± range of 2 independent experiments. *Differs from wild-type value at *p*<0.01, two-tailed Student’s *t*-test. (C) The σ^S^-dependent HPII catalase activity (Iwase et al., 2013) is induced by cipro in both log- and stationary-phase cells. HPII measured as bubbles per 10^9^ cells. Means ± SEM of 3 independent experiments. *Differs from wild-type value at *p*<0.01, one-way ANOVA with Tukey’s post-hoc test. (D) Extents of inhibition of *E. coli* growth at various cipro concentrations. Growth curves of *E. coli* in the presence of 0, 1, 2, 4, and 8.5 ng/mL of ciprofloxacin. Where not visible, error bars are smaller than the symbol. Means ± 95% confidence intervals of 3 independent experiments. (E) ROS are required for cipro induction of σ^S^ general stress response activity. ROS-preventing agent 2,2’ bipyridyl (BP, 0.25mM) inhibits cipro induction of σ^S^ activity in flow cytometric fluorescence assay of log-phase cells carrying the *yiaG-yfp* σ^S^-response reporter. Afu, arbitrary fluorescence units. Histograms show the distribution of individual cells’ σ^S^ activity in the presence of BP. Means ± range of 2 independent experiments. *Differs from wild-type value at *p*<0.01, one-way ANOVA with Tukey’s post-hoc test. (F) ROS are required for accumulation of σ^S^ in response to cipro, assayed with the σ^S^-beta-galactosidase translational-fusion reporter in log-phase growing cells. Data represented as relative to data in untreated cells (no-cipro) in Miller units. Means ± range of 2 independent experiments. *Differs from wild-type value at *p*<0.01, one-way ANOVA with Tukey’s post-hoc test. (G) Spontaneous GamGFP foci are not the result of spontaneous ROS (control for cipro-treated cells, Figure 2F). ROS reducers thiourea (100 mM, TU) and 2,2’ bipyridine (0.25mM, BP) do not change spontaneous levels of GamGFP DSB foci in log-phase growing cells. The ferrous iron chelator 2,2’ bipyridyl (bp) inhibits ROS-forming Fenton reactions. Thiourea (TU) scavenges hydroxyl radicals. White scale bar indicates 5μm. (H-J) Cipro binding to its target type-II topoisomerases, gyrase and/or Topo IV (encoded by *parC*) required for activation of (H) the SOS response, (I) generation of ROS, and (J) activation of the σ^S^ general stress response. The *gyrA\** and *parC\** mutant alleles encode subunits of gyrase and Topo IV, respectively, that are functional but are not bound by cipro, and so are not inhibited by the drug. Representative flow cytometry histograms using the *gyrA\** S83L/D87Y *parC\** S80I/E84G. SOS, σ^S^ activity, and ROS measured by flow cytometry in strains carrying the chromosomal SOS fluorescence reporter transgene *P_sulA_mCherry*, the σ^S^-response reporter *yiaG-yfp*, or stained with the ROS specific dye dihydrorhodamine 123 (DHR). (Means ± range of 2 independent experiments. *Differs from wild-type value at *p*<0.01, two-tailed Student’s *t*-test.

**Figure S4.**
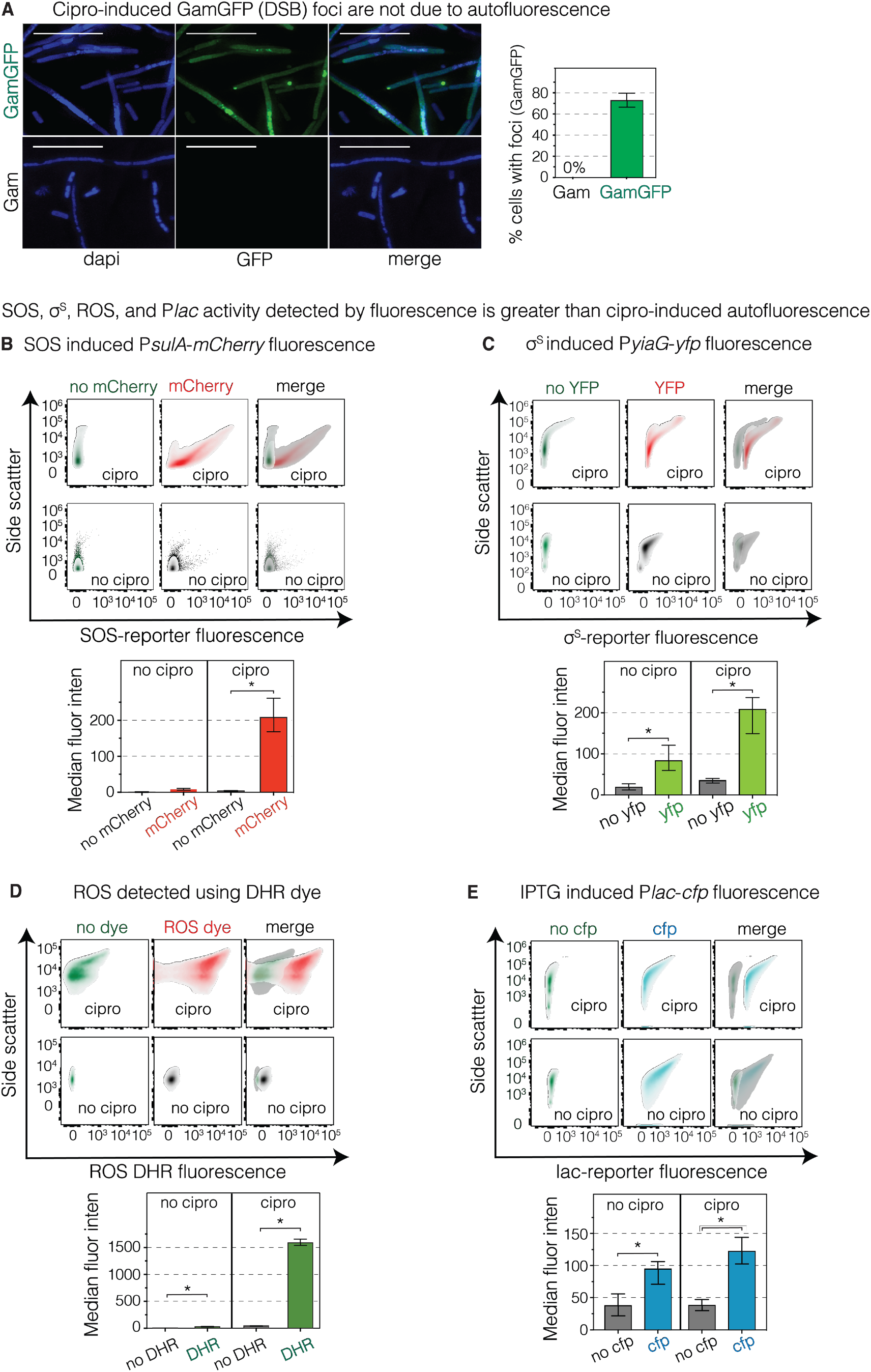
Fluorescence Data Exceed Cipro-Induced Autofluorescence (Figures 1, 2, 3, 4, 5, and 6) (A) Autofluorescence induced by cipro does not cause foci. Cells that produce Gam or GamGFP were grown in 8.5ng/mL cipro. Microscopy was performed to capture GFP fluorescence. Only cells with GamGFP turn green and form foci. At least 200 cells were quantified per experiment. (B-E) Autofluorescence does not contribute to cipro-promoted fluorescence-reporter activity detected in flow-cytometric assays. Autofluorescence has been reported in bacterial cells treated with bactericidal antibiotics (Renggli et al., 2013). To ensure that fluorescence signals are above auto-fluorescence background levels, we compared MAC cipro treatment of cells without and with the reporters of— (B) the SOS response (the Δ*attλ:.P_sulA_mCherry* transgene); (C) σ^S^ activity (*yiaG-yfp* reporter); (D) ROS, assayed using the dye dihydrorhodamine (DHR); or (E) cyan fluorescence from *Placcfp* activity induced by IPTG (1mM). Cipro-induced autofluorescence in cells without fluorescence reporters in the red, yellow, green, and cyan emission wavelengths produce less fluorescence than the positive-fluorescence readings in cells with the chromosomal fluorescence reporters, or ROS measured using DHR. The autofluorescence does not overlap with induced fluorescence signals in cells carrying the fluorescent reporters or dye. A-E, data represent mean and range of 2 independent experiments. *Different from wild-type, *p*<0.01 using a (A) two-tailed Student’s *t*-test and (B-E) one-way ANOVA with Tukey’s post-hoc test.

**Figure S5.**
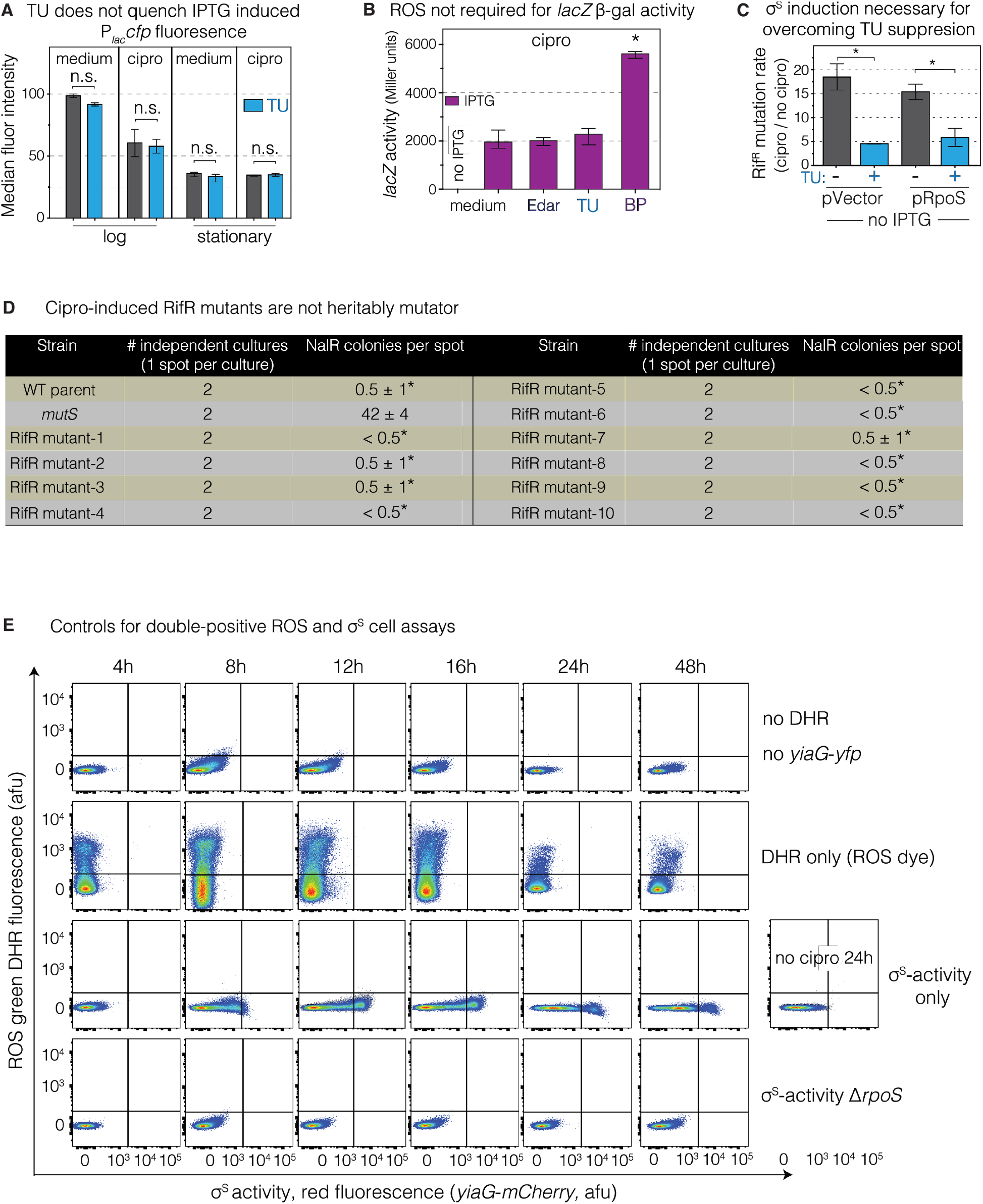
No Effect of ROS Reducers on β-gal Activity or Fluorescent-Protein Activation; IPTG Induction of σ^S^ Substitutes for ROS in Mutagenesis; and Single and No Fluorescence Controls for ROS and σ^S^ Detection in Same Cells (Figures 2, 3, 4, and 5) (A) Thiourea (TU) does not quench or inhibit the accumulation of IPTG-induced cyan fluorescent protein under the control of the *lac* promoter in either log- or stationary-phase cells. Cells with a chromosomal *P_lac_cfp* reporter were grown under conditions of the mutation assays in the presence or absence of 1 mM IPTG and TU (100 mM), and flow cytometric assays performed. Negative control for Figure 4B. Means ± range of 2 independent experiments. n.s. not significant, one-way ANOVA with Tukey’s post-hoc test. (B) ROS are not required for *lac*-reporter activity. ROS reducers TU, BP, and edaravone do not inhibit activation or activity of β-galactosidase enzyme. Cells were grown with IPTG (100mM), cipro and TU, BP, or edaravone and β-gal activity measured. Negative control for Figure 5D. Means ± range of 2 independent experiments. *Differs from no-cipro control IPTG value, *p*<0.01, one-way ANOVA with Tukey’s post-hoc test. (C) Induction of *rpoS* transcription from the engineered expression cassette is required for σ^s^ substitution for ROS in mutagenesis. TU inhibition of mutagenesis is not suppressed in cells with pRpoS plasmid if no IPTG is added. Negative control for Figure 2H. Cipro-induced RifR mutagenesis was measured in cells containing IPTG-inducible vector or pRpoS plasmid growing in the presence or absence of TU (100 mM) and no IPTG. Means ± range of 2 independent experiments. *Differs as indicated in figure, *p*<0.01, one-way ANOVA with Tukey’s post-hoc test. (D) Cipro-induced RifR mutants are not heritably mutator. Nalidixic-acid-resistance-mutagenesis assay. Two independent cultures of the wild-type parent, stable mutator (mismatch repair defective *mutS*), and 10 different cipro-induced RifR mutant isolates were spread on plates, then spotted with nalidixic acid, incubated, and NalR mutant papillae in the zones of inhibition counted. *Differs from *mutS* mutator strain, *p*<0.0001, one-way ANOVA with Tukey’s post-hoc test. These data show that the state of increased mutagenesis seen in σ^S^-active gambler cells, relative to the whole population and to the σ^S^-low main subpopulation (Figure 3A), is transient, and not a heritable mutator state. (E) No fluorescence and single-color controls for detecting both ROS (DHR dye) and σ^S^-activity (*yiaG-mCherry*) in cultures grown in the presence of ciprofloxacin. Cells were harvested for flow cytometry serially from cultures at 4, 8, 12, 16, and 24 hours after the addition of cipro. Negative controls for Figure 4A and B. Representative flow-cytometry plots from 3 experiments.

**Figure S6.**
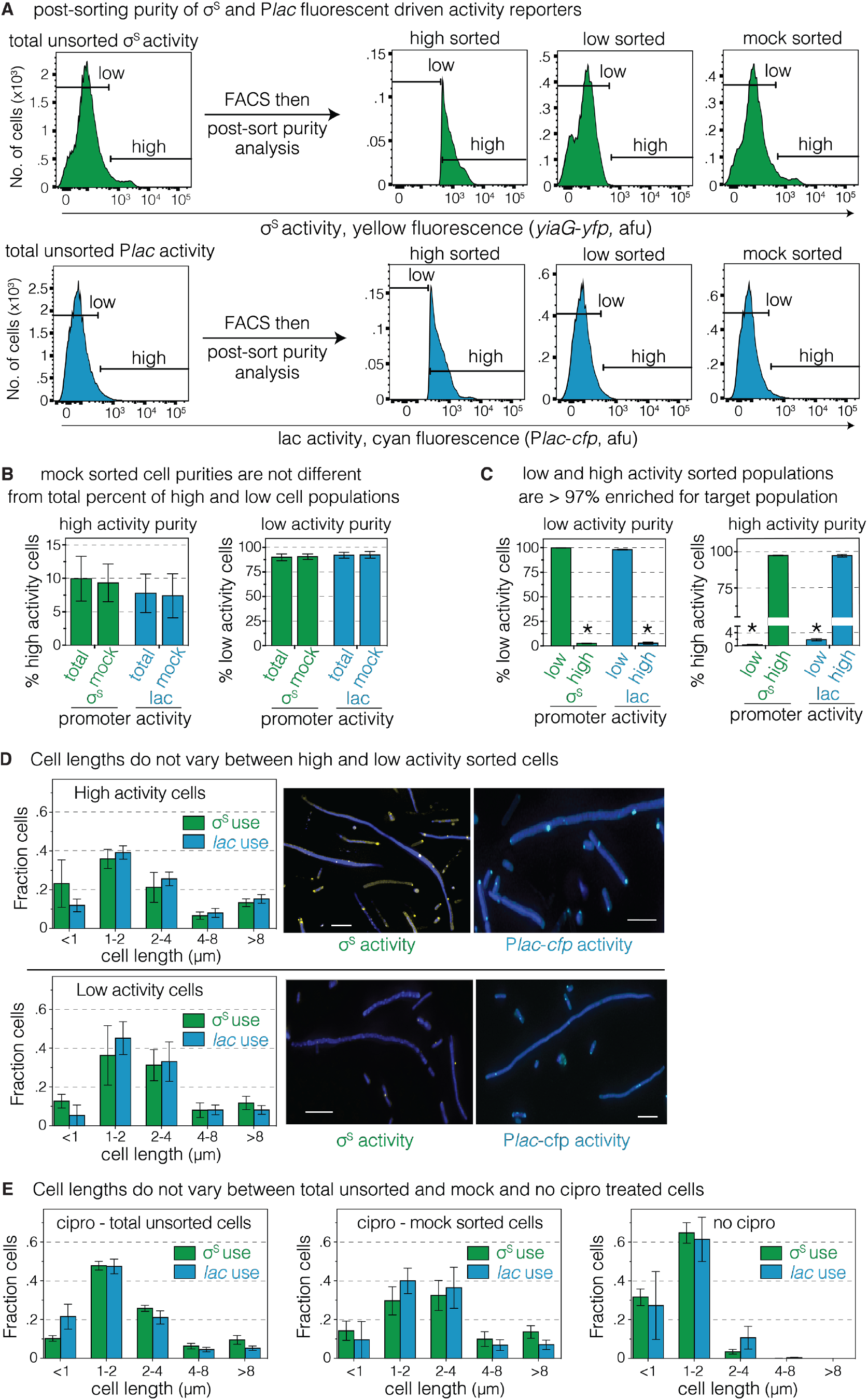
Purity of Sorted Cells and Cell Lengths in Subpopulations (Figure 3) (A) Sorted cell populations are at least 97% pure. Post-sort purity checks verify expected fluorescence intensities in cells in mock-sorted, and low- and high-fluorescence sorted cell populations. (B and C) Quantification of post-sort purity. Data represent mean and range of 2 independent experiments; * *p*<0.0001, using a one-way ANOVA with Tukey’s post-hoc test. (D and E) Cell length frequencies do not differ between *lac*-reporter and *yiaG-yjp* σ^S^-response reporter sorted populations. Mean and SD of 3 independent experiments counting > 300 cells. No differences in cell lengths were detected using a one-way ANOVA with Tukey’s post-hoc test. (D) High and low σ^S^ activity sorted cells. Representative merged images showing both DAPI DNA staining and either CFP or YFP fluorescence and quantitation of cell lengths from different populations. Scale bar represents 5μM. (E) Unsorted, mock-sorted, and untreated cell controls.

**Figure S7.**
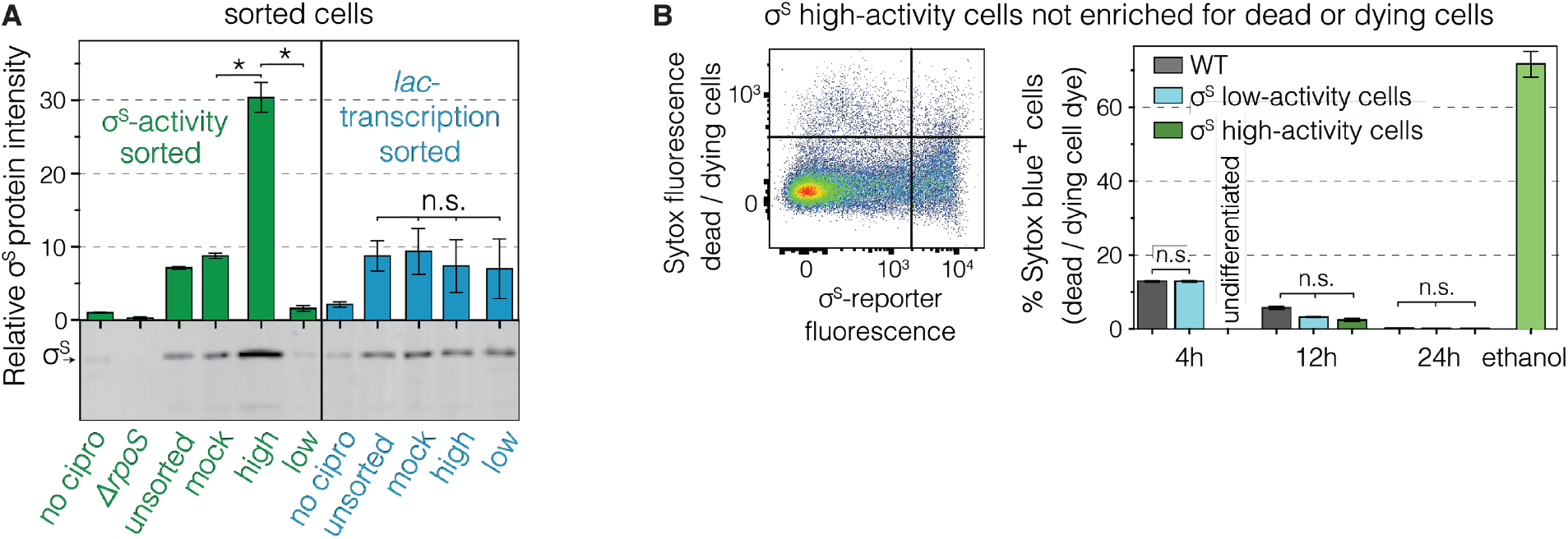
Accumulation of protein in σ^S^-high cells, and negligible dead cells in both σ^S^ high- and σ^S^ low-activity cell subpopulations (Figure 3) (A) High σ^S^ protein levels in FACS sorted σ^S^ high-activity cells. Western blots from cell subpopulations. Means ± SEM of 3 independent experiments. \**p*<0.01, one-way ANOVA with Tukey’s post-hoc test; n.s., not significant. (B) Cell death is not different in σ^S^ high- or low-activity cell subpopulations. Flow cytometry assay for cell death in log phase (4h and 12h) and stationary phase (24h) of strains with the *yiaG-mCherry* σ^S^-response reporter stained with SYTOX blue dead-cell stain, quantifies single cells with dye-permeable membranes. Representative flow cytometry distribution of live and dead cells with high and low σ^S^ activity. Means ± range of 2 independent experiments. n.s., not significant, one-way ANOVA with Tukey’s post-hoc test.

