## Supplementary material for "Gamblers: an Antibiotic-induced Evolvable Cell Subpopulation Differentiated by Reactive-oxygen-induced General Stress Response"

**Table S1. Cipro concentrations and growth inhibition, Related to Figures 1-7 and Star Methods**

| Strain  RifR  (AmpR) | Range [Cipro ng/mL]  RifR  (AmpR) | Mode [Cipro ng/mL]  RifR  (AmpR) | Relative growth  (treatment/control) x 100  RifR  (AmpR) | MIC  (ng/ml) |
| --- | --- | --- | --- | --- |
| ∆*recB* SMR20467  (SMR5228) | 0.9–1.2  (1.0) | 1.2  (1.0) | 3.4–7.6  (11.7–17.5) | 3 |
| ∆*recA* SMR20475  (SMR5226) | 1.3–1.5  (1.3–1.6) | 1.3  (1.5) | 2.2–11.9  (11.5–40.0) | 2 |
| ∆*ruvC* SMR20477  (SMR23928) | 1.6, 1.7  (1.2–1.7) | 1.6  (1.6) | 4.0–13.7  (4.0–41.7) | 3 |
| *lexA*Ind^-^ SMR21338  (SMR11642) | 2.5–3.5  (2.9–3.1) | 2.5  (2.9) | 2.7–16.3  (18.7–43.3) | 4 |
| *∆rpoS* SMR20479  (SMR11641) | 5.0–7.0  (5.0–6.3) | 5.0  (5.0) | 2.5–22.7  (26.3–79.5) | 12 |
| *∆rpoS* thiourea (100mM)  SMR24134 | 5.0  (ND) | 5.0  (ND) | 46.7–86.1  (ND) | ND |
| ∆*pol II* SMR23932  (SMR23974) | 6.0–10.0  (6.0–7.9) | 6.0  (6.0) | 1.7–15.2  (38.0–69.0) | 12 |
| ∆*mfd* SMR21946  (SMR21919) | 7.0  (7.0) | 7.0  (7.0) | 24.5–26.5  (51.6) | 7 |
| ∆*recD* SMR21915  (SMR21942) | 7.5  (7.0, 7.5) | 7.5  (7.5) | 21.7  (15.9–22.5) | 12 |
| ∆*rpoE* SMR21938  (SMR21911) | 7.5  (7.5, 8.0) | 7.5  (7.5) | 19.5  (8.2–12.8) | 10 |
| ∆*rnhA* SMR21940  (SMR21913) | 7.5  (7.5, 8.0) | 7.5  (7.5) | 26.8  (17.2–26.9) | 9 |
| ∆*pol IV* SMR21321  (SMR11640) | 8.0–18.5  (7.0–7.5) | 8.0  (7.0) | 1.3–11.0  (33.2–46.7) | 12 |
| ∆*pol* *II* ∆*pol IV* ∆*pol* *V*  SMR23982  (SMR23963) | 6.5–10.0  (6.5–7.5) | 8.2  (6.5) | 1.5–10.2  (29.2–37.6) | 14 |
| WT SMR15482  (SMR5223) | 4.0–12.8  (4.0–12.8) | 8.5  (8.5) | 1.0–56.4  (4.0–84.0) | 12 |
| WT thiourea (100mM)  SMR15482 *yiaG-yfp*  (SMR5223) | 8.5  (8.5) | 8.5  (8.5) | 47.6–74.1  (87.6–99.0) | ND |
| WT 2’,2’ bipyridyl (0.25mM)  SMR15482  (SMR5223) | 8.5  (ND) | 8.5  (ND) | 36.7–90.4  (ND) | ND |
| WT (vehicle bipyridyl)  SMR15482  (SMR5223) | 8.5  (ND) | 8.5  (ND) | 3.7–7.2  (ND) | ND |
| WT edaravone (1mM)  SMR15482  (SMR5223) | 8.5  (ND) | 8.5  (ND) | 18.7–48.1  (ND) | ND |
| WT (vehicle edaravone)  SMR15482  (SMR5223) | 8.5  (ND) | 8.5  (ND) | 3.8–6.6  (ND) | ND |
| WT GamGFP  Doxy 0 ng/mL  SMR14334 | 8.5  (ND) | 8.5  (ND) | 7.6–11  (ND) | ND |
| WT GamGFP  Doxy 10 ng/mL  SMR14334 | 8.5  (ND) | 8.5  (ND) | 7.6–30.5  (ND) | ND |
| WT GamGFP  Doxy 20 ng/mL  SMR14334 | 8.5  (ND) | 8.5  (ND) | 7.2–17  (ND) | ND |
| ∆*pol V*  SMR23930  (SMR23925) | 6.0–10.0  (7.0, 7.5) | 9.5  (7.5) | 3.9–16.6  (34.0–55.0) | 12 |
| ∆*rpos lexA*Ind^-^ SMR24004  (SMR24002) | 2.8–5.5  (2.8–4) | 5.5  (2.8) | 4.5–22.4  (22.3–35.8) | ND |
| *parC* gyrA** SMR24600 | 8.5  (ND) | 8.5  (ND) | 91.9–121.6  (ND) | ND |
| *parC* gyrA** thiourea (100mM) SMR24600 | 8.5  (ND) | 8.5  (ND) | 84.7–107.9  (ND) | ND |
| *ubiD* SMR24682  (SMR24676) | 8.5  (8.5) | 8.5  (8.5) | 5.5–8.6  (15.8–61.8) | ND |
| ∆*nuoC* SMR24678  (SMR24672) | 8.5  (8.5) | 8.5  (8.5) | 10.7–11.7  (24.6–40.0) | ND |
| ∆*cyoD* SMR24680  (SMR24674) | 8.5  (8.5) | 8.5  (8.5) | 8.4–9.9  (16.5–46.0) | ND |
| ∆*hfq* pVector SMR24452 | 6–8.5  (ND) | 6  (ND) | 8.8–89  (ND) | ND |
| ∆*hfq* pRpoS  SMR24453 | 6–8.5  (ND) | 6  (ND) | 8.8–89  (ND) | ND |
| *ubiD* pVector SMR24684 | 6–8.5  (ND) | 6  (ND) | 8.8–89  (ND) | ND |
| *ubiD* pRpoS  SMR24686 | 6–8.5  (ND) | 6  (ND) | 8.8–89  (ND) | ND |
| WT pVector  SMR24450 | 8.5  (ND) | 8.5  (ND) | 0.8–12.9  (ND) | ND |
| WT pRpoS  SMR24451 | 8.5  (ND) | 8.5  (ND) | 2.4–11.5  (ND) | ND |
| WT pVector  Thiourea (100mM)  SMR24450 | 8.5  (ND) | 8.5  (ND) | 3.0–37.5  (ND) | ND |
| WT pRpoS  Thiourea (100mM)  SMR24451 | 8.5  (ND) | 8.5  (ND) | 2.7–19.6  (ND) | ND |
| ∆*sulA* SMR21774  (SMR21772) | 11-20  (8.5-20) | 20  (17) | 0.5-29.8  (11.7-44.3) | ND |
| ∆*sulA* ∆*ruvC* SMR23985  (SMR23991) | 2-2.4  (1.35-2) | 2.2  (1.35) | 5.68-22.6  (2.4-29.7) | ND |

ND, not determined in this study
